# Proteome-wide quantification of inositol pyrophosphate-protein interactions

**DOI:** 10.1101/2024.07.31.606004

**Authors:** Annika Richter, Max Ruwolt, Sarah Lampe, Fan Liu, David Furkert, Dorothea Fiedler

**Affiliations:** Leibniz-Forschungsinstitut für Molekulare Pharmakologie (FMP), Robert-Rössle-Straße 10, 13125 Berlin, Germany; Institut für Chemie, Humboldt-Universität zu Berlin, Brook-Taylor-Straße 2, 12489 Berlin, Germany; Charité - Universitätsmedizin Berlin, Charitéplatz 1, 10117, Berlin, Germany

## Abstract

Inositol polyphosphates (InsPs) and inositol pyrophosphates (PP-InsPs) constitute a group of highly phosphorylated molecules that are involved in many cellular signaling processes. To characterize discrete signaling events of these structurally closely related molecules, a mass spectrometry approach was developed to derive apparent binding constants for these ligands on a proteome-wide scale. The method employed a series of chemically synthesized, biotinylated affinity reagents for inositol hexakisphosphate (InsP_6_), and the inositol pyrophosphates 1PP-InsP_5_, 5PP-InsP_5_ and 1,5(PP)_2_-InsP_4_ (also termed InsP_8_). Application of these affinity reagents at different concentrations, in combination with tandem mass tag (TMT) labeling, provided binding data for thousands of proteins from a mammalian cell lysate. Investigation of different enrichment conditions, where Mg^2+^ ions were either available or not, showcased a strong influence of Mg^2+^ on the protein binding capacities of PP-InsPs. Gene ontology analysis closely linked PP-InsP-interacting proteins to RNA processing in the nucleus and nucleolus. Subsequent data analysis enabled a targeted search for protein pyrophosphorylation among PP-InsP interactors, and identified four new pyrophosphorylated proteins. The data presented here constitute a valuable resource for the community, and application of the method reported here to other biological contexts will enable the exploration of PP-InsP dependent signaling pathways across species.

## INTRODUCTION

The soluble inositol polyphosphates (InsPs) are *myo*-inositol based signaling molecules that occur ubiquitously in eukaryotes. A classic example of these messengers is inositol-1,4,5-trisphosphate (InsP_3_), which can be cleaved from the cell membrane upon phospholipase C activation, and trigger calcium release from the endoplasmatic reticulum^1^. Phosphorylation of InsP_3_ can yield fully phosphorylated inositol hexakisphosphate (InsP_6_), which has been reported to act as a signaling molecule and structural cofactor in mammals, and serves as phosphate storage in plants^2–6^. The subsequent action of inositol hexakisphosphate kinases (IP6Ks), which transfer a phosphoryl group onto the phosphate at the 5-position of the *myo*-inositol ring, and diphosphoinositol pentakisphosphate kinases (PPIP5Ks), which phosphorylate the phosphate at the 1-position, leads to the generation of the inositol pyrophosphates 5-diphosphoinositol pentakisphosphate (5PP-InsP_5_), 1-diphosphoinositol-pentakisphosphate (1PP-InsP_5_), and bis-1,5-diphosphoinositol tetrakisphosphate (1,5(PP)_2_-InsP_4_) (**Figure 1**)^7–13^.

**Figure 1.**
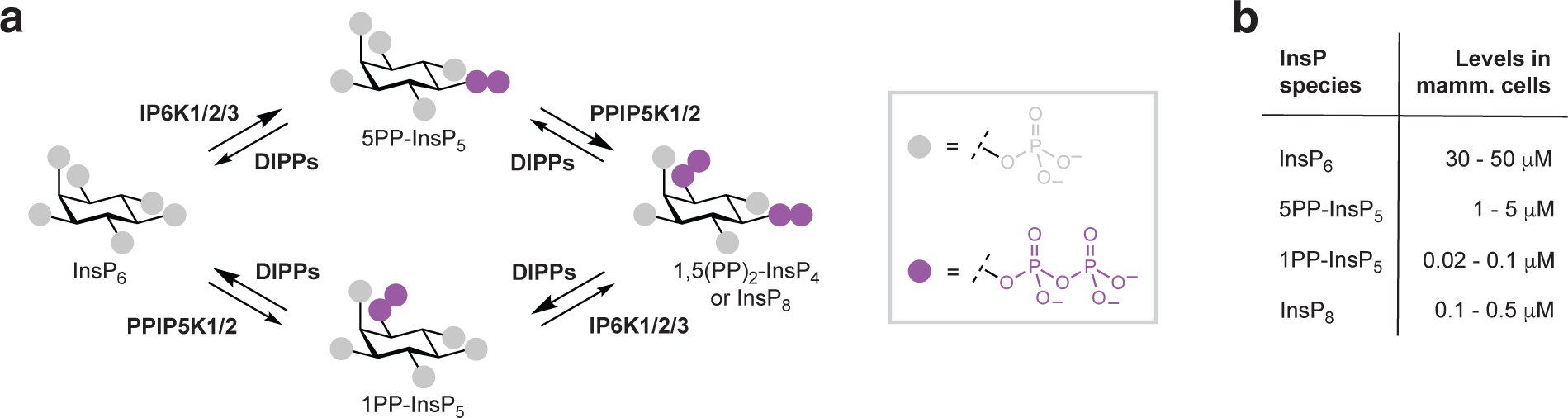
Inositol pyrophosphate biosynthesis and estimated levels in mammalian cells. **a**: Pathway for the synthesis and turn-over of inositol pyrophosphates. InsP_6_ (inositol hexakisphosphate) is converted to 5PP-InsP_5_ (5-diphosphoinositol pentakisphosphate) by the enzymes IP6K1/2/3 (inositol hexakisphosphate kinase 1, 2, or 3) or to 1PP-InsP_5_ (1-diphosphoinositol-pentakisphosphate) by enzymes PPIP5K1/2 (diphosphoinositol pentakisphosphate kinase 1 or 2). Subsequently, 5PP-InsP_5_ can be converted to 1,5(PP)_2_-InsP_4_ (bis-1,5-diphosphoinositol tetrakisphosphate) by IP6K1/2/3 and 1PP-InsP_5_ can be converted to 1,5(PP)_2_-InsP_4_ by PPIP5K1/2. The dephosphorylation reactions are catalyzed by DIPPs (DIPP1, DIPP2, DIPP3α and DIPP3β; diphosphoinositol polyphosphate phosphohydrolase 1, 2, or 3). **b**: Concentration ranges of (PP)-InsPs detected in different human cell lines^14–17^.

A putative role in cellular signaling has been attributed to the PP-InsP for a long time, as these molecules are rapidly turned over in cells (up to ten times per hour)^18,19^. Current analytical methods are not capable of resolving the detection and quantification of InsP/PP-InsPs with subcellular resolution, therefore many questions regarding local turnover and availability for signaling purposes have remained unanswered. At the cellular level, InsP/PP-InsP levels have been reported to range from nanomolar to micromolar concentrations, suggesting significant differences in their availability for signaling processes (**Figure 1b**)^14–17^. Among the PP-InsPs, 5PP-InsP_5_ is the most abundant representative in most cell lines, with concentrations typically ranging between 1 µM and 5 µM.

Over the past years, the signaling functions of PP-InsPs have mainly been investigated using genetic methods, as well as pharmacologic tools targeting IP6Ks^10,20^. These kinases – and in several cases by proxy the molecule 5PP-InsP_5_ – have been associated with insulin secretion^21,22^, focal adhesion dynamics^23,24^, and apoptosis^25^. In these examples, 5PP-InsP_5_ accessed different modes of action for signal transduction, including competition for phospholipid binding domains. In addition, 5PP-InsP_5_ can also transmit signals by transferring the β-phosphoryl group onto pre-phosphorylated proteins in a process termed pyrophosphorylation^26–29^. This unusual protein modification was demonstrated, for example, to regulate protein localization, protein degradation, and glycolysis^30–32^. Compared to the functions of 5PP-InsP_5_, relatively little is known about the closely related messengers 1PP-InsP_5_ and 1,5(PP)_2_-InsP_4_. Nevertheless, analyses of cell lines lacking PPIP5Ks, and consequently 1,5(PP)_2_-InsP_4_, have sparked recent interest because these cell lines exhibited a growth-inhibited phenotype and a hypermetabolic state, indicating a potential application in tumor therapy^33,34^.

Deciphering the concrete signaling functions of individual PP-InsPs remains challenging. Genetic or pharmacologic perturbation of IP6Ks not only reduces 5PP-InsP_5_ levels but simultaneously also diminishes 1,5(PP)_2_-InsP_4_ levels. Deletion of PPIP5Ks depletes 1,5(PP)_2_-InsP_4_, but concomitantly increases the cellular amounts of 5PP-InsP_5_^34^. Therefore, phenotypic observations must be complemented by biochemical and/or biophysical data to assign specific functions to a defined PP-InsP molecule. For example, a recent study demonstrated that the xenotropic and polytropic retrovirus receptor 1 (XPR1) - a protein that binds to InsP_6_, 5PP-InsP_5,_ and 1,5(PP)_2_-InsP_4_ - is regulated predominantly by the latter^35^. The *in vitro* binding affinities did not differ drastically, but were found to be highest for 1,5(PP)_2_-InsP_4_^36,37^. Interestingly, NMR experiments revealed a differential conformational plasticity of the different protein-ligand complexes, where 1,5(PP)_2_- InsP_4_ engages in the most dynamic interaction mode^38^. These findings highlight the need for a systematic determination of InsP/PP-InsP protein binding affinities, to elucidate which proteins are potentially regulated by which member of the PP-InsP family. Thus far, only purified proteins have been used to study PP-InsP-protein binding affinities *in vitro*, using methods such as isothermal titration calorimetry, surface plasmon resonance spectroscopy, and microscale thermophoresis^24,38,39^. These measurements can provide precise information, but cannot be used for large-scale analyses. Recently, a mass spectrometric method for the global determination of apparent binding affinities to immobilized small molecule ligands was reported^40–42^. Based on previous experiments where InsP_6_ and a non-hydrolyzable bisphosphonate analog of 5PP-InsP_5_ were used for affinity enrichment, we wanted to implement a mass spectrometric approach to quantify the proteome-wide interactions of InsP_6_, 5PP-InsP_5_, 1PP-InsP_5_ and 1,5(PP)_2_-InsP_4_.

Here, we report the synthesis of affinity reagents for 5PP-InsP_5_, 1PP-InsP_5_ and 1,5(PP)_2_-InsP_4_, in which the pyrophosphate groups have been replaced by non-hydrolyzable bisphosphonate (PCP) moieties. These reagents were applied over a wide concentration range, and, in combination with tandem mass tag (TMT) labeling, allowed us to extract binding data for these ligands under different conditions on a proteome-wide scale. The presence of Mg^2+^ ions had an influence on the enriched proteins and this effect became more pronounced as the number of pyrophosphate groups increased. Over 700 binding proteins were characterized for 5PP-InsP_5_ and 1,5(PP)_2_- InsP_4_, many of which are involved in RNA processing and ribosome biogenesis. Furthermore, based on the affinity enrichment data, previously unknown pyrophosphorylation targets could be identified. In the future, the reported affinity reagents, in combination with the quantitative MS method, can be applied in different biological settings, to facilitate the elucidation of PP-InsP signaling mechanisms.

## RESULTS

### Design and synthesis of biotinylated inositol pyrophosphate analogs

While 5PP-InsP_5_ appears to be the most abundant PP-InsP in many mammalian cell lines,^7,15^ a few reports have invoked a signaling role for the closely related molecule 1PP-InsP_5_^43^. In addition, several recent studies have substantiated a unique signaling function for 1,5(PP)_2_-InsP_4_^34,35,44^. Building on our previously developed PCP-InsP affinity reagents^45,46^, we wanted to expand the set of reagents to include probes for 1PP-InsP_5_ and 1,5(PP)_2_-InsP_4_. Because the modification of a phosphoryl group with a linker can influence the protein interactions of PP-InsPs, phosphate groups immediately adjacent to the pyrophosphate moiety/moieties were not considered for derivatization. Therefore, the emerging target structures were 1PCP-InsP_5_, derivatized at the 3- or 5-position, and 1,5(PCP)_2_-InsP_4_, derivatized a the 3-position (**Figure 2a**). Finally, we sought to incorporate a biotin group in place of the primary amine for attachment to the resin, to facilitate the immobilization step^47^.

**Figure 2.**
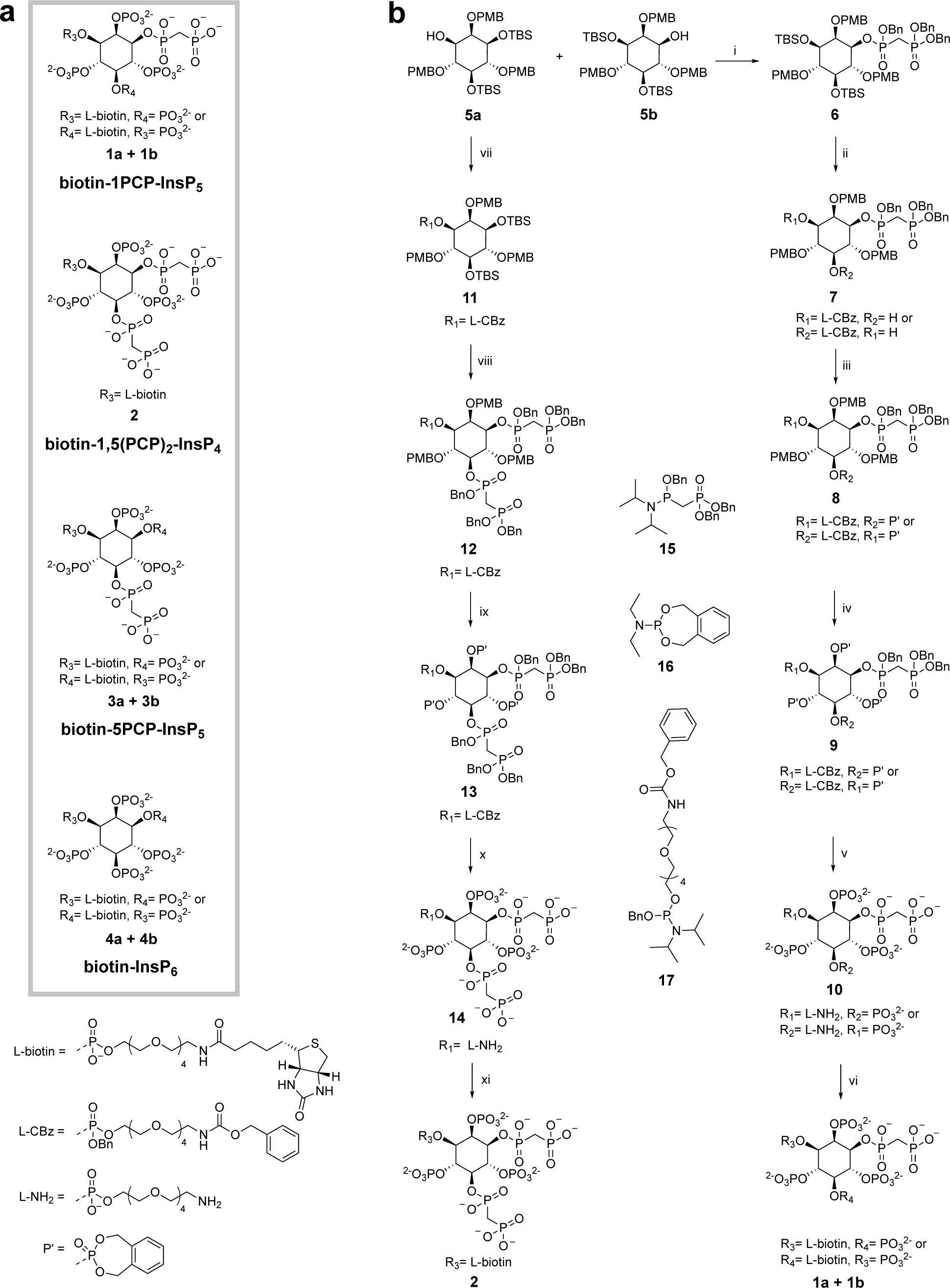
Synthetic route for biotin-(PCP)-InsP compounds. **a**: Overview of biotin-(PCP)-InsP compounds. **b**: Synthesis of biotin-1PCP-InsP_5_ (**1a+ 1b**) and biotin-1,5(PCP)_2_-InsP_4_ (**2**): (i) **15**, 5-phenyl-1H-tetrazole, CH_2_Cl_2_ then *m*CPBA, 49% (ii) TBAF, THF, 75%. Afterwards **17**, 5-phenyl-1H-tetrazole, CH_2_Cl_2_ then *m*CPBA, directly used as crude (iii) **16**, tetrazole, ACN, then *m*CPBA, 56% over four steps, (iv) TFA in CH_2_Cl_2_ then **16**, tetrazole, ACN, then *m*CPBA, 17% (v) Pd/C, *t*BuOH/H2O, 83% (vi) NHS-biotin, KH_2_PO_4_, H_2_O, 81% (vii) **17**, tetrazole, ACN then *m*CPBA, 58% (viii) TBAF, DMF, 97%. Afterwards **15**, tetrazole, ACN, then *m*CPBA, 57% (ix) TFA in CH_2_Cl_2_ then **18**, tetrazole, ACN, then *m*CPBA, 71% (x) Pd/C, *t*BuOH/H_2_O, 98% (xi) NHS-biotin, KH_2_PO_4_, H_2_O, 58%.

Because the affinity reagents for 1PCP-InsP_5_ and 1,5(PCP)_2_-InsP_4_ are asymmetrical, a synthetic strategy to obtain enantiopure material was needed. Starting from *myo*-inositol, an enantiomeric mixture of tert-butyl-dimethylsilyl (TBS)- and para-methoxybenzyl (PMB)-protected compounds **5a** and **5b** could be prepared in six steps (**Figure 2b**)^48,49^. The two enantiomers were separated on a gram scale using a chiral aromatic stationary phase column (**Supplementary Figure S1a**) The two enantiomers were assigned through phosphitylation of the free hydroxyl group, followed by global deprotection, yielding the enantiomeric 1-InsP_1_ and 3-InsP_1._ Finally, optical rotations of these compounds were then measured to complete the assignment of **5a** and **5b** (**Supplementary Figure S1b**).

Enantiomer **5b** was then further used for the synthesis of biotin-1PCP-InsP_5_ (**Figure 2b**). The free hydroxyl group at the 1-position was reacted with phosphoramidite **15** followed by oxidation to yield compound **6**. Deprotection of the TBS groups and subsequent addition of one equivalent of phosphoramidite **17**, followed by oxidation, yielded compound mixture **7**, in which the linker was attached either at the 3- or 5-position. To increase the solubility in organic solvents after deprotection, the remaining hydroxyl group was modified with phosphoramidite **16** to provide compound mixture **8**. Subsequently, the PMB groups were removed and the resulting triol was phosphitylated and oxidized to furnish fully protected compounds **9**. Final deprotection *via* palladium-catalyzed hydrogenolysis yielded linker-derivatized 1PCP-InsP_5_ (**10**), modified at either the 3- or the 5-position. The primary amine in the linker was then coupled to NHS-biotin to provide the final product mixture of biotin-1PCP-InsP_5_ (**1a + 1b**).

The chemical structures of compound mixture **1a + 1b** were additionally investigated by 2D ^31^P- HMBC, as well as ^1^H-^13^C-DEPT-CLIP-COSY NMR experiments (**Supplementary Figure S2**). The spectra confirmed the correct attachment of the linker and PCP-groups at the desired positions and also revealed a diastereomeric mixture of 80% **1a**, where the linker is attached at the 3- position, and 20% **1b**, with a linker incorporated at the 5-position. Likely, the linker attachment proceeds faster at the stereochemically less hindered 3-position, due to the neighboring axial 2- OH group.

Enantiomer **5a** was the central starting material for the synthesis of the 1,5(PCP)_2_-InsP_4_ affinity reagent (**Figure 2b**). The free hydroxyl group at the 3-position was derivatized with the linker using phosphoramidite **17**, and oxidized to yield intermediate **11**. Following the removal of the TBS protecting groups, two PCP moieties were appended at the 1- and 5- positions utilizing phosphoramidite **15**, and subsequent oxidation yielded **12**.

A 55% yield was obtained over two steps, highlighting the good compatibility of phosphoramidite **15** with challenging, sterically hindered substrates. From here on, the same synthetic sequence was applied as above, ultimately yielding enantiopure biotin-1,5(PCP)_2_-InsP_4_ (**2**). The structure was corroborated by 2D-NMR spectroscopy, validating the linker attachment at the 3-position and the PCP groups at the 1- and 5-positions (**Supplementary Figure S3**).

To complete the series of biotinylated PCP affinity reagents, biotin-5PCP-InsP_5,_ and biotin-InsP_6_ probes, with two alternative linker attachment sites, were also synthesized (**Supplementary Figure S4, Figure 2a**)^39,46^. All affinity reagents were quantified using ^1^H-NMR spectroscopy and an internal standard (3-(trimethylsilyl)-2,2,3,3-propanoate-d_4_), and prepared as 1 mM stock solutions for subsequent experiments. The described synthesis of enantiomer **5a** also provided a useful starting material to access soluble 1PCP-InsP_5_ in overall good yields (**Supplementary Figure S5**). Combined with the previously described syntheses of 1,5(PCP)_2_-InsP_4_ and 5PCP-InsP_5_, all non-hydrolyzable analogs were obtained in good quantities^50^.

In sum, a synthetic strategy relying on enantiomeric separation was developed to provide 1PCP- InsP_5_ and 1,5(PCP)_2_-InsP_4_ affinity probes. This series of affinity probes can now be used for global analysis of PP-InsP-protein interactions.

### Validation and dose-dependent binding of PCP-InsP affinity reagents

With this set of affinity reagents in hand, we next investigated their ability to retain known binding partners. The biotinylated probes were immobilized on streptavidin-coated sepharose beads, and subsequently human diphosphoinositol polyphosphate phosphohydrolase 1 (DIPP1), the SPX domain of XPR1 (XPR1^SPX^), and the C2B domain of synaptotagmin 1 (SYT1^C2B^) were applied to the different beads. Unmodified streptavidin beads (Ctrl) were handled in parallel as control experiments. Following incubation and washing, the bound proteins were eluted with an excess of the corresponding (PCP)-InsP ligand. For all three proteins, strong retention by the four different affinity reagents, but not the unmodified beads, was observed (**Figure 3a**).

**Figure 3.**
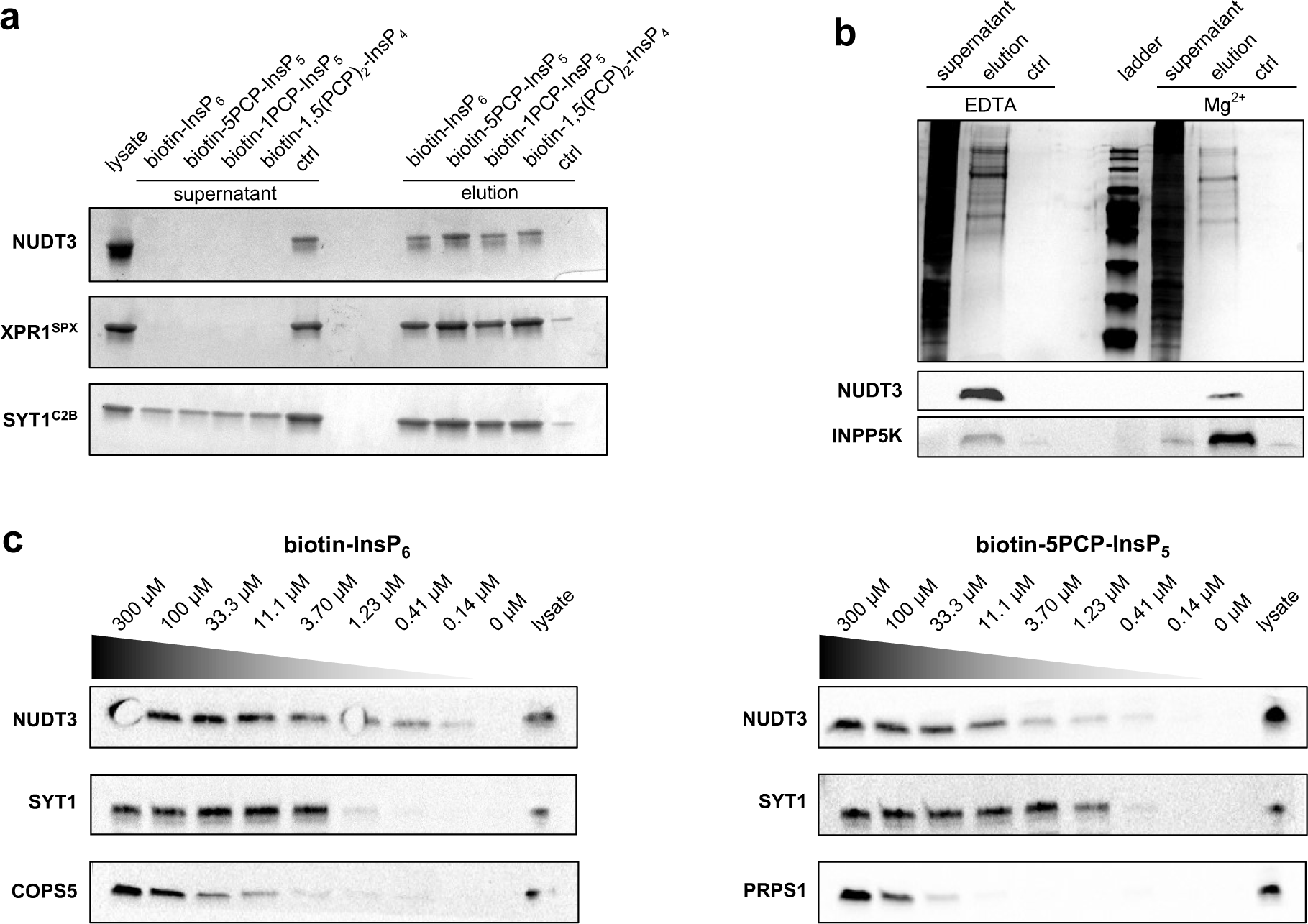
Validation of affinity reagents and concentration-dependent affinity enrichment. **a**: Biochemical validation of affinity reagents biotin-1PCP-InsP_5_ (**1**), biotin-1,5(PCP)_2_-InsP_4_ (**2**), biotin-5PCP-InsP_5_ (**3**), and biotin-InsP_6_ (**4**), (20 nmol each). Each affinity reagent was incubated with 5 nmol of DIPP1 (GST-DIPP1), 5 nmol of SPX domain of XPR1 (His_6_ -MBP-XPR1^SPX^), or 5 nmol of the C2B domain of SYT1 (MBP-SYT1^C2B^). After 30 min incubation, the supernatant was collected and the beads were washed three times. The bound fraction was eluted with an excess of InsP_6_, 5PCP-InsP_5_, 1PCP-InsP_5,_ and 1,5(PCP)_2_-InsP_4_ (5 mM each). Fractions were separated by SDS-page and visualized by silver stain. Ctrl denotes streptavidin coated sepharose beads. **b**: HEK293T cells were lysed with two different buffers (common components: 25 mM HEPES, pH 7.4, 150 mM NaCl, 1 % NP-40) containing either 1 mM EDTA or 1 mM MgCl_2_. The resulting lysates were incubated with immobilized biotin-5PCP-InsP_5_ and supernatants as well as eluates were separated by SDS-page and visualized by silver stain. All fractions were also analyzed by western blot using antibodies targeting DIPP1 and INPP5K. **c**: Concentration-dependent affinity enrichment using immobilized biotin-InsP_6_ and biotin-5PCP-InsP_5_. HEK293T cell lysate (1 mg/mL) was incubated with a three-fold dilution series of affinity reagents for 1 h, under Mg^2+^-depleted conditions. Bound fractions were washed and eluted with an excess of InsP_6_ and 5PCP-InsP_5_. Resulting fractions were analyzed by western blot using antibodies targeting DIPP1, SYT1, COPS5, and PRPS1.

Next, we wanted to evaluate how the affinity reagents retained proteins from cell lysates. Given the high negative charge density of PP-InsPs, these molecules interact strongly with di- and trivalent metal cations^49,51,52^. Since the formation of these metal complexes likely influences the binding preferences towards different proteins, HEK293T cell lysates were prepared with 1 mM EDTA to deplete di- and trivalent cations and with 1 mM MgCl_2_ (Mg^2+^ ions were chosen since they are the most abundant divalent metal ions in cells)^53^. The lysates (1 mg/mL) were incubated with immobilized biotin-5PCP-InsP_5_. Following a washing step, the bound fractions were eluted with an excess of 5PCP-InsP_5_ (**Figure 3b**). The banding pattern of the eluted proteins displayed differences between the two conditions. Additionally, western blot analysis of two known binding proteins, DIPP1 and inositol polyphosphate 5-phosphatase K (INPP5K),^46,54^ showed enrichment under both conditions, albeit with varying retention. The qualitative analysis of the elution profiles illustrates that the formation of PP-InsP-protein interactions is notably influenced by the presence of Mg^2+^ ions. Therefore, investigation of both conditions can provide insights into how PP-InsP– protein interactions respond to the presence of divalent/coordinating metal cations.

Because the amount of biotinylated PCP-InsP probes can be readily altered during immobilization, we next investigated the retention of proteins from cell lysates at different probe concentrations. A three-fold dilution series (between 300 µM and 140 nM) of biotin-InsP_6_ and biotin-5PCP-InsP_5_ was prepared and immobilized, and subsequently incubated with HEK293T cell lysate (under metal-depleted conditions). The eluates were analyzed by western blot for known binding proteins (DIPP1, SYT1, and COP9 signalosome complex subunit 5 (COPS5) for biotin-InsP_6_; and DIPP1, SYT1, and ribose-phosphate pyrophosphokinase 1 (PRPS1) for biotin-5PCP-InsP_5_)^46,55^. While a concentration-dependent enrichment was observed for all analyzed proteins, the apparent affinities towards InsP_6_ and 5PCP-InsP_5_ varied (**Figure 3c** and **Supplementary Figure S6**). For example, DIPP1 was retained more strongly by biotin-InsP_6_ compared to COPS5 and SYT1, as evidenced by the elution profile (**Figure 3c**). For biotin-5PCP-InsP_5_ on the other hand, SYT1 exhibited a higher affinity towards the ligand, compared to DIPP1 and PRPS1. These experiments demonstrated that a qualitative comparison of the binding affinities of individual proteins towards biotin affinity probes is possible, using standard western blot analysis. We conclude that the biotinylated probes can be applied to reveal the dose-dependent binding of various proteins. Analysis of the eluted proteins by mass spectrometry should, in principle, provide binding affinities on a proteome-wide level.

### A proteomics workflow for determination of apparent binding constants

Inspired by recent progress in the identification of polyADPr binding proteins by Kliza *et al.*^41^, we sought to combine the biotin affinity probes with Tandem Mass Tag (TMT) isobaric labeling to determine binding affinities on a proteome-wide scale (**Figure 4a**)^40–42^. Cells were lysed using two different lysis buffers (containing either 1 mM EDTA, or alternatively containing 1 mM Mg^2+^), and the lysates were separated into nuclear and cytosolic fractions to provide deeper proteome coverage (**Supplementary Figure S7**). Both fractions were incubated with a serial dilution (100 µM – 5 nM) of the immobilized probes biotin-InsP_6_, biotin-5PCP-InsP_5_, biotin-1PCP-InsP_5_ and biotin-1,5(PCP)_2_-InsP_4_ (**Figure 4a**). Following quick washing steps, the retained proteins were eluted with an excess of the corresponding free ligand. The samples were digested with trypsin, followed by 11-plex TMT isobaric labeling, subsequent sample pooling, and LC-MS/MS analysis (**SI Figure 8**). Each enrichment was conducted in triplicate, and only proteins where at least one replicate was fully quantified were included. Hill-like curves were generated, and for curves with R^2^ < 0.9 the corresponding K_D_^app^ values were calculated (**Supplementary Tables 1-4**). Exemplary curves are shown in **Supplementary Figure S9** and encompass an interaction of InsP_6_ with C2 domain-containing protein 5 (C2CD5), in which the C2-domain likely binds to InsP_6_. Furthermore, we observed tight interactions between biotin-5PCP-InsP_5_ and beta-arrestin-1 (ARRB1) as had been reported before^23,56^, as well as strong binding between biotin-1,5(PCP)_2_-InsP_4_ and DNA-directed RNA polymerase II subunit RPB1 (POLR2A). In addition to these interactions, many known binding partners, including the (PP)-InsP metabolizing enzymes IP6K1, IP6K2, PPIP5K2, DIPP1, and DIPP2 could be identified. As anticipated, depending on the protein interaction partner the distinct ligands displayed a difference in K_D_^app^ values. For example, in the presence of Mg^2+^ ions PPIP5K2 bound tightly to 5PCP-InsP_5_ and 1,5(PCP)_2_-InsP_4_, while the K_D_^app^ value was increased for 1PP-InsP_5_ and even more so for InsP_6_ (**Figure 4b**). The motor protein KIF14 showed the highest affinity towards 5PP-InsP_5_. And phosphomevalonate kinase (PMVK) exhibited a very strong preference for binding to 1,5(PCP)_2_-InsP_4_ (K_D_^app^<1 µM); the apparent binding constants for InsP_6_ and 5PCP-InsP_5_ were more than 40-fold higher.

**Figure 4.**
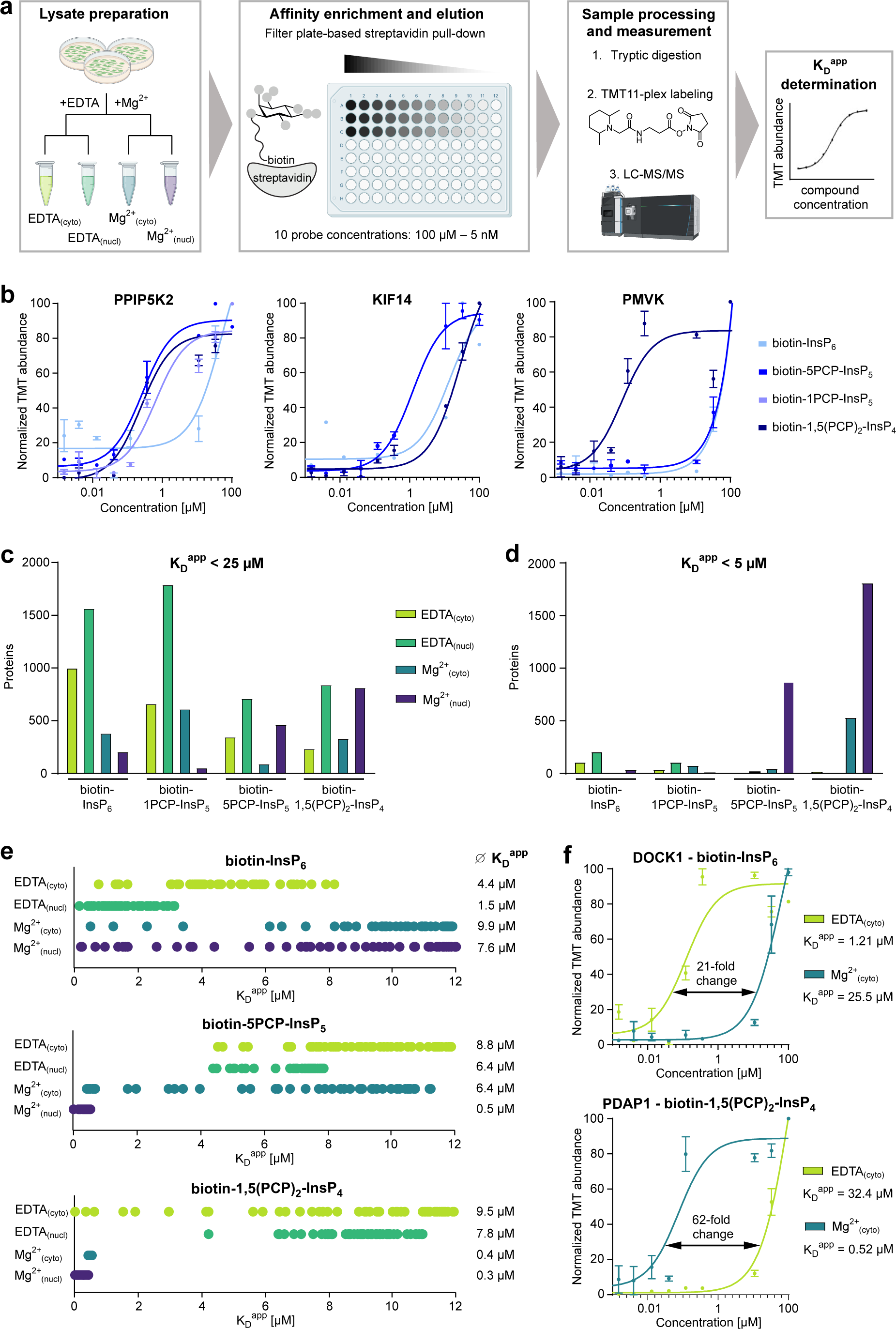
Proteome-wide quantification of (PP)-InsP binding proteins from cell lysates. **a**: Schematic overview of the work-flow for quantitative affinity enrichment. Four different lysate preparations were applied to the four affinity reagents (used at concentrations between 5 nM and 100 µM). Following elutions, the samples are subjected to TMT11-plex labeling and LC-MS/MS analysis. The binding curves were used to calulate K_D_^app^ values. The different colors indicate the lysate buffer composition/cellular fraction: light green cytosolic fraction of EDTA-containing buffer; dark green: nuclear fraction of EDTA-containing buffer; blue: cytosolic fraction of Mg^2+^-containing buffer; purple: fraction of Mg^2+^-containing buffer. **b**: Binding curves for the (PP)-InsP binding proteins PPIP5K2, kinesin-like protein KIF14 (KIF14) and phosphomevalonate kinase (PMVK). All values are obtained from the Mg^2+^ cytosolic dataset and the affinity probe used is shown under the curves. Every data point represents the mean of at least two replicates and the error bars depict the SEM. **c**, **d**: Number of all fully quantified proteins with R^2^ < 0.90 for the different affinity reagents in the four lysate preparations. The cut-off for the K_D_^app^ values is either set to K_D_^app^ < 25 µM (c), or K_D_^app^ < 5 µM (d). **e**: For the four lysate preparations, the top 50 proteins with the lowest K_D_^app^ values are displayed for the affinity probes biotin-InsP_6_, biotin -5PCP-InsP_5,_ and biotin -1,5(PCP)_2_-InsP_4_. Each dot represents the K_D_^app^ value for a specific protein. The average K_D_^app^ values (across the top 50 hits) are listed on the right side for the respective datasets. **f**: Binding curves for the InsP_6_ and 1,5(PCP)_2_-InsP_4_ interacting partners dedicator of cytokinesis protein 1 (DOCK1) and PDGFA-associated protein 1 (PDAP1). Each data point represents the mean of three replicates, and the error bars depict the SEM. The fold-change between the enrichment conditions (plus or minus Mg^2+^) is shown in the graphs.

### Global analysis shows general and PP-InsP specific trends

As the examples above illustrate, the proteomic analysis revealed a very wide distribution of K_D_^app^ values across the different probes, fractions, and enrichment conditions. Therefore, cut-off values were defined, based on the reported intracellular levels of InsPs/PP-InsPs. In most human cell lines, InsP_6_ is quite abundant with a concentration range of 10 – 50 µM^7^. PP-InsPs, by contrast, are less abundant and their levels have been estimated to be 1–5 µM for 5PP-InsP_5_, 0.1 – 0.5 µM for 1,5(PP)_2_-InsP_4_, and 0.02 – 0.1 µM for 1PP-InsP_5_ (**Figure 1b**)^44,57^.

When the cut-off is set at K_D_^app^ < 25 µM, a total number of ca. 10,000 interacting proteins across all 16 datasets were identified (**Figure 4c**). This cut-off, however, is likely only relevant for InsP_6_-binding proteins, which is why a lower cut-off K_D_^app^ < 5 µM was implemented for the less abundant PP-InsPs. With this lower cut-off, a very different picture emerged: A large number of interactors was identified for biotin-5PCP-InsP_5_ and biotin-1,5(PCP)_2_-InsP_4_ (with Mg^2+^ present), and only very few proteins were enriched with the biotin-1PCP-InsP_5_ probe (**Figure 4d**). In fact, the biotin-1PCP-InsP_5_ affinity reagent enriched only 85 proteins with a K_D_^app^ < 5 µM, and none were observed with a K_D_^app^ < 1 µM. In many aspects, the 1PCP-InsP_5_ interactome displayed high similarity to the enrichment with biotin-InsP_6_. When the interactomes of 1PCP-InsP_5_ and InsP_6_ were compared under Mg^2+^-free conditions, more than 60% of enriched proteins were identified with both ligands (**Supplementary Figure S10**). Considering the low physiological concentrations of 1PP-InsP_5_, however, our data calls into question whether 1PP-InsP_5_ serves as an actual signaling molecule in mammalian cells, given the comparatively weak protein interactions of this ligand.

The influence of coordinating divalent cations on PP-InsP protein interactions is well apparent in the proteomics data. Interestingly, a general weakening of InsP_6_-protein interactions was observed when Mg^2+^-ions were added during the affinity enrichment, while biotin-1,5(PCP)_2_-InsP_4_ displayed the opposite behavior. When the average K_D_^app^ values of the top 50 proteins in each dataset were examined, biotin-InsP_6_ bound to proteins with higher affinities when coordinating di- and trivalent ions were depleted (**Figure 4e)**. In contrast, the interactions of biotin-5PCP-InsP_5_ shifted towards higher affinities with Mg^2+^-ions present. For biotin-1,5(PCP)_2_-InsP_4_ the effect became even more pronounced and all K_D_^app^ values were below 1 µM. These trends are exemplified by the interaction of biotin-InsP_6_ with dedicator of cytokinesis protein 1 (DOCK1), where the interaction is weakened 21-fold by the presence of Mg^2+^-ions (**Figure 4f**). The opposite was observed for binding of 28 kDa heat- and acid-stable phosphoprotein (PDAP1) to biotin-1,5(PCP)_2_-InsP_4_, where a 64-fold increase in affinity occured upon Mg^2+^ addition. Interestingly, the Mg-coordination of PP-InsPs appears to impose some selectivity of protein target recognition in general. While many identical proteins (from the nuclear fraction) were quantified in the presence or absence of Mg^2+^ using the InsP_6_-reagent, much less overlap between these two conditions was observed for the interactors of 5PP-InsP_5_ and 1,5(PCP)_2_-InsP_4_ (**Supplementary Figure S10**). Intrigued by the overall high number of proteins that were enriched by 5PCP-InsP_5_ and 1,5(PCP)_2_-InsP_4_ in the magnesium-containing nuclear fraction, we decided to further analyze these datasets.

### Strong Mg^2+^ dependent interactions of 5PP-InsP_5_ and 1,5(PP)_2_-InsP_4_ in the nucleus

For many protein interactions of 5PP-InsP_5_ and 1,5(PP)_2_-InsP_4_ (in the presence of Mg^2+^ ions) the K_D_^app^ values were below 1 µM. To rule out that the affinity enrichment is biased towards highly abundant proteins, the iBAQ (intensity-based absolute quantification) values of the nuclear Mg^2+^ lysate were analyzed and the 20 proteins with the lowest K_D_^app^ values towards 1,5(PP)_2_-InsP_4_ were plotted against the iBAQ values of a HEK293T lysate (**Figure 5a**)^26,58^. The enriched proteins were relatively evenly distributed over four orders of magnitude and included highly abundant proteins, such as inosine triphosphate pyrophosphatase (ITPA), and proteins of low abundance like RNA-binding protein 20 (RMB20).

**Figure 5.**
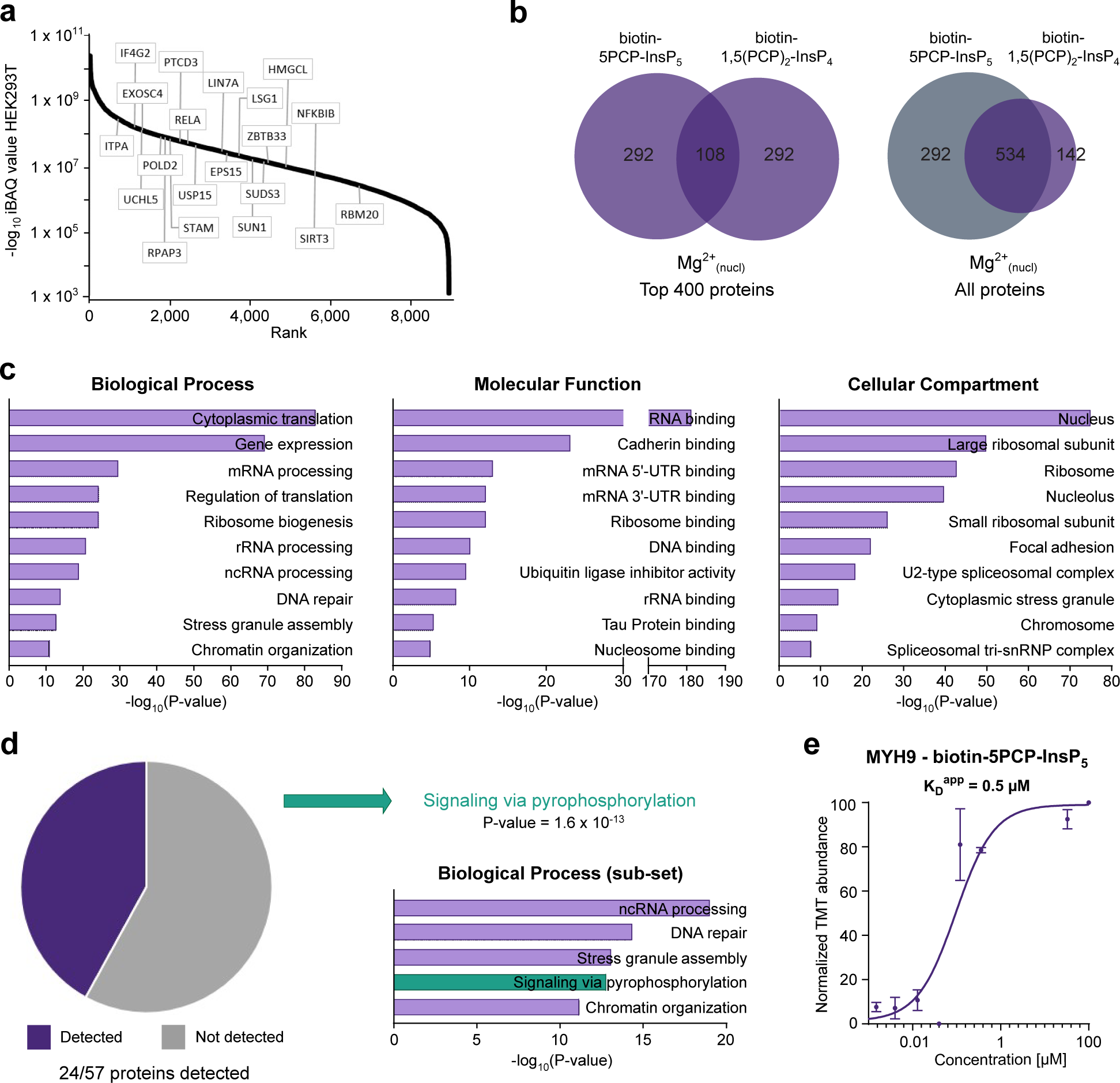
Associated functions for nuclear proteins interacting with 5PCP-InsP_5_ and 1,5(PCP)_2_-InsP_4_. **a**: iBAQ values of a whole HEK293T proteome were ranked and plotted against their –log 10 iBAQ values. The top 20 proteins displaying the lowest K_D_^app^s that were enriched by biotin-1,5(PCP)_2_-InsP_4_ from the Mg^2+^_nucl_ lysate are highlighted in the graph. **b**: Venn diagrams comparing the number of proteins enriched from Mg^2+^_nucl_ lysate with either biotin-5PCP-InsP_5_ or biotin-1,5(PCP)_2_-InsP_4_. Either the top 400 proteins, sorted by K_D_^app^ values, were considered (left), or all proteins for which a binding curve could be determined were included (right). **c**: Gene ontology (GO) analysis of for all 863 proteins enriched by biotin-5PCP-InsP_5_ and biotin-1,5(PCP)_2_-InsP_4_ (K_D_^app^ < 5 µM, Mg^2+^_nucl_).^59^ **d**: Pie chart of all proteins enriched by biotin-5PCP-InsP_5_ and/or biotin-1,5(PCP)_2_-InsP_4_ in the Mg^2+^_nucl_ lysate with a K_D_^app^_nucl_ < 5 µM that were also identified as pyrophosphoproteins in a previous pyrophosphoproteomic study of HEK293T cell lysates ^26^. A new GO term “signaling *via* pyrophosphorylation” was calculated and added to the biological processes to illustrate overrepresentation. **e**: Binding curve for the 5PCP-InsP_5_ binding protein myosin 9 (MYH9) from Mg^2+^_nucl_ lysate. Every data point represents the mean of three replicates and the error bars depict the SEM.

Given the close structural similarity between 5PCP-InsP_5_ and 1,5(PCP)_2_-InsP_4_, we wondered what the overlap between the proteins targeted by these two ligands was (**Figure 5b**). When comparing the top 400 proteins, sorted for K_D_^app^ values, only around 27% of the proteins were enriched in both datasets. However, when all proteins were included in the comparison (without a K_D_^app^ cut-off), more than 80% of the proteins enriched by biotin-5PCP-InsP_5_ were bound by biotin-1,5(PCP)_2_-InsP_4_ as well. This indicates that in most cases proteins can interact with both PP-InsPs, but the differentiation appears to be driven by the affinity towards the respective PP-InsPs.

### Nuclear PP-InsP interaction partners are associated with RNA processing and ribosome biogenesis

For a general view of putative biological functions of 5PP-InsP_5_ and 1,5(PP)_2_-InsP_4_ in the nucleus, the gene ontology (GO) terms for all 863 proteins enriched by biotin-5PCP-InsP_5_ and biotin-1,5(PCP)_2_-InsP_4_ (K_D_^app^ < 5 µM, Mg^2+^, **Figure 5c)** were determined (**Supplementary Table 5**).^59^ While we expected an enrichment of nuclear proteins, there appeared to be preferences between subnuclear compartments, including a localization to the nucleolus, and the spliceosomal complex. Additionally, components of the ribosome were overrepresented among the protein interactors, which aligns well with the biological processes, where many proteins involved in ribosome biogenesis, rRNA processing and cytoplasmic translation were identified. Also, the molecular functions aligned with these observations, and included ribosome binding and rRNA binding. Overall, the data implies a functional role of 5PCP-InsP_5_ and 1,5(PCP)_2_-InsP_4_ in ribosome biogenesis and transcriptional processes. Supporting these observations, a role for 5PP-InsP_5_ in regulating the transcription of rDNA was recently reported, putatively *via* pyrophosphorylation of several factors that localize to the nucleolus^45,46^.

In general, the nucleolus stands out as an interesting compartment where regulation by 5PP-InsP_5_ can take place^26,28,60,61^, and therefore we were curious if any of the recently identified pyrophosphoproteins were enriched by biotin-5PCP-InsP_5_ and/or biotin-1,5(PCP)_2_-InsP_4_ in the Mg^2+^ containing nuclear sample. Indeed, 24 out of the 57 known pyrophosphoproteins (in HEK293T) were enriched by biotin-5PCP-InsP_5_ and/or biotin-1,5(PCP)_2_-InsP_4_ **(Figure 5d**).^26^ Among the interactors were multiply pyrophosphorylated proteins, such as nucleolar and coiled-body phosphoprotein 1 (NOLC1) and treacle protein (TCOF1), but also proteins with only one identified pyrophosphorylation-site, for example myosin 9 (MYH9; **Figure 5e**). To expand the GO analysis to include protein pyrophosphorylation, the GO-term “signaling *via* pyrophosphorylation” was created for the 57 known pyrophosphoproteins. “Signaling *via* pyrophosphorylation” significantly stood out as a GO term (P-value of 1.6 x 10^-13^). The significant overrepresentation of pyrophosphorylated proteins in the proteomic data sets therefore raised the possibility that additional pyrophosphoproteins may be found among 5PP-InsP_5_/1,5(PP)_2_-InsP_4_ interactors.

### Enriched proteins can be linked to pyrophosphorylation–based signal transduction

Pyrophosphorylation sites are typically located within intrinsically disordered regions (IDR) and are pre-phosphorylated by acidophilic or proline-directed Ser/Thr kinases^26,27^. We therefore tested the 863 proteins that were enriched with biotin-5PCP-InsP_5_ and biotin-1,5(PCP)_2_-InsP_4_ (Mg^2+^, K_D_^app^ < 5 µM) for acidophilic or proline-directed Ser/Thr kinase motifs using Scansite 4.0 (www.scansite4.mit.edu)^62^, followed by an analysis of protein disorder with IUPred (www.iupred3.elte.hu/)^63^. About 35% (305 proteins) of the enriched targets fulfilled both criteria (**Figure 6a**). The proteins were then compared with previous pyrophopsphoproteomic analyses of HEK293T cell lysates, revealing that 11 proteins had tryptic peptides that displayed a characteristic neutral loss during collision induced dissociation (CID), which subsequently triggered an electron transfer high-collision dissociation (EThcD) scan. However, they could not be confirmed in the manual assignment afterwards^26^. Site-assignment was not possible in these cases because the EThcD spectra showed insufficient sequence coverage, with crucial fragments missing due to co-elution with corresponding bisphosphorylated peptides, or the peptides exhibited poor sequence-dependent ionization and fragmentation. We reasoned that a targeted approach would increase the chances of detecting these pyrophosphorylation sites by increasing the number of spectra per target peptide and overcoming the masking by other species.

**Figure 6.**
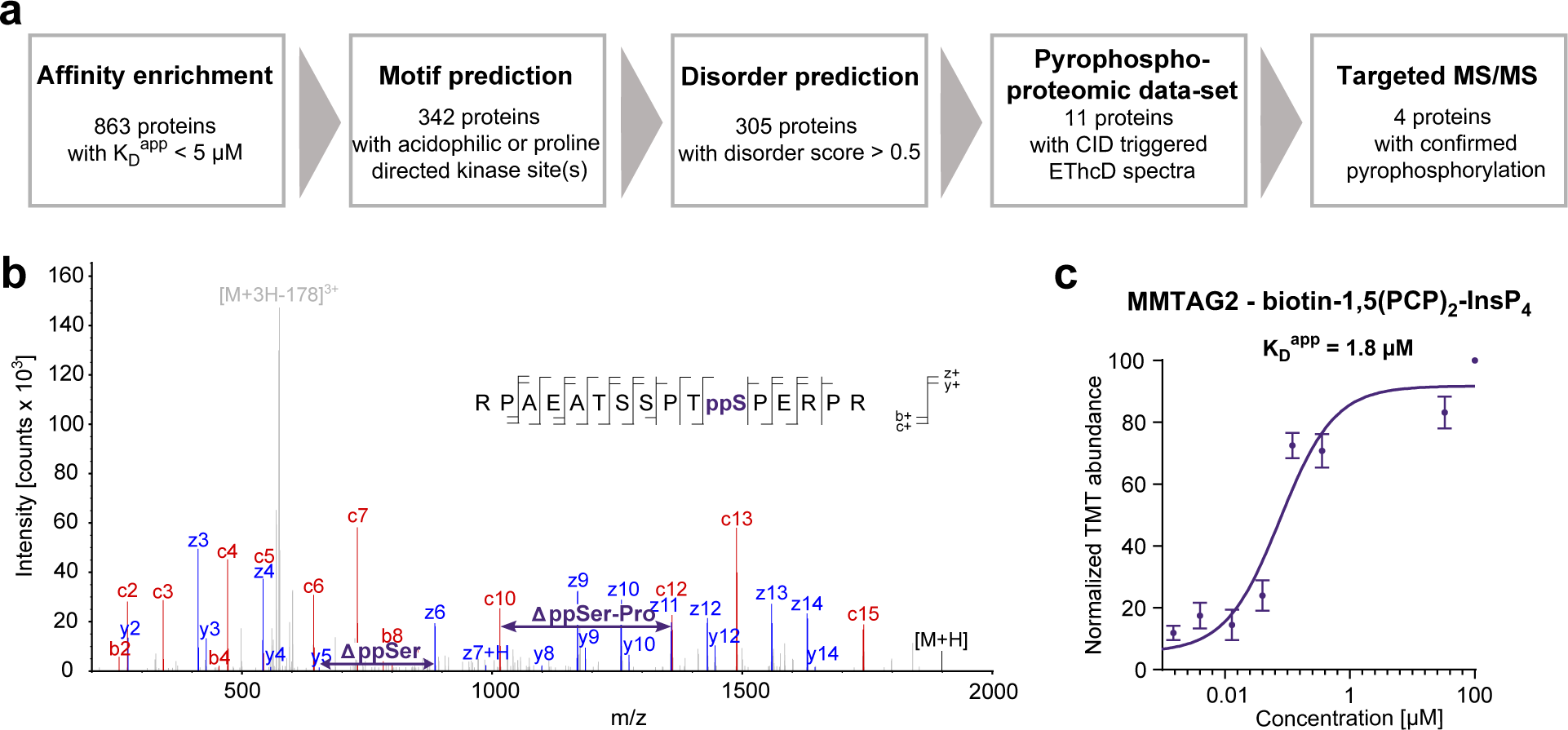
Targeted search for pyrophosphorylated proteins. **a**: Workflow to predict potential pyrophosphorylated proteins from the affinity enrichment data. All 863 proteins enriched by biotin-5PCP-InsP_5_ and biotin-1,5(PCP)_2_-InsP_4_ (K_D_^app^ < 5 µM) from Mg^2+^_nucl_ lysate were searched for acidophilic or proline-directed serine/threonine kinase site(s) and disordered regions. Subsequently, the 305 proteins were compared with a previous global MS/MS analysis, and the resulting 11 proteins were specifically targeted in an MS approach, confirming four pyrophosphoproteins. **b**: EThcD spectrum of the pyrophosphorylated peptide of multiple myeloma tumor-associated protein 2 (MMTAG2) obtained from a complex sample using a targeted MS approach. Fragment ions that are critical for the assignment of the pyrophosphorylation site at Ser220 are shown in red (b- and c-ion series) or blue (y- and z-ion series). **c**: Binding curve for the enrichment of MMTAG2 with biotin-1,5(PCP)_2_-InsP_4_ from Mg^2+^_nucl_ lysate. Every data point represents the mean of three replicates and the error bars depict the SEM.

Using such a targeted MS approach with an enriched HEK293T cell lysate, four proteins with five pyrophosphorylation sites could indeed be confirmed. Exemplarily, the EThcD spectrum of the multiple myeloma tumor-associated protein 2 (MMTAG2) is shown in **Figure 6b**, confirming the identity of the pyrophosphorylation site on serine 220. The corresponding affinity curve of MMTAG2 binding to biotin-1,5(PCP)_2_-InsP_4_ indicated a K_D_^app^ value of 1.8 µM (**Figure 6c**). The other confirmed pyrophosphoproteins are CLIP-associating protein 2 (CLASP2; Ser 324), RNA polymerase-associated protein CTR9 homolog (CTR9; Ser 1017 and Ser 1021), and pre-mRNA-splicing factor ATP-dependent RNA helicase DHX16 (DHX16; Ser 103). The results demonstrate that potential pyrophosphoproteins can be identified using the affinity enrichment method, which can complement the mass spectrometry pipeline currently used for detection of pyrophosphorylation.

## DISCUSSION

Affinity enrichment is a well-established technique in chemical biology, frequently employed to identify binding partners of biologically relevant molecules. In many cases, the binding affinities of these interactions are of high relevance, as they determine the strength and duration of the interaction and are central for comprehending the underlying molecular mechanisms. However, binding affinities are still predominantly determined at the biochemical level, using purified proteins. Here, we introduce the ability to determine binding affinities into a rapid quantitative mass spectrometric approach and applied this method to the group of inositol pyrophosphate messengers.

A desymmetrization strategy was employed to generate affinity reagents biotin-1PCP-InsP_5_ and biotin-1,5(PCP)_2_-InsP_4_ affinity reagents, immobilized *via* a biotin-PEG_5_ linker. Application of these reagents to cell lysates, alongside biotin-InsP_6_ and biotin-5PCP-InsP_5_, demonstrated concentration-dependent affinity enrichment of known binding proteins using immunoblotting. These results motivated the application of mass spectrometry, to determine apparent binding constants on a proteome-wide level. This quantitative analysis was carried out using a mammalian cell lysate, which had been separated into cytosolic and nuclear fraction, and which either contained Mg^2+^ ions or was depleted for di- and trivalent cations. Hill-like curves were obtained for more than a thousand binding proteins and apparent dissociation constants (K_D_^app^) were derived. The good performance of the reagents, and the quantitative nature of the method, was confirmed by the retention and K_D_^app^ determination of InsP and the PP-InsP metabolizing enzymes IP6K1, IP6K2, PPIP5K2, DIPP1, and DIPP2^64^. As such, the data reported here constitutes a valuable resource for researchers in the field, and will hopefully inspire follow-up investigations on PP-InsP signaling mechanisms in mammalian systems. Importantly, for these follow-up studies, the strength of the PP-InsP protein interaction (i.e. K_D_^app^) can aid in the prioritization of these often times demanding biochemical/cell biological experiments. Furthermore, the affinity reagents should be applied to different organisms, so that the conservation (and divergence) of PP-InsP signaling can be studied in various biological contexts.

As anticipated, a wide range of apparent binding constants was observed in our proteomics experiments, ranging from K_D_^app^ values between 17 nM to 25 µM. To aid the subsequent analysis, we defined cut-off values based on averaged cellular concentrations of InsPs/PP-InsPs. It is important to note that these cut-off values are somewhat arbitrary and cannot accurately reflect the local concentrations of these messengers. Even though many observed binding interactions lie above the defined thresholds, these interactions may still be biologically relevant, given the appropriate cellular surroundings. A method that reliably measures the concentration of the different PP-InsPs with subcellular resolution – akin to the reporters used for phosphatidyl inositols – would be of great use. Such a reporter could also provide insight into another unexplained phenomenon: many interactions of high affinity were observed for 1,5(PP)_2_-InsP_4_ with nucleolar proteins, yet, it is not known if this messenger actually localizes to this compartment in intact cells.

Somewhat unexpectedly, the proteomics data indicated that biotin-1PCP-InsP_5_ exhibited weaker interactions with numerous proteins, compared to the other inositol pyrophosphates. A possible interpretation is that 1PP-InsP_5_ is not primarily involved in cellular signaling, but rather just constitutes an intermediate during dephosphorylation of 1,5(PP)_2_-InsP_4_. The concept, that the pathway for 1,5(PP)_2_-InsP_4_ turnover comprises a predominantly cyclical interconversion (InsP_6_ → 5PP-InsP_5_ → 1,5(PP)_2_-InsP_4_ → 1PP-InsP_5_ → InsP_6_) has been postulated before^7,9^. In this scheme, 1PP-InsP_5_ is predominantly generated by the dephosphorylation of 1,5(PP)_2_-InsP_4_ by DIPPs, and not *via* PPIP5K catalyzed phosphorylation of InsP_6_. Consistent with this model, our data indicates that 1PP-InsP_5_ is a less dominant signaling molecule, compared to 5PP-insP_5_ and 1,5(PP)_2_-InsP_4_, in mammalian cells.

Not only the arrangement of the pyrophosphate groups around the myo-inositol scaffold is important for recognition, but also the speciation (i.e. protonation or metal-coordination) of the different PP-InsPs. The highly phosphorylated inositols have a special relationship with Mg^2+^ ions, the most abundant divalent metal-ions in cells^53^. Coordination of Mg^2+^-ions strongly influences the solubility of InsP_6_/PP-InsPs, their charge, and their hydration shells, which all, in turn, can have an effect on the binding strength towards different target proteins^49,51,65^. Depending on the binding sites that are targeted by the PP-InsPs, it may be necessary to remove the Mg^2+^ ions for protein binding – a process that will likely be energetically costly. In the structurally characterized examples of PP-InsP-protein complexes, Mg^2+^ ions are rarely found. Instead, a large number of positively charged side chains are involved in neutralizing the charge of the highly negatively charged ligand. It would therefore be interesting to investigate, if cells employ regulatory mechanisms that can restrict local Mg^2+^ ion availability, thereby strengthening a specific set of PP-InsP protein interactions.

Interestingly, many PP-InsP-protein interactions were strengthened by the presence of Mg^2+^ ions, suggesting that the metal ions may play a role in stabilizing either the protein, the PP-InsP ligand, or the protein-ligand interface. A possible scenario is that certain proteins may actually prefer binding to PP-InsPs in a Mg^2+^-coordinated, “flipped” conformation, in which five substituents of the *myo*-inositol ring are placed in axial positions. It was previously reported that app. 30% of 1,5(PP)_2_-InsP_4_ adopts such a flipped conformation in the presence of Mg^2+^-ions^51^. However, to date no characterized example of flipped PP-InsP-protein complex exists. With the advancement of high-resolution structural methods though, such as cryo-electron microscopy and NMR, such characterization is not out of reach anymore.

Another, maybe more feasible, scenario is that Mg^2+^-ions help to bring together the negatively charged PP-InsPs and protein sequences that are targets of pyrophosphorylation, as pyrophosphorylation sites are commonly found in serine-rich polyacidic stretches. Mg^2+^ ions had been reported to be essential for this unusual phosphoryl transfer reaction and likely acts as a molecular glue to bring together the two negatively charge reacting partners. Consistent with this, numerous pyrophosphorylation targets were enriched with biotin-5PCP-InsP_5_ and biotin-1,5(PCP)_2_-InsP_4_ in the presence of Mg^2+^, which implies a specific interaction between the PP-InsP-magnesium complex and negatively charged pyrophosphorylation sequences. A similar observation was made previously for 5PP-InsP_5_ interacting proteins from *S. cerevisiae*, where several pyrophosphorylation targets were retained upon the addition of Mg^2+^-ions^45^. Although the involvement of 1,5(PP)_2_-InsP_4_ in pyrophosphorylation chemistry remains speculative, it appears reasonable that this molecule is also capable of transferring its β-phosphoryl group. Further research is required to elucidate the contribution of distinct PP-InsP messengers to protein pyrophosphorylation. With the development of new chemical methods to obtain different PP-InsP isomers, and the recent implementation of a mass spectrometry method to detect pyrophosphorylation sites, such studies have become feasible^15,26^.

Given the significant enrichment of pyrophosphoproteins in our data sets, we investigated the possibility of that additional targets of pyrophosphorylation were retained by the affinity reagents. Indeed, by using a targets mass spectrometry approach for 11 putative pyrophosphorylation sites, we could confirm five novel sites on four proteins. Because the detection of pyrophosphorylation sites on peptides still remains a challenge (due to low abundance, poor ionization, and conflicting phosphorylation patterns), it is necessary to enrich pyrophosphorylated peptides in the currently implemented mass spectrometry workflow. This enrichment relies on sequential elution from immobilized metal affinity chromatography (SIMAC), followed by fractionation. The affinity reagents presented here could offer an alternative approach to enrich pyrophosphorylation targets at the protein level, especially when applied to nuclear or nucleolar lysates.

One limitation of the current method is the capture of protein complexes, such as ribonucleoprotein complexes. Within these assembled structures, not every protein interacts with the InsP or PP-InsP ligand. These complexes will bias the gene ontology analysis, since all identified protein components are included in the data query input. The enrichment of indirect binding partners could be reduced by forming a covalent bond between the ligand and its direct binding partner, allowing for much more stringent washes. Such a covalent capture could, for example, be achieved by incorporating photo-crosslinkers into the PEG linker region.

In sum, the reagents and the method described here constitute a resource to the community. It can be used as a starting point for subsequent investigation of a certain protein, or as a tool to annotate PP-InsP signaling pathways in other cell lines or organisms. One could also focus future endeavors on specific organelles, such as the nucleolus, where many pyrophosphoproteins are localized, to better understand the regulation of this unusual phosphorylation mode. Considering the various signaling modes of PP-InsPs, and their ubiquitous occurrence in eukaryotes, many of their signaling functions still remain to be discovered.

## METHODS

Detailed methods can be found in the supplemental materials and methods.

## SUPPLEMENTAL INFORMATION

The supplemental information contains supplementary figures, supplementary materials and methods, and supplementary tables. All of the files are available online.

## DATA AVAILABILITY STATEMENT

The mass spectrometry Raw data and ProteomeDiscoverer outputs were deposited with the ProteomeXchange Consoritum partner repository jPOSTrep under the accession code JPST003145^66^ and PXD052682.

The R scripts used for the analysis of quantitative proteomic data are available via Zenodo at https://doi.org/10.5281/zenodo.11388151

## ACKNOWLEDGEMENTS

We thank Peter Schmieder for his 2D-NMR expertise and Lena von Oertzen for performing the cell culture experiments. We also thank Leonie Kurz for providing SYT1^C2B^, and Meike Amma and Simon Bartsch for their help with the high-resolution mass spectrometry. We thank all group members of the Fiedler group for reading and discussing the manuscript. Annika Richter gratefully acknowledges funding by the Studienstiftung des Deutschen Volkes and the Deutsche Forschungsgemeinschaft (TRR186).

## AUTHOR CONTRIBUTIONS

Conceptualization, A. Richter, D. Furkert, M. Ruwolt, and D. Fiedler; Methodology, A. Richter, D. Furkert, M. Ruwolt, S. Lampe, and D. Fiedler; Formal Analysis, A. Richter, M. Ruwolt, and S. Lampe; Investigation, A. Richter, M. Ruwolt, and S. Lampe; Writing – Original Draft, A. Richter and D. Fiedler; Writing – Review & Editing, A. Richter, and D. Fiedler; Visualization, A. Richter, S. Lampe, and D. Fiedler; Supervision, F. Liu and D. Fiedler; Funding Acquisition, L. Liu and D. Fiedler.

## DECLARATION OF INTEREST

The authors declare no competing interests.

## Notes

### Competing Interest Statement

The authors have declared no competing interest.

