## Supplementary material for "Proteome-wide quantification of inositol pyrophosphate-protein interactions": SI Figures

**a**

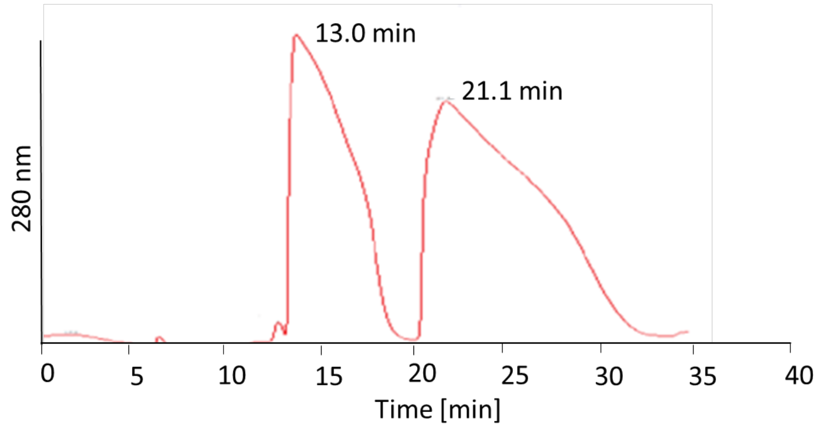


**b**

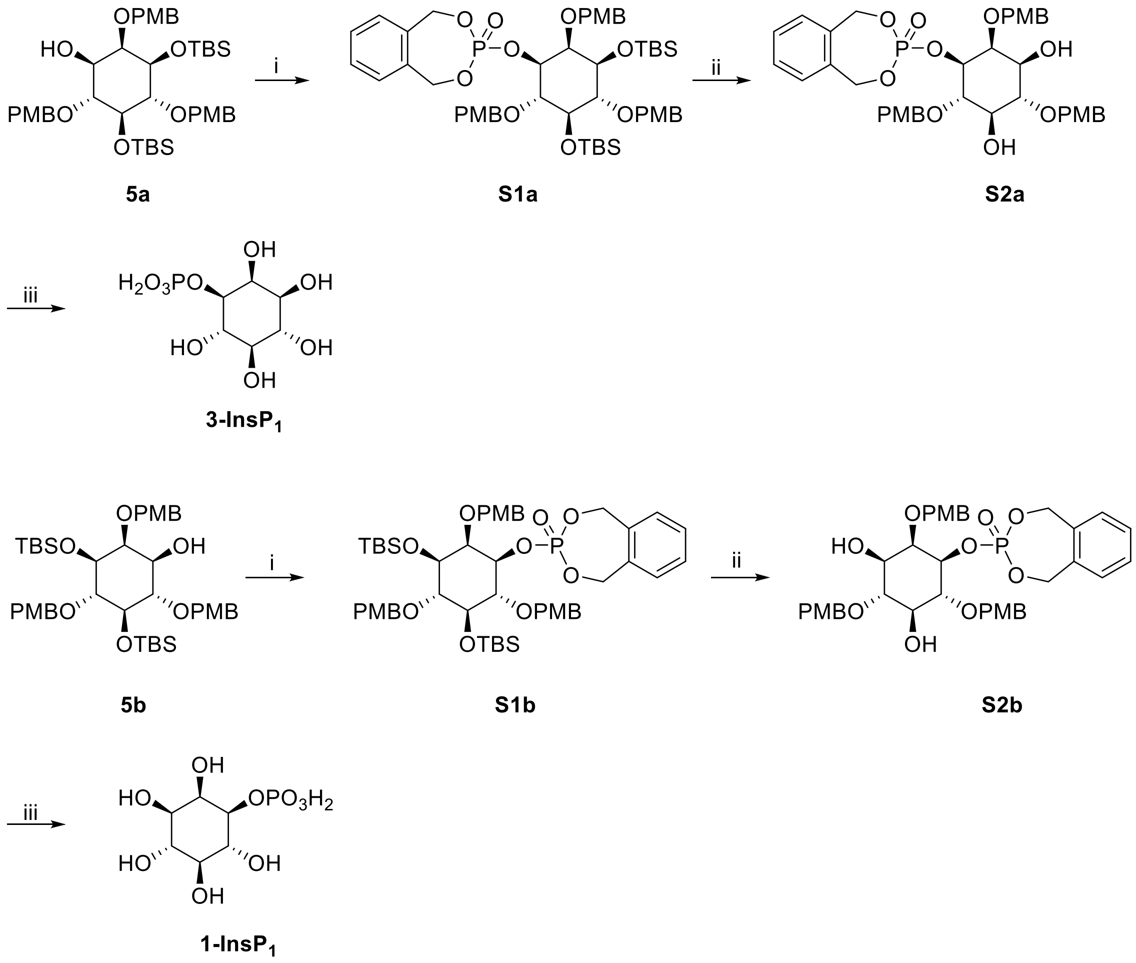


**Supplementary Figure S1: Preparation and validation of enantiopure precursors.**
**a:**Chiral separation of 200 mg enantiomer mix **5a** and **5b** on a Registech (S, S) Whelk-O 1 chiral column.
**b**:Synthesis of 1/3-InsP$_1$ for assignment of **5a** and **5b**: (i) **16**, 4,5-dicyanoimidazole, ACN, then mCPBA, 89% (ii) TBAF, THF, 97% (iii) Pd/C, *t*BuOH/$H_2O$, quantitative.

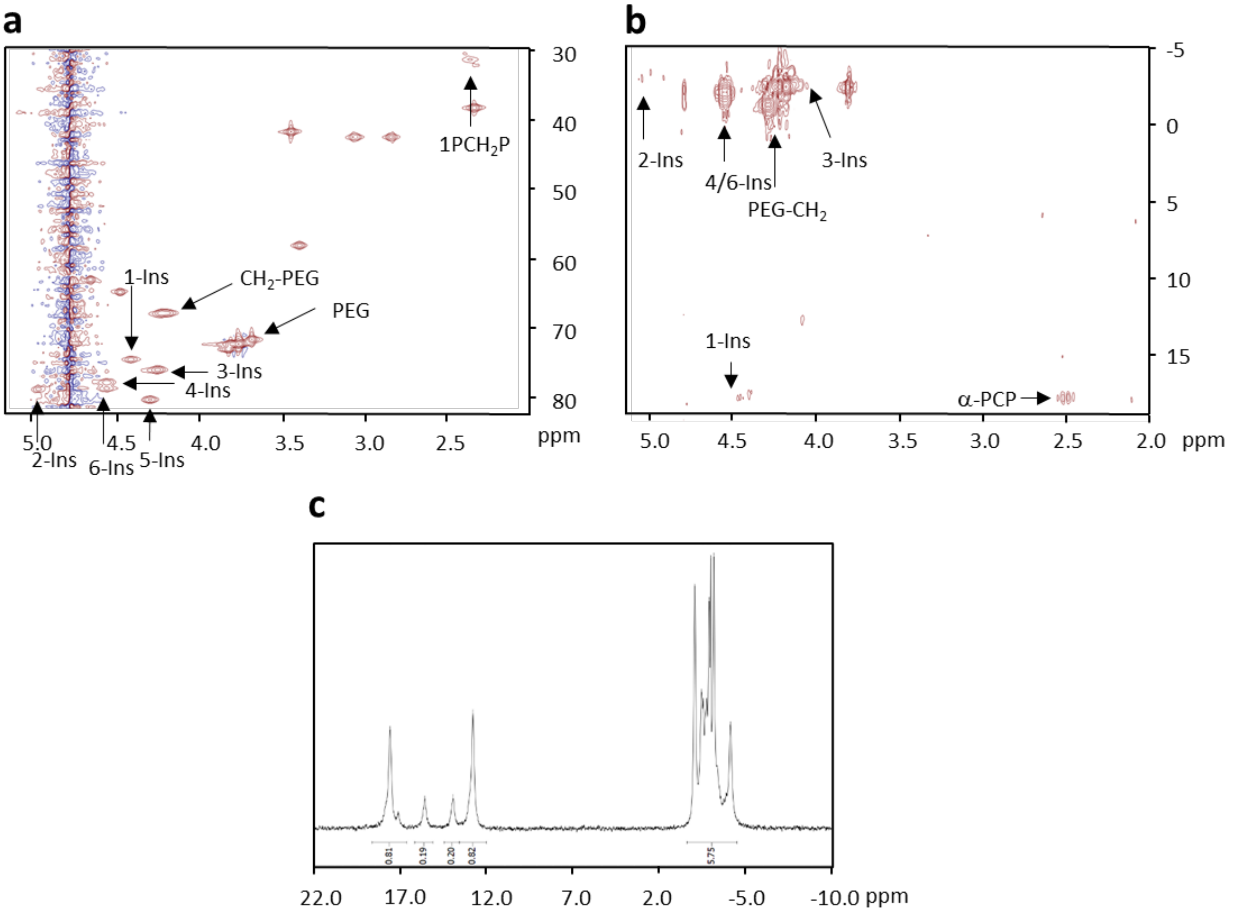


**Supplementary Figure S2: 2D-NMR experiments confirming the correct attachment of linker at the 3- or 5-position and PCP group at 1-position for biotin-3/5L-1PCP-InsP$_5$ (1a and 1b).**

**a**: $^1$H-$^{13}$C-DEPT-CLIP-COSY NMR experiment. The X-axis shows the $^1$H-dimension and the y-axis displays the $^{13}$C-dimension. C-H correlation of the *myo*-inositol scaffold as well as other interesting correlations like PCH$_2$P, PEG, and CH$_2$-PEG are assigned by black arrows.

**b**: $^{31}$P-HMBC NMR experiment. The X-axis shows the $^1$H-dimension and the y-axis displays the $^{31}$P-dimension. Correlation for the *myo*-inositol scaffold as well as CH$_2$-PEG and α-PCH$_2$P correlation are highlighted with black arrows. β-PCH$_2$P correlation is not visible due to low material availability and can only be detected in the 1D-$^{31}$P NMR. **C**: $^{31}$P-NMR experiment. The α- and β-phosphates of the 1PCP group show a ratio of 20% 5L-compound and 80% 3L-compound. Together A, B, and C confirm the attachment of linker amidate at 3-OH or 5-OH and PCP-amidate at 1-OH.

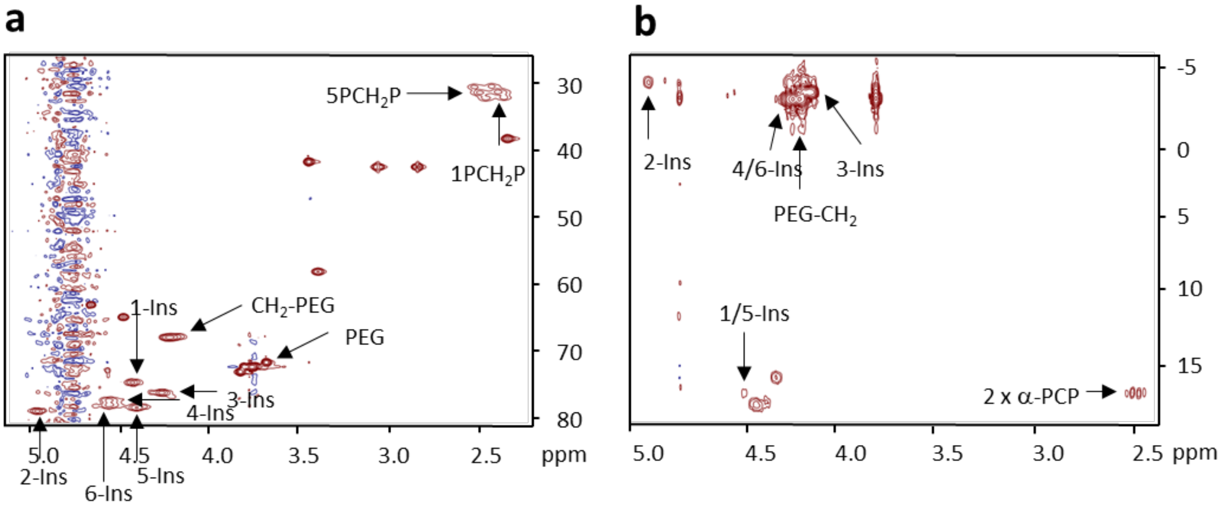


**Supplementary Figure S3: 2D-NMR experiments confirming the correct attachment of linker at the 3-position and PCP groups at 1-, and 5-position for biotin-3L-1,5(PCP)$_2$-InsP$_4$ (2).**

**a**: $^{1}$H-$^{13}$C-DEPT-CLIP-COSY NMR experiment. The X-axis shows the $^{1}$H-dimension and the y-axis displays the $^{13}$C-dimension. C-H correlation of the *myo*-inositol scaffold as well as other interesting correlations like PCH$_2$P, PEG, and CH$_2$-PEG are assigned by black arrows.

**b**: $^{31}$P-HMBC NMR experiment. The X-axis shows the $^{1}$H-dimension and the y-axis displays the $^{31}$P-dimension. Correlation for the myo-inositol scaffold as well as CH$_2$-PEG and $\alpha$-PCH$_2$P correlation are highlighted with black arrows. $\beta$-PCH$_2$P correlation is not visible due to low material availability and can only be detected in the 1D-$^{31}$P NMR. Together A and B confirm the attachment of linker amidate at 3-OH and PCP-amidate at 1- and 5-OH.

**a**

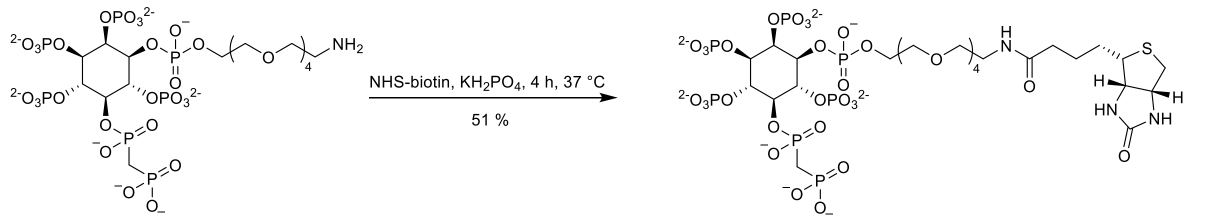


**18** **3b**

**b**

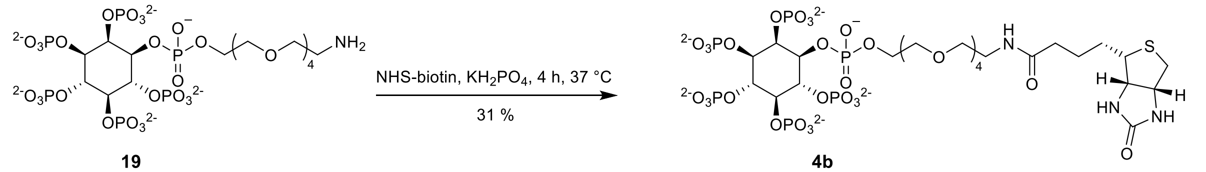


**19** **4b**

**Supplementary Figure S4: Synthesis of biotin-InsP$_6$ and biotin-5PCP-InsP$_5$ with the linker attachment either at position 1 or 3.**

For simplicity, only the 1-linked enantiomer is shown. **18** and **19** were obtained according to a reported procedure[1].

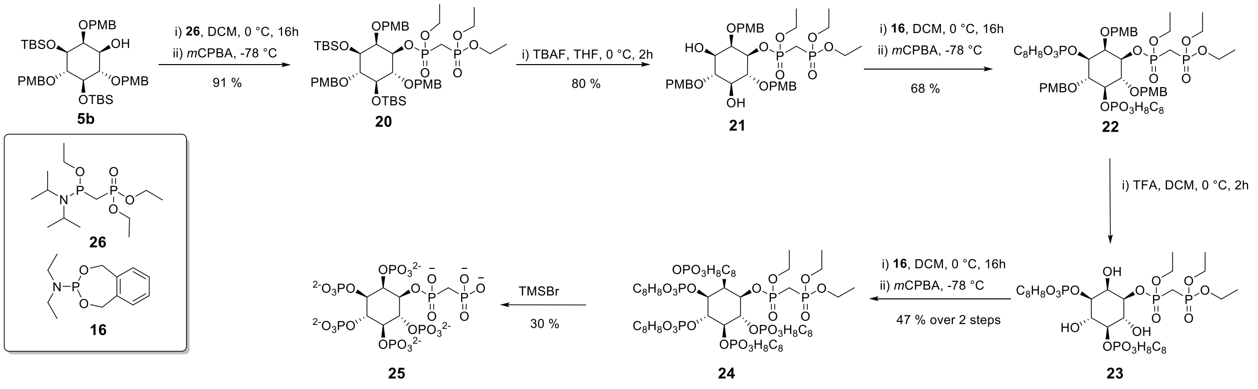


**Supplementary Figure S5: Synthesis of 1PCP-InsP$_5$ starting with pure enantiomer 5a.** Phosphoramidite **26** was synthesized following a published procedure[2].

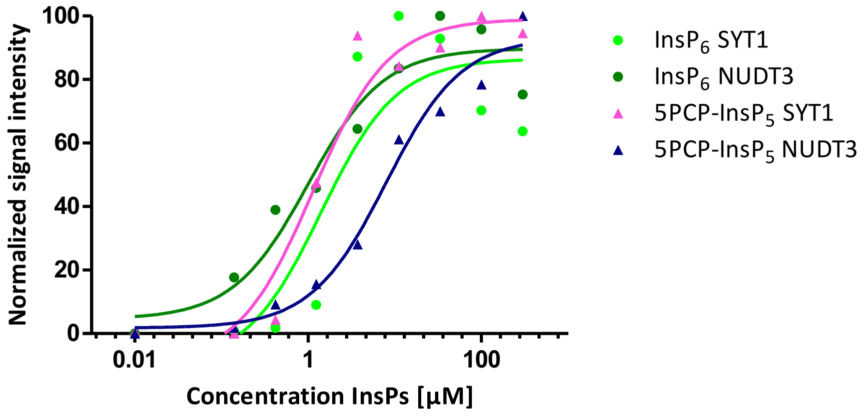


**Supplementary Figure S6: Concentration-dependent affinity enrichment and western blot analysis**. The obtained signals were quantified and plotted against the biotin-$InsP_6$ or biotin-5PCP-$InsP_5$ concentration.

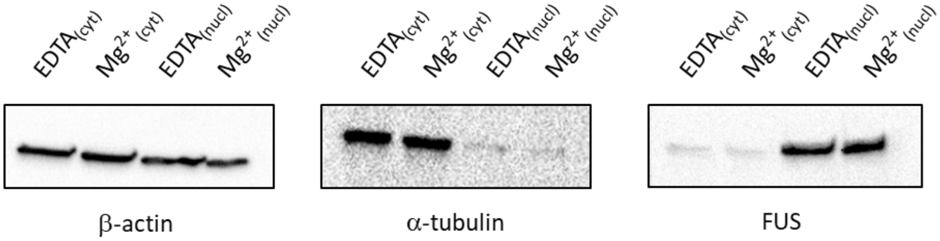


**Supplementary Figure S7: Cell lysis and separation of cytosolic and nuclear fraction.**
Western blot analysis using cytosolic markers β-actin and α-tubulin and nuclear markers β-actin and FUS.

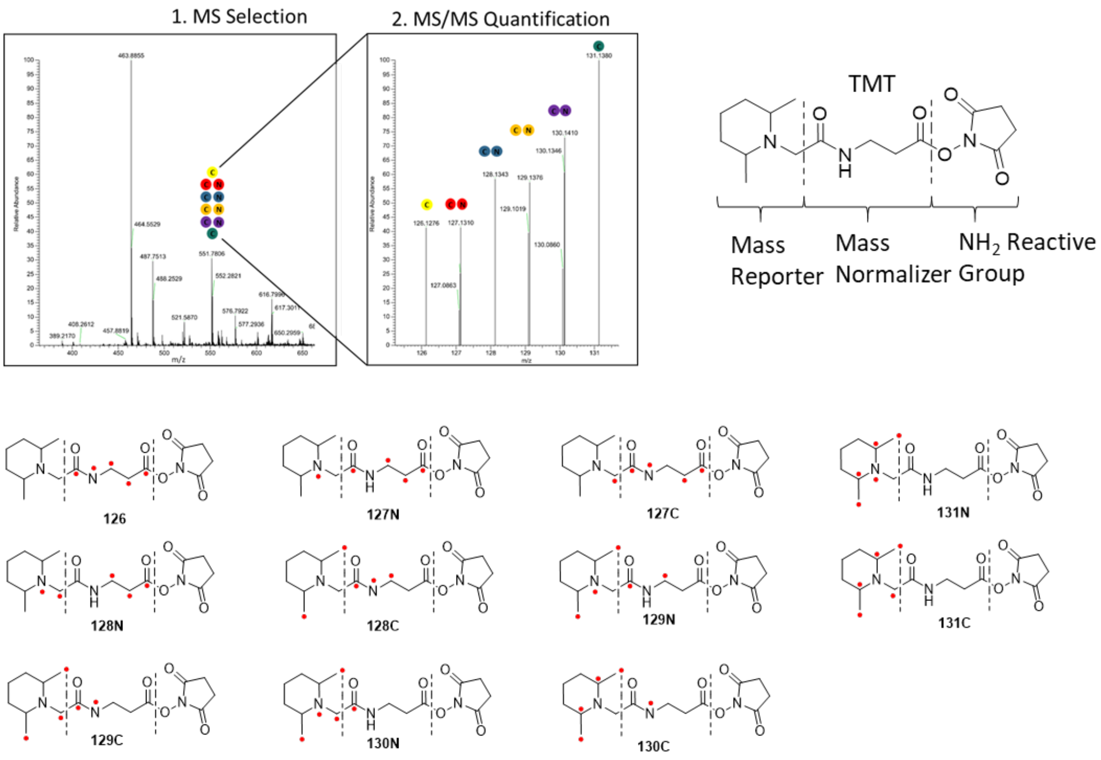


**Supplementary Figure S8: Principle of 11-plex tandem mass tag (TMT) isobaric labeling.**
Each TMT molecule has a different isotopic label pattern, indicated in red dots at the carbon or nitrogen atom. N-termini of peptides react with the $NH_2$ reactive group, resulting in a unique label for each dataset. All sample sets can be pooled subsequently and measured in one experiment. Upon MS 1 selection all peptides elute with the same m/z. However during MS 2, the tandem mass tag gets fragmented and each tag results in another signal, enabling retrospective assignment of the original sample set.

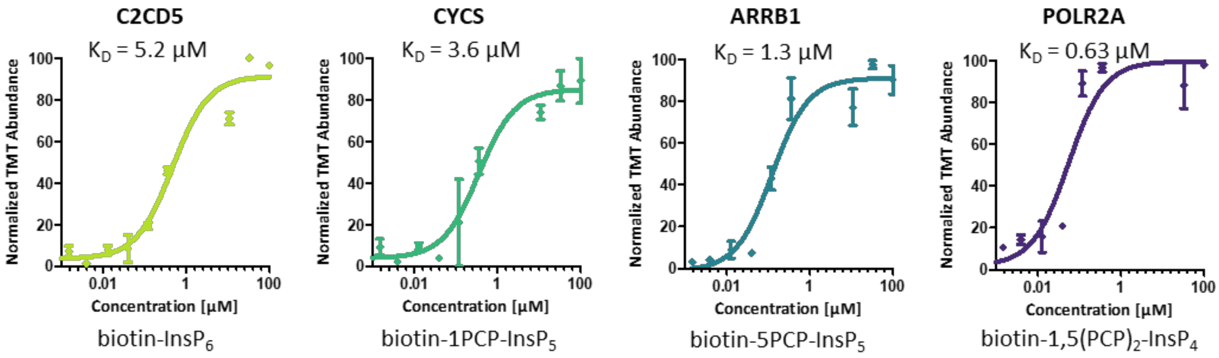


**Supplementary Figure S9: $K_D^{app}$ binding curves for the (PP)-InsP binding proteins C2CD5, CYCS, ARRB1, and POLR2I (from left to right).**

The color indicates the lysate conditions and the affinity probe used is shown under the curves. Every data point represents the mean of three replicates and the error bars depict the SEM.

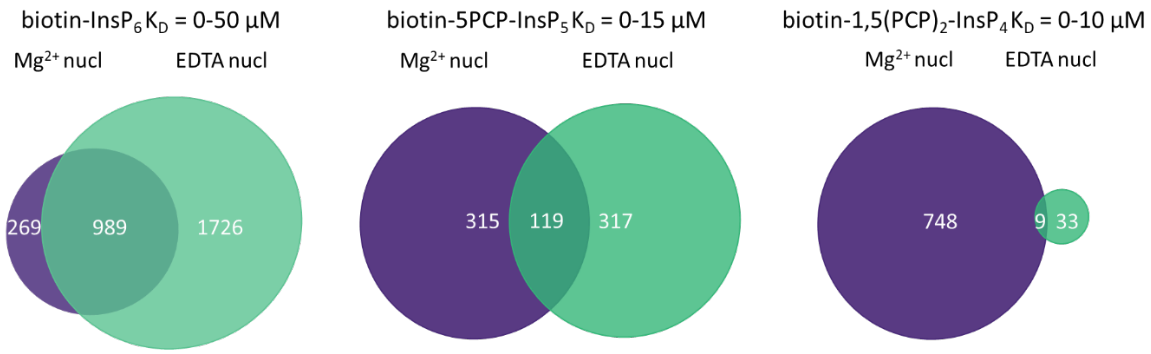


**Supplementary Figure S10: Venn diagrams of all proteins quantified for biotin-InsP$_6$ ($K_D^{app}$ = 0 – 50 µM), biotin-5PCP-InsP$_5$ ($K_D^{app}$ = 0-15 µM), and biotin-1,5(PCP)$_2$-InsP$_4$ ($K_D^{app}$ = 0 – 10 µM).** Numbers are shown in the circles and colors indicate the lysate conditions[3].
