## Supplementary material for "Proteome-wide quantification of inositol pyrophosphate-protein interactions": SI Materials and Methods

### TABLE OF REAGENTS AND RESOURCES

| Reagent or Resource | Source | Identifier |
| --- | --- | --- |
| Antibodies | | |
| COPS5/JAB1 | Proteintech | Cat# 27511-1-AP |
| SYT1 | Proteintech | Cat# CL488-68043 |
| PRPS1 | Proteintech | Cat# 15549-1-AP |
| NUDT3 | Proteintech | Cat# 20542-1-AP |
| Polyclonal rabbit HRP | Cell Signalling Technology | Cat# 7074; RRID: AB_2099233 |
| FUS | Santa Cruz | Cat# sc-47711 |
| b-actin | Santa Cruz | Cat# sc-47778 |
| a-tubulin | Santa Cruz | Cat# sc-8035 |
| Chemicals, Peptides and Recombinant Proteins | | |
| Dulbecco's Modified Eagle Medium (DMEM) | Gibco | Cat# 11960-044 |
| Penicillin/streptomycin | Gibco | Cat# 15140-122 |
| L-Glutamine | Gibco | Cat# 25030-024 |
| DPBS | Gibco | Cat# 14190-094 |
| Pierce IP lysis buffer | Thermo Scientific | Cat# 87787 |
| PhosSTOP Phosphatase inhibitor | Sigma-Aldrich | Cat# 4906845001 |
| cOmplete, Mini, EDTA-free Protease Inhibitor Cocktail | Sigma-Aldrich | Cat# 11836170001 |
| Affi-Gel 15 Gel | Bio-Rad | Cat# 1536051 |
| Triethylammonium bicarbonate buffer | Sigma-Aldrich | Cat# T7408 |
| Trypsin | Serva | Cat# 5763_37286 |
| Lys C | Fujifilm | Cat# 121-05063 |
| CAA | Sigma-Aldrich | C0267-100g, CAS: 79-07-2 |
| TCEP | Sigma-Aldrich | Art. No.; 646547-1ML, CAS: 51805-45-9 |
| biotin-$InsP_6$ | This study | N/A |
| biotin-5PCP-$InsP_5$ | This study | N/A |
| biotin-1PCP-$InsP_5$ | This study | N/A |
| biotin-1,5$(PCP)_2$-$InsP_4$ | This study | N/A |
| 1/3***L***-$InsP_6$ | Furkert *et al.* 2020 | N/A |
| 1/3***L***-5PCP-$InsP_5$ | Furkert *et al.* 2020 | N/A |
| $InsP_6$ | SciChem | Cat# 6-0-123456-Na |
| 5PCP-$InsP_5$ | Wu *et al.* 2013 | N/A |
| 5PP-$InsP_5$ | Puschmann *et al.* 2019 | N/A |
| 1PCP-$InsP_5$ | Wu *et al.* 2014 | N/A |
| 1,5$(PCP)_2$-$InsP_4$ | Hostachy *et al.* 2021 | N/A |
| 1,5$(PP)_2$-$InsP_4$ | Puschmann *et al.* 2019 | N/A |
| GST-hDipp1 | Wu *et al.* 2015 | N/A |
| MBP-SPX XPR1 | Li *et al.* 2020 | N/A |
| SYT1 | Wangt *et al.* 2014 | N/A |
| 5-phenyl-1H-tetrazole | VWR | B25664; CAS: 18039-42-4 |
| p-Toluenesulfonic acid monohydrate | Sigma Aldrich | 416665; CAS: 75-91-2 |

| | | |
|---|---|---|
| 3-Chloroperbenzoic acid | Sigma Aldrich | 273031; CAS: 937-14-4 |
| o-Xylylene N,N-diethylphosphoramidite | Sigma Aldrich | 393835; CAS: 82372-35-8 |
| 1H-tetrazole | Sigma Aldrich | 88158; CAS: 288-94-8 |
| 4,5-Dicyanimidazol | Sigma Aldrich | 324132; CAS: 1122-28-7 |
| Tetrabutylammoniumfluorid Trihydrat | Sigma Aldrich | 86872; CAS: 87749-50-6 |
| Trimethylsilyl bromide | Sigma Aldrich | 194409; CAS: 2857-97-8 |
| Streptavidin Sepharose | GE Healthcare Life Sciences | Cat#17-5113-01 |
| Medronic acid | Sigma Aldrich | Cat#64255-1G-F |
| Critical Commercial Assays | | |
| TMT 10plex™ | Thermo Fisher Scientific | Cat# 90111 |
| TMT11-131C label reagent | Thermo Fisher Scientific | Cat# A37724 |
| Pierce BCA-Protein Assay | Thermo Fisher Scientific | Cat# 24612 |
| Pierce Silver Stain kit | Thermo Fisher Scientific | Cat# 23227 |
| TNIK kinase enzyme system | Promega | Cat# V4158 |
| Kinase-Glo Plus | Promega | Cat# V3771 |
| Instant coomassie | Abcam | Cat# ab119211 |
| Deposited Data | | |
| HEK293T raw and analyzed data | This study | |
| Experimental Models: Cell lines | | |
| HEK293T | ATCC | CRL-3216 |
| Software and Algorithms | | |
| Proteome Discoverer 3.0 | Thermo Fisher Scientific | |
| BioRender | N/A | https://www.biorender.com/ |
| MestReNova v.10.0.2 | N/A | https://mestrelab.com/ |
| RStudio 2023.06.2 | Posit Software, PBC | https://www.r-studio.com/de/ |
| TOPSPIN v3.5 | N/A | https://www.bruker.com/en/products-and-solutions/mr/nmr-software/topspin |
| Scansite4.0 | | https://scansite4.mit.edu/#home |
| IUPred3 | | https://iupred3.elte.hu/ |
| BioVenn | | https://www.biovenn.nl/index.php |
| Prism 5.04 | | https://www.graphpad.com |
| Image Lab 6.1.0 | Bio-Rad | https://www.bio-rad.com/de-de/product/image-lab-software |
| ChemDraw 18.2.0.48 | Perkin Elmer | https://www.additive-net.de/de/software/produkte/perkinelmer/chemdraw/neu#version-18 |
| Inkscape 1.3.2-4 | | https://inkscape.org/de/ |
| Orbitrap Fusion Lumos Tribrid Mass Spectrometer | Thermo Fisher Scientific | |

| | | |
|---|---|---|
| FAIMS Pro Interface | Thermo Fisher Scientific | Cat# FMS02-10001 |
| Orbitrap Fusion Tribrid Mass spectrometer | Thermo Fisher Scientific | |
| Loading column 0.075 x 70 mm | Thermo Fisher Scientific | Cat# 164946 |
| C18 column: Poroshell 120-EC-C18, 2.7 µm (inhouse packed) | Agilent | |
| PlatePrep 96-well vacuum distributor | Sigma Aldrich | Cat# 57192-U |
| Multiscreen® 96 well Plate, hydrophilic PVDF membrane | Sigma Aldrich | Cat# MSBV1210 |

**Materials availability**

Requests for unique/stable reagents should be directed to Dorothea Fiedler. Availability may be limited due to multistep synthesis.

### EXPERIMENTAL DETAILS

#### BIOCHEMICAL AND PROTEOMICS EXPERIMENTS

##### Cell culture

HEK293T cells (female human origin) were cultured in Dulbecco´s Modified Eagles´Medium (DMEM), with 10 % FBS, Penicillin-Streptomycin (100 U/mL), and Gliutamine (2 mM) in a 5 % humidified $CO_2$ incubator at 37 °C. The cells were received from the american type culture collection (ATCC) and tested for mycoplasma before use.

##### Cell lysate preparation

Whole-cell extracts for validation experiments were prepared as described previously with certain modifications[2,3]. Briefly, HEK293T cells were grown to 90 % confluency in 15 cm dishes. Cells were washed twice with ice-cold 0.9 mM NaCl in $H_2O$ (10 mL) and lysed with either Pierce™ IP Lysis buffer (TRIS EDTA conditions), buffer 2 (150 mM NaCl, 25 mM HEPES pH 7.4 at 4 °C, 1 mM EDTA, 1 % NP-40, HEPES EDTA conditions)) or buffer 3 (150 mM NaCl, 25 mM HEPES pH 7.4 at 4 °C, 1 mM $MgCl_2$, 1 % NP-40, HEPES $Mg^{2+}$ conditions)). 2.7 mL lysis buffer, supplemented with phosphatase and protease inhibitors (Roche PhosStop™ and cOmpleteTM EDTA-free protease inhibitor cocktail) were added, scraped off, added to protein low binding microcentrifuge tubes, and incubated on ice for 10 min. The lysate was centrifuged at 4 °C for 15 min at 21.000 g. The supernatants were combined and protein concentration was measured by Pierce™ BCA Protein Assay kit.

Nuclear and cytosolic cell extracts for affinity enrichment experiments were prepared the following: HEK293T cells were grown to 90 % confluency in 15 cm cell culture dishes. Cells were washed twice with ice-cold 0.9 mM NaCl in $H_2O$ (10 mL), carefully scraped off, added to protein low-binding microcentrifuge tubes, and ice-cold PBS (1 mL) was added. The cells were pelleted for 2 min, 250 g at 4 °C, and the supernatant was removed. The pellet was washed twice with ice-cold PBS (1 mL), the supernatant was removed and the pellet was kept on ice. Afterward, the cells were either lysed by EDTA-containing buffer (150 mM NaCl, 25 mM HEPES pH 7.4 at 4 °C, 1 mM EDTA, 0.1 % Triton™ X-100, Roche PhosStop™, and cOmpleteTM EDTA-free protease inhibitor cocktail) or $Mg^{2+}$-containing buffer (150 mM NaCl, 25 mM HEPES pH 7.4 at 4 °C, 1 mM $MgCl_2$, 0.1 % Triton™ X-100, Roche PhosStop™ and cOmpleteTM EDTA-free protease inhibitor cocktail). 1 mL lysis buffer was added and the cells were incubated on a rotation wheel for 30 min. The cells were centrifuged for 2 min, 250 g at 4 °C. The supernatant, containing the cytosolic proteins, was transferred into falcon tubes, and the lysis was repeated with 1 mL lysis buffer, incubation for 30 min and centrifugation. Subsequently, the supernatant was recovered and added into the same falcon tube. After the lysis of the cytosolic fraction, the cell pellet was dissolved in 1 mL respective lysis buffer followed by sonication at 4 °C (on ice, 50 % output, 0.5 cycle rate, 5 x 30 s, 30 s rest between pulses). The lysate was centrifuged (30 min, 21.000 g, 4 °C) and the supernatant was transferred into falcon tubes. The fractions were aliquoted, snap-frozen, and stored at -70 °C.

The concentration of both lysate fractions was determined by Pierce™ BCA Protein Assay kit.

**Affinity enrichment for proteomics analysis and immunoblotting**

Multiscreen® 96 well Plate, hydrophilic PVDF membrane was placed on the PlatePrep 96-well vacuum distributor and wells were washed with 70 % EtOH (50 µL) followed by two washes with PBS (200 µL, 4 °C). Streptavidin sepharose was added (80 µL) and washed with ice-cold PBS (three times 200 µL). The bottom was dried with a clean wipe and the plate was positioned on a tabletop shaker at 4 °C. For SDS-Page and immunoblotting experiments 20 µL of a 1 mM biotin-(PCP)-InsP solution was dissolved in 150 µL PBS. For the proteomics experiments a threefold-dilution series was prepared according to the table below and the experiments were performed in triplicate:

| | **1** | **2** | **3** | **4** | **5** | **6** | **7** | **8** | **9** | **10** | **11** |
|---|---|---|---|---|---|---|---|---|---|---|---|
| **µM final** | 0 | 0.005 | 0.015 | 0.0045 | 0.137 | 0.41 | 1.23 | 3.7 | 11.1 | 33.3 | 100 |
| **µL PBS** | 200 | 200 | 200 | 200 | 200 | 200 | 200 | 200 | 200 | 200 | 270 |
| **µL sample** | - | 100 from 3 | 100 from 4 | 100 from 5 | 100 from 6 | 100 from 7 | 100 from 8 | 100 from 9 | 100 from 10 | 100 from 11 | 30 from 1 mM stock |

150 µL of sample-containing solutions were added to the wells. For control experiments 150 µL PBS was added and the plate was incubated at 300 rpm for 30 min. The plate was placed on the vacuum and washed with ice-cold PBS (two times 200 µL) followed by one wash with EDTA containing buffer (200 µL, 150 mM NaCl, 25 mM HEPES pH 7.4 at 4 °C, 1 mM EDTA, 0.1 % Triton™ X-100) or $Mg^{2+}$-containing buffer (200 µL, 150 mM NaCl, 25 mM HEPES pH 7.4 at 4 °C, 1 mM $MgCl_2$, 0.1 % Triton™ X-100). The bottom of the plate was dried and it was placed on the tabletop shaker at 4 °C. Cell lysate (150 µL, 0.7 mg/mL) was added and incubated at 300 rpm for 60 min. The unbound lysate was filtered off and the wells were quickly washed six times either with EDTA-containing washing buffer (200 µL, 25 mM HEPES pH 7.4 at 4 °C, 1 mM EDTA) or $Mg^{2+}$-containing washing buffer (200 µL, 25 mM HEPES pH 7.4 at 4 °C, 1 mM $MgCl_2$). The filter plate was placed on a conical 96-well plate and competing (PCP)-InsPs were added (100µL, 5 mM (PCP)-InsP in 25 mM HEPES pH 7.4 at 4 °C, 1 mM EDTA or 1 mM $MgCl_2$). For control experiments, all competing InsPs used were added (50 µL in total). The filter plate was incubated at 300 rpm for 15 min followed by centrifugation at 1000 g for 2 min and the competition was repeated once more. The resulting affinity-enriched lysates were either used for immunoblotting or proteomic sample preparation.

**In solution digestion and TMT labeling**

Eluted proteins (100 µL per sample) were denatured by the addition of MeOH-containing buffer (100 µL, 40 % MeOH, 160 mM TEAB pH 8.5) and mixed gently. The samples were reduced by the addition of TCEP (5 mM final concentration in TEAB pH 8.5) and alkylated with CAA (40 mM final concentration in TEAB pH 8.5, prepared freshly) and incubated in the dark at rt for 1 h at 1000 rpm. LysC (1:200 in 50 mM TEAB pH 8.5) was added and incubated for 2 h at 37 °C, 1,000 rpm. Trypsin (1:100 in 50 mM TEAB pH 8.5) was added and the mixture was incubated at 37 °C for 16 h at 1,000 rpm. Afterward, the solvents were removed by vacuum centrifugation for 7 h at 37 °C.

The dried samples were resolubilized in 20 µL TEAB pH 8.5 and gently pipet mixed. TMT11-plex (0.8 mg each) was dissolved in 42 µL water-free ACN and vortexed. To each sample 3.3 µL of the corresponding TMT reagent was added and the plate was incubated for 1 h at RT in the dark, 1,000 rpm on a tabletop shaker. The reaction was quenched by adding TRIS-HCl (10 µL, 1 M, pH 8.0) and incubated for 30 min at 1000 rpm at rt. Afterwards, the fractions were pooled, the wells were washed again with 20 µL ACN and the solvents were removed by vacuum centrifugation for 2 h at 37 °C. Samples were dissolved in Milli Q water, acidified with 10 % TFA, and desalted using StageTipping[4].

**Immunoblot analysis**

30 µL of affinity-enriched lysate were mixed with 10 µL 4x laemmli sample buffer and boiled at 95 °C for 10 min. The reagent was run on a 4-20 % SDS-gel and transferred to a 0.45 µM pore size PVDF membrane (Trans turbo blot, mixed MW, 7 min). The membrane was blocked for 1 h with 5 % non-fat dried milk powder in TBS-T buffer at rt. The primary antibody in TBS-T buffer was incubated for 1.5 h and washed five times for five minutes with TBS-T buffer. Subsequently, the secondary HRP-conjugated antibody in TBS-T was incubated for 1.5 h and washed five times for five minutes with TBS-T buffer. The western blot was visualized using SuperSignal™ West Femto (Thermo Fisher Scientific) and analyzed by ImageLab 6.1.0 (BioRad).

**Recombinant protein expression**

Expression was performed as previously described for GST-hDipp1[5], MBP-XPR1$^{SPX}$[6], and SYT1$^{C2B}$[7].

**Affinity enrichment with recombinant proteins**

Multiscreen® 96 well Plate, hydrophilic PVDF membrane was placed on the PlatePrep 96-well vacuum distributor and wells were washed with 70 % EtOH (50 µL) followed by two washes with PBS (200 µL, 4 °C). Streptavidin sepharose was added (80 µL) and washed with ice-cold PBS (three times 200 µL). The bottom was dried with a clean wipe and the plate was positioned on a tabletop shaker at 4 °C. 20 µL of a 1 mM b-(PCP)-InsP solution was dissolved in 150 µL PBS and added to the plate. For control experiments 150 µL PBS was added and the plate was incubated at 300 rpm for 30 min. The plate was placed on the vacuum and washed with ice-cold PBS (two times 200 µL) followed by one wash with ethylenediamine tetraacetic acid (EDTA) containing buffer (200 µL, 150 mM NaCl, 25 mM HEPES pH 7.4 at 4 °C, 1 mM EDTA, 0.1 % TritonTM X-100). The bottom of the plate was dried and it was placed on the tabletop shaker at 4°C. Recombinant protein (150 µL, 5 nmol) was added and incubated at 300 rpm for 60 min. The plate was placed on a 96-well plate and centrifuged (4 °C, 1000 g, 2 min) to collect the unbound proteins in the supernatant. The wells were quickly washed six times with the incubation buffer. The filter plate was placed on a conical 96 well-plate and competing (PCP)-InsPs were added (50 µL, 5 mM (PCP)-InsP in 25 mM HEPES pH 7.4 at 4 °C, 1 mM EDTA ). For control experiments, all competing InsPs used were added (50 µL in total). The filter plate was incubated at 300 rpm for 15 min followed by centrifugation at 1000 g for 2 min and the competition was repeated once more. The resulting affinity-enriched lysates were used for SDS-page analysis. 30 µL of affinity-enriched lysate was mixed with 10 µL 4x laemmli sample buffer and boiled at 95 °C for 10 min. The reagent was run on a 4-20 % SDS-gel and visualized by InstantBlue® Coomassie Protein Stain (abcam).

**Pyrophosphosproteomics sample preparation**

The pyrophosphoproteomics sample preparation workflow was performed as described by Morgan *et al.*[8].

**Liquid chromatography and mass spectrometry**

Desalted peptides were resuspended in 1 % ACN with 0.05 % TFA and 1 µg was injected into a Thermo Scientific™ Dionex™ UltiMate™ 3000 system connected to a PepMap C-18 trap-column (0.075 mm x 50 mm, 3 µm particle size, 100 Å pore size, Thermo Fisher Scientific) followed by an in-house packed C18 column for reverse phase separation (Poroshell 120 EC-C18, 2.7 µm, Agilent Technologies). With a flowrate of 300 nL/min, peptides were separated using a 117 min gradient with increasing ACN concentration and analyzed on an Orbitrap Fusion Lumos mass spectrometer with FAIMS Pro™ device (Thermo Fisher Scientific) and Instrument Control Software version 4.0. MS1 and MS2 scans were acquired in the Orbitrap with a mass resolution of 120,000 and 50,000 respectively. MS1 parameters were as following: scan range m/z 400 – 1600, standard AGC target, 246 ms maximum injection time. MS2 parameters were as following: scan range first m/z 110, AGC target 1.25e5, 86 ms maximum injection time, isolation window 0.7 m/z, NCE 38 %. Previously isolated precursors were excluded from fragmentation for

60 s. Only precursors with charges +2 – +6 were subjected to MS2. Data were acquired using 2 s per column volume (CV) with an internal stepping of CVs from -50 to -65 and -85.

The LC-MS measurement for the pyrophosphoproteomics samples was performed in a similar manner as described by Morgan *et al*. with minor changes using a targeted data-dependent neutral loss triggered EThcD approach[8]. The inclusion list contained masses of expected pyrophosphorylated peptides. Fractionated pyrophosphoproteomics samples were resuspended with 50 mM medronic acid in 3 % ACN and injected in a Thermo Scientific™ Dionex™ UltiMate™ 3000 system connected to a PepMap C-18 trap-column (0.075 mm x 50 mm, 3 µm particle size, 100 Å pore size, ThermoFisher Scientific) followed by an in-house packed C18 column for reverse phase separation (Poroshell 120 EC-C18, 2.7 µm, Agilent Technologies). With a flow rate of 250 nL/min, peptides were separated using a 117 min gradient with increasing ACN concentration and analyzed on an Orbitrap Fusion mass spectrometer device (Thermo Fisher Scientific) and Instrument Control Software version 4.0. MS1 scans were acquired in the Orbitrap with a mass resolution of 120,000, the MS1 parameters were as following: scan range m/z 380 –1400, standard AGC target, 50 ms maximum injection time. Precursor ions were selected using targeted mass inclusion lists with an unscheduled time mode. Precursor ions with charge states 2 – 4 were isolated with an isolation window of 1.6 m/z and priority to the higher charge state. MS2 CID scans were acquired in the Orbitrap with a mass resolution of 15,000, the MS2 CID parameters were as following: AGC target2.5e4, 100 ms maximum injection time, NCE 25%. If neutral losses of 177.9432 Da were measured within the Top 10 most intense ions in the CID scan, an additional spectrum of the same precursor ion was acquired using EThcD. MS2 EThcD scans were acquired in the Orbitrap with a mass resolution of 120,000, the MS2 EThcD scan parameters were as following: AGC target 1e5, 2000 ms maximum injection time, normalized supplemental activation energy 30%.

**Data analysis pyrophosphoproteomics data**

Raw files were analyzed using Proteome Discoverer (ThermoFisher Scientific) version 3.0 as described by Morgan *et al.*[8]

**Mass Spectrometry Data Analysis**

Proteins have been identified and quantified from RAW files using SEQUEST and MSAmanda 2.0[9] in ProteomeDiscoverer v3.0 using the following search parameters: spectrum recalibration, MS1 mass tolerance, 10 ppm; MS2 mass tolerance, 20 ppm; maximum number of missed cleavages, 2; minimum peptide length, 6; peptide-mass, 350 – 8,000 Da. Carbamidomethylation (+57.021 Da) on cysteines was used as a static modification. Oxidation of methionines (+15.995 Da) and TMT6plex on lysines and peptide N-termini (+229.163 Da) were set as variable modifications. Data was searched against the human proteome (retrieved from Uniprot, one protein sequence per gene). The FDR has been set to 1 % on the protein level using target decoy FDR nodes. TMT quantification was achieved using the reporter ions quantifier with default settings with application of quantification value correction.

**Affinity determination using TMT abundance**

Affinity determination was performed as previously described[10-12]. In brief, the protein table for each replicate was exported from ProteomeDiscoverer. Further data processing was performed in R. Replicates were separately normalized to their maximum value. Quantification values from all three replicates were merged for individual proteins. All quantified replicates were averaged for individual proteins and the standard deviation was calculated. Dose-response curves and $K_D$s were determined using the drc R package[13]. The agreement of the dose-response curve fit with the data was assessed by calculating the Pearson correlation coefficient. Only moderate (Pearson > 0.90), good (Pearson > 0.95), and perfect hits (Pearson > 0.99) and proteins in which the first TMT channel has an abundance lower than the last channel were accepted.

**Calculation of GO Terms**

GO Terms were either calculated in R using the clusterprofiler package[14] or in Python performing a Fishers T-test calculation and the GO_biological_process_2023 table from Enrichr[15].

**CHEMICAL SYNTHESIS**

**General information**

All chemicals were purchased from the commercial suppliers VWR, Sigma Aldrich, Carl Roth, TCI, Thermo Scientific, and Roche and used without further purification. Solvents were purchased from Fisher Chemicals and dried over a 3 Å molecular sieve or dried in an MBraun-SPS-5 solvent purification system.

Silica-based flash chromatography was performed using the Combiflash Rf+™ Teledyne Isco and Redi Sep Rf disposable columns. The crude material was applied on Telos NM support from Kinesis Scientific Expert. Detection was performed at either 254 nm and 280 nm for all hexane/ethyl acetate (EtOAc)-based separation of 220 nm and 254 nm for DCM/MeOH-based separation. Silica-based purification for phosphoramidites was done using high-purity grade silica (Davisil Grade 633, pore size 60 Å, 200-425 mesh particle size) from Sigma Aldrich. Thin layer chromatography was performed using silica gel F254 plates and visualized at 254 nm or stained by potassium permanganate and heating to approximately 200 °C.

Preparative HPLC was performed using a 1260 Agilent Infinity II detector, pump, and autosampler, and a 1290 Agilent Infinity II fraction collector.

HPLC Method 1: YMC Actus Triart C18 (15 x 200 mm) column, solvent: MilliQ + 0.1 % TFA (A), acetonitrile + 0.1 % TFA (B), 35 mL/min. Gradient: 62 % B for 1 minute, followed by a gradient to 68 % B for 6 minutes and a wash at 95 % B for 2 minutes. UV detection was carried out at 220 nm.

HPLC Method 2: YMC Actus Triart C18 (15 x 200 mm) column, solvent: MilliQ + 0.1 % TFA (A), acetonitrile + 0.1 % TFA (B), 35 mL/min. Gradient: 10 % B for 3 minutes, followed by a gradient to 30 % B for 3 minutes and a wash at 95 % B for 2 minutes.

Reactions were monitored on an Agilent Infinity 1260 LC system connected to an Agilent 6130 Quadrupole. A ZOBRAX Rapid Resolution HT Narrow Bore SB-C18 column (2.1 x 50 mm) was used at 30 °C. Water with 0.1 % FA in water (A) and acetonitrile with 0.1 % FA in water (B) were used as mobile phase at a 0.7 mL/min flow rate. Gradients were chosen dependent on the polarity of the molecules either at 10 %-60 % B, 10 %-90 % B, or 40-90 % B. High resolution (HR)-MS was measured on a Thermo Fisher Q-Exactive with direct injection in pESI-FullMS positive or nESI-FullMS negative ion mode.

NMR spectra of $^{1}H$, $^{13}C$, and $^{31}P$ were recorded on a Bruker AV-600 (600 MHz) instrument at 277 K using deuterated solvents ($D_2O$, $CDCl_3$; $CD_3CN$) from Deutero. $CDCl_3$ was neutralized and stored over $K_2CO_3$ before use. Chemical shifts are depicted in ppm. For NMR-based quantification purposes, tetramethylphosphonium bromide solution in $D_2O$ of a known concentration was added and quantitative

$^{31}P$-NMR was recorded (NMR Method 1). For proton quantification, 3-(trimethylsilyl)-propionic acid-$d_4$ was dissolved in $D_2O$ and added to the compound solution and $^1H$-NMR was recorded (NMR method 1). MestreNova 10.0.2. was used for NMR data analysis.

**Synthetic precursors**

The following precursors were prepared following published procedures. **16** is commercially available. **15** was prepared according to Hostachy *et al*[16], **17** was decribed by Capolicchio *et al.*[17], **S3** was prepared following the procedure by Hager *et al.*[18], **18** was first described by Furkert *et al*[3]., and **4** was prepared using a procedure from Couto *et al*[19].

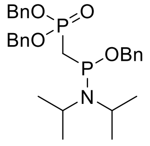


**15**

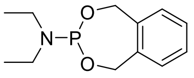


**16**

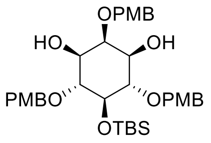


**S3**

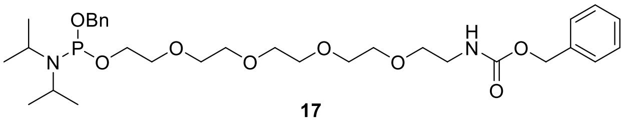


**17**

$R_1 = PO_3^{2-}$, $R_2$ = L-$NH_2$

or

$R_1$ = L-$NH_2$, $R_2 = PO_3^{2-}$

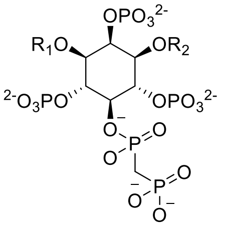


**18**

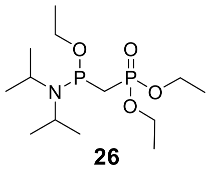


**26**

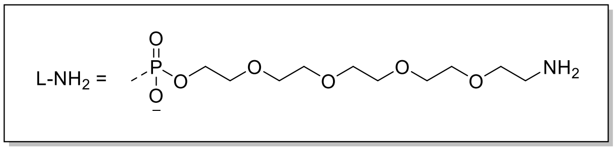


$R_1 = PO_3^{2-}$, $R_2$ = L-biotin

or

$R_1$ = L-biotin, $R_2 = PO_3^{2-}$

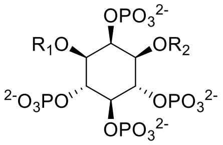


**4**

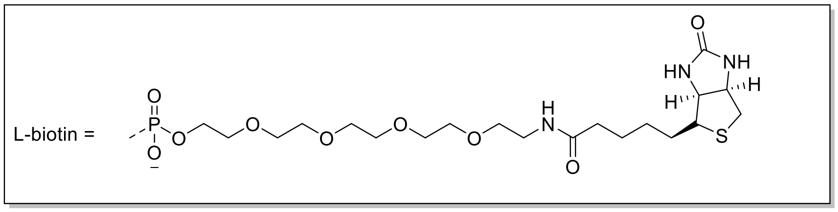

**Chemical synthesis**

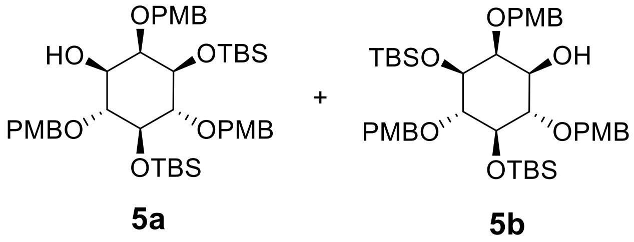

*tert*-Butyldimethylsilyl chloride (137 mg, 915 µmol, 1.2 eq.) was added to a solution of inositol **S3** (500 mg, 763 µmol, 1 eq.), and imidazole (124 mg, 1.83 mmol, 2.4 eq.) in DMF (30 mL) and stirred at 60 °C overnight. Progress was followed via LC-MS. The solution was allowed to reach room temperature before it was diluted with EtOAc (300 mL) and washed twice with LiCl (10%, 200 mL) and brine (200 mL). The organic layer was filtered through a water-repellant filter and concentrated under reduced pressure and purified by preparative HPLC (Luna 5u C8 (21.2 x 250 mm) column, solvent: MilliQ+ 0.1 % TFA (A), acetonitrile +0.1 % TFA (B), 35 mL/min. Solvent: 100% B for 5 minutes.) followed by separation of enantiomers on a (S,S) WHELK-O 1 10/100 Kromasil column (25 cm x 21.1 mm) (2.5% *i*-PrOH in *n*-hexane) to give the title compound (242 mg, 41%, 79% with reisolation of starting material (170 mg)) as a viscous oil.

$t_R$ (Enantiomer 1(**5a**)): 18.2 min

$t_R$ (Enantiomer 2(**5b**)): 26.9 min

$^1$H NMR (600 MHz, $CDCl_3$) [ppm] δ = 7.28 (m, 4H), 7.22 (m, 2H), 7, 6.94 – 6.89 (m, 2H), 6.90 – 6.84 (m, 2H), 6.85 – 6.82 (m, 2H), 4.88 (dd, *J* = 16.3, 11.3 Hz, 2H), 4.73 (t, *J* = 11.9 Hz, 2H), 4.65 (d, *J* = 11.2 Hz, 1H), 4.57 (d, *J* = 10.9 Hz, 1H). 4.12 (q, *J* = 7.2 Hz, 2H),
3.83 (s, 3H), 3.81 (s, 3H), 3.80 (s, 3H) 3.80 (m, 1H), 3.68 – 3.61 (m, 2H), 3.61 – 3.57 (m, 1H), 3.53 – 3.46 (m, 2H), 0.85 (s, 18H), 0.10 (s,3H), 0.05 (s,3H), -0.01 (s,3H), -0.09 (s,3H).
$^{13}$C NMR (151 MHz, $CDCl_3$) [ppm] δ = 171.11, 159.20, 159.16, 158.25, 131.42, 131.27, 131.05, 129.68, 129.23, 127.57, 113.81, 113.77, 113.15, 82.31, 81.85, 80.86, 75.24, 74.84, 74.81, 74.48, 72.33, 60.38, 55.29, 55.24, 55.16, 26.08, 25.95, 18.05, 18.00, 14.20, -3.93, -4.00, -4.25, -4.76.
Calculated $[M+Na]^+$: 791.3982; Measured: 791.3964.

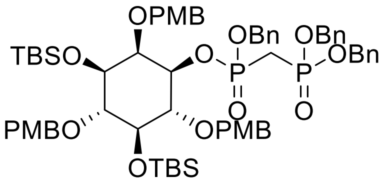

**6**

Enantiomer **5b** (68 mg, 88.4 µmol, 1 eq.) and phosphoramidite **15** (90.8 mg, 176 µmol, 2.0 eq.) were coevaporated twice with dry ACN and dried under high vacuum for 2 h. Dry $CH_2Cl_2$ (2 mL) was added and the reaction mixture was cooled in an ice bath. 5-phenyl-1H-tetrazole (90.8 mg, 176 µmol, 2.0 eq.) was added as a solid. After 5-phenyl-1*H*-tetrazole was dissolved, the ice bath was removed and the mixture was additionally stirred for 16 h. The solution was cooled in a $CO_{2(s)}$/acetone bath and *m*CPBA (77%, 48.8 mg, 217 µmol, 2.5 eq.) was added carefully: the mixture was stirred for 30 min in a $CO_{2(s)}$/acetone bath and 2 h at room temperature. EtOAc (120 mL) was added and the organic phase was washed with aq. $Na_2S_2O_3$ (100 mL), sat. aq. $NaHCO_3$ (100 mL) and brine (100 mL). The organic layer was filtered through a water-repellant filter, concentrated under reduced pressure, and purified by flash chromatography (0% to 4% MeOH in $CH_2Cl_2$) to give the title compound (51 mg, 42.6 µmol, 49%) as a colorless oil and a mixture of two diastereomers.

$^1$H NMR (600 MHz, $CD_3CN$) [ppm] δ = 7.44 – 7.14 (m, 21H), 6.86 – 6.65 (m, 6H), 5.07 – 4.49 (m, 12H), 3.73 – 3.69 (m, 6H), 3.62 (d, J = 18.2 Hz, 3H), 2.55 – 2.27 (m, 2H), 0.81 (d, J = 5.9 Hz, 6H), 0.78 (s, 3H), 0.76 (d, J = 4.2 Hz, 9H), 0.13 – 0.08 (m, 2H), 0.02 (s, 1H), -0.01 (d, J = 6.2 Hz, 2H), -0.04 (s, 1H), -0.12 (d, J = 13.4 Hz, 3H), -0.17 (d, J = 6.6 Hz, 3H).
$^{31}$P NMR (243 MHz, $CD_3CN$) [ppm] δ = 20.36, 19.86, 19.84, 19.73.
$^{13}$C NMR (151 MHz, $CD_3CN$) [ppm] δ = 160.18, 159.80, 159.44, 132.31, 131.95, 131.90, 130.23, 130.11, 129.90, 129.77, 129.66, 129.53, 129.48, 129.47, 129.41, 129.29, 129.20, 128.96, 128.90, 128.90, 128.66, 118.26, 114.56, 114.38, 114.23, 114.08, 82.65, 82.60, 81.64, 75.98, 75.94, 75.71, 75.68, 75.44, 75.29, 68.64, 68.58, 68.54, 55.86, 55.77, 55.74, 26.49 (3C), 26.34 (3C), 23.07, 18.54 (2C), -3.47, -3.87, -3.92, -4.53.
Calculated $[M+2H]^{2+}$: 599.2588; Measured: 599.2573.

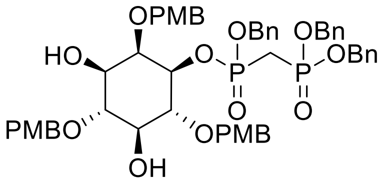

**S1**

Compound **6** (50 mg, 51.6 µmol, 1 eq.) was dissolved in dry THF (2 mL) and cooled under an ice bath. TBAF in THF (1 M, 53.9 mg, 206 µmol, 0.5 mL, 4 eq.) was added dropwise. The reaction mixture was allowed to reach room temperature and was stirred for 2 h. The reaction mixture was diluted with EtOAc (50 mL) and the organic layer was washed twice with saturated $CaCl_2$ solution (50 mL) and brine (50 mL). The organic layer was filtered through a water-repellant filter, concentrated under reduced pressure, and purified by flash chromatography (0% to 5% MeOH in $CH_2Cl_2$) to give the title compound (30 mg, 30.9 µmol, 75%) as a colorless oil and a mixture of two diastereomers.

$^1$H NMR (600 MHz, $CD_3CN$) [ppm] δ = 30 – 7.09 (m, 21H), 6.84 – 6.63 (m, 6H), 5.02 – 4.81 (m, 6H), 4.71 – 4.46 (m, 6H), 3.69 – 3.63 (m, 6H), 3.60 – 3.55 (m, 3H), 2.57 – 2.33 (m, 2H).
$^{31}$P NMR (243 MHz, $CD_3CN$) [ppm] δ = 20.55, 20.03, 19.32, 18.63.
$^{13}$C NMR (151 MHz, $CD_3CN$) [ppm] δ = 159.23, 159.17, 159.03, 136.48, 136.46, 136.36, 131.35, 131.12, 131.09, 130.96, 130.94, 130.20, 130.17, 130.16, 129.93, 129.90, 129.88, 129.74, 129.15, 129.07, 128.88, 128.85, 128.83, 128.72, 128.46, 128.00, 127.96, 127.78, 127.43, 127.40, 114.12, 114.09, 114.06, 113.06, 113.03, 113.00, 81.65, 80.68, 80.34, 79.35, 74.07, 74.04, 72.12, 71.15, 68.67, 67.62, 66.70, 56.32, 55.36, 54.41, 53.45, 25.32.
Calculated $[M+H]^+$: 969.3375; Measured: 969.3378.

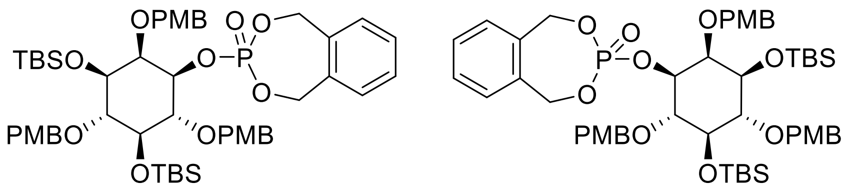


**S1a** **S1b**

Protected inositol **5a/b** (95 mg, 0.124 mmol, 1 eq.) and phosphoramidite **16** (73 mg, 0.284 mmol, 2.3 eq.) were co-evaporated twice with $CH_3CN$ (2 mL) and then dried under vacuum for 1 h, dissolved in dry $CH_2Cl_2$ (2 mL) and cooled in an ice bath, before 4,5-dicyanoimidazole (44 mg, 0.371 mmol, 3.0 eq.) was added as a solid. After 4,5-dicyanoimidazole was dissolved, the ice bath was removed and the mixture was additionally stirred for 1 h. The solution was cooled in an ice bath and *m*CPBA (100 mg, 77%; 0.433 mmol, 3.5 eq.) was added carefully: The mixture was stirred for 2.5 h at room temperature. EtOAc (20 mL) was added and the organic phase was washed with aq. $Na_2S_2O_3$ (15 mL), sat. aq. $NaHCO_3$ (10 mL) and sat. aq. NaCl (10 mL). The organic layer was dried over $Na_2SO_4$, concentrated under reduced pressure and purified by silica-based flash chromatography (0% to 40% EtOAc in hexane) to give the titled compound as a colorless oil (105 mg, 89%).

$^{1}$H NMR (600 MHz, $CDCl_3$) [ppm] δ = 7.50 – 7.08 (m, 11H), 6.95 – 6.89 (m, 2H), 6.89 – 6.82 (m, 2H), 6.77 – 6.70 (m, 2H), 5.14 (dd, J = 16.9, 13.7 Hz, 1H), 5.05 – 4.90 (m, 5H), 4.83 – 4.68 (m, 4H), 4.42 (ddd, J = 9.8, 6.9, 2.6 Hz, 1H), 3.93 – 3.87 (m, 1H), 3.67 (dd, J = 9.6, 2.3 Hz, 1H), 3.54 (t, J = 8.9 Hz, 1H), 0.87 (s, 8H), 0.84 (s, 8H), 0.15 (s, 3H), 0.03 (s, 3H), -0.00 (s, 3H), -0.08 (s, 3H).

$^{13}$C NMR (151 MHz, $CDCl_3$) [ppm] δ = 159.07, 158.64, 158.21, 135.35, 135.20, 131.47, 131.39, 130.88, 129.04, 128.92, 128.85, 128.73, 128.71, 128.68, 127.49, 113.63, 113.30, 113.13, 81.49, 80.60, 80.36, 80.31, 79.25, 79.21, 75.49, 75.19, 75.09, 74.63, 73.81, 68.42, 68.38, 68.26, 68.22, 55.28, 55.15, 55.11, 26.03, 25.93, 17.98, -3.87, -4.22, -4.83.

$^{31}$P NMR (243 MHz, $CDCl_3$) [ppm] δ = -0.43.

Calculated $[M+Na]^+$: 973.4114; Measured: 973.4092.

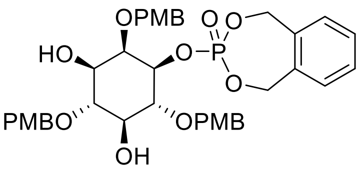


**S2a**

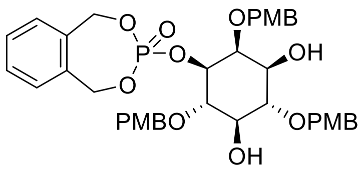


**S2b**

Compound **S1a/b** (102 mg, 0.107 mmol, 1 eq.) was dissolved in THF (8 mL) and cooled under an ice bath. TBAF 1 M in THF (0.4 mL, 4 eq.) was added dropwise. The reaction mixture was allowed to reach room temperature and was stirred for 2 h. The reaction mixture was diluted with EtOAc (50 mL) organic layer was washed aq. $CaCl_2$ and brine. The organic layer was dried over $Na_2SO_4$, concentrated under reduced pressure and purified by silica-based flash chromatography (0% to 4% MeOH in $CH_2Cl_2$) to give the title compound (70 mg, 90%) as a colorless oil.

$^{1}$H NMR (600 MHz, $CDCl_3$) [ppm] δ = 7.45 – 7.10 (m, 10H), 6.93 – 6.88 (m, 4H), 6.78 – 6.68 (m, 2H), 5.26 – 4.70 (m, 10H), 4.41 – 4.31 (m, 2H), 3.94 (t, J = 9.3 Hz, 1H), 3.83 (s, 3H), 3.81 (s, 3H), 3.72 (s, 3H), 3.66 (t, J = 9.5 Hz, 1H), 3.52 (t, J = 9.2 Hz, 1H).

$^{13}$C NMR (151 MHz, $CDCl_3$) [ppm] δ = 159.38, 159.29, 159.23, 135.46, 135.23, 130.79, 130.70, 130.49, 129.87, 129.77, 129.75, 129.49, 129.13, 129.04, 128.96, 128.93, 113.99, 113.84, 113.75, 81.14, 79.79, 79.74, 78.81, 78.58, 78.54, 75.29, 75.14, 74.83, 74.61, 71.73, 68.73, 68.69, 68.59, 68.54, 60.40, 55.27, 55.16.

$^{31}$P NMR (243 MHz,) δ -1.29.

Calculated $[M+Na]^+$: 745.2385; Measured: 745.2373.

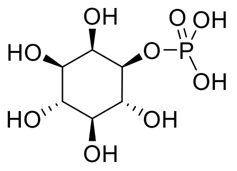

**1-InsP$_1$**

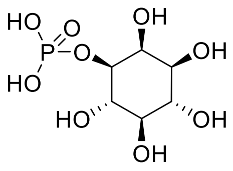

**3-InsP$_1$**

Protected inositol **S2a/b** (45 mg, 0.062 mmol, 1 eq.) was dissolved in *t*BuOH/$H_2O$ (4:1, 6 mL) and 10% Pd/C (150 mg) were added under $N_2$ atmosphere before purging the reaction vessel with $H_2$. The mixture was stirred overnight and filtered through a pad of Celite®. The residue on Celite® was additionally washed with water (4 x 4 mL). The water was filtered through a 0.2 µm nylon syringe filter and washed with $Et_2O$ once. The aqueous layer was lyophilized to yield the title product as a white solid (16 mg, quantitative). For NMR analysis cyclohexylamine (12 mg, 0.125 mmol, 2 eq.) was added.

NMR spectra are in agreement with literature[17].

Calculated $[M+H]^+$: 261.0370 Measured: 261.0370.

Ent-1 (c, 1.4, $H_2O$): $[\alpha]_D^{20}$ = -9,2 (free acid; Lit: -9.8), $[\alpha]_D^{20}$ = +3.3 (pH 10; Lit: +4.4)

Ent-2 (c, 1.6, $H_2O$): $[\alpha]_D^{20}$ = +8,1 (free acid; Lit: +9.8), $[\alpha]_D^{20}$ = -4.0 (pH 10; Lit: -4.4)

$R_1$ = P', $R_2$ = L-Cbz
or
$R_1$ = L-Cbz, $R_2$ = P'

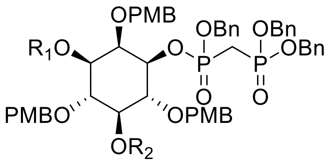


**8**

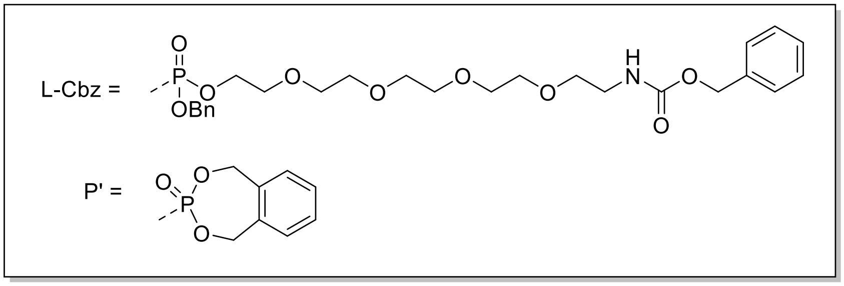


Compound **S1** (70 mg, 72.2 µmol, 1 eq.) and phosphoramidite **17** (57 mg, 93.9 µmol, 1.3 eq.) were coevaporated twice with dry ACN and dried under high vacuum for 2 h. Dry $CH_2Cl_2$ (2 mL) was added and the reaction mixture was cooled in an ice bath. 5-phenyl-1*H*-tetrazole (17 mg, 116 µmol, 1.6 eq.) was added as a solid. After 5-phenyl-1*H*-tetrazole was dissolved, the ice bath was removed and the mixture was additionally stirred for 16 h. The solution was cooled in a $CO_{2(s)}$/acetone bath and *m*CPBA (77%, 32 mg, 145 µmol, 2 eq.) was added carefully: the mixture was stirred for 30 min in a $CO_{2(s)}$/acetone bath and 2 h at room temperature. EtOAc (60 mL) was added and the organic phase was washed with aq. $Na_2S_2O_3$ (60 mL), sat. aq. $NaHCO_3$ (60 mL) and brine (60 mL). The organic layer was filtered through a water-repellant filter, concentrated under reduced pressure, and dried under high vacuum for 16 h. Phosphoramidite **16** (54 mg, 227 µmol, 5 eq.) was added and coevaporated twice with dry ACN (2 mL) and dried under high vacuum for 2 h. Dry ACN (2 mL) was added and the reaction mixture was cooled in an ice bath. 1*H*-Tetrazole (0.45 M in ACN, 17.6 mg, 250 µm, 0.56 mL, 5.5 eq.) was added, and the reaction mixture was allowed to reach room temperature and stirred overnight. The solution was cooled in a $CO_{2(s)}$/acetone bath and *m*CPBA (77%, 56 mg, 250 µmol, 5.5 eq.) was added carefully: the mixture was stirred for 30 min in a $CO_{2(s)}$/acetone bath and 2 h at room temperature. EtOAc (60 mL) was added and the organic phase was washed with aq. $Na_2S_2O_3$ (60 mL), sat. aq. $NaHCO_3$ (60 mL) and brine (60 mL). The organic layer was filtered through a water-repellant filter, concentrated under reduced pressure, and purified by flash chromatography (0% to 5% MeOH in $CH_2Cl_2$) to give the title compound (68 mg, 54.6 µmol, 56% over four steps) as a colorless oil and a mixture of multiple diastereomers and regioisomers.

$^1$H NMR (600 MHz, $CDCl_3$) [ppm] δ = 7.51 – 6.59 (m, 41H), 5.48 – 3.25 (m, 55H), 2.51 – 2.20 (m, 2H)

$^{31}$P NMR (243 MHz, $CDCl_3$) [ppm] δ = 20.80 – 18.60 (m, 2P), 1.61 - -2.51 (m, 2P).

$^{13}$C-NMR was not informative due to the formation of six diastereomers.

Calculated $[M+H]^+$: 1674.5479; Measured: 1674.5494.

$R_1$ = P', $R_2$ = L-Cbz
or
$R_1$ = L-Cbz, $R_2$ = P'

**9**

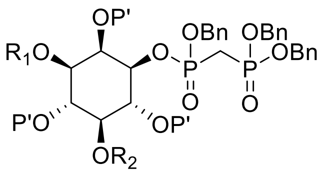

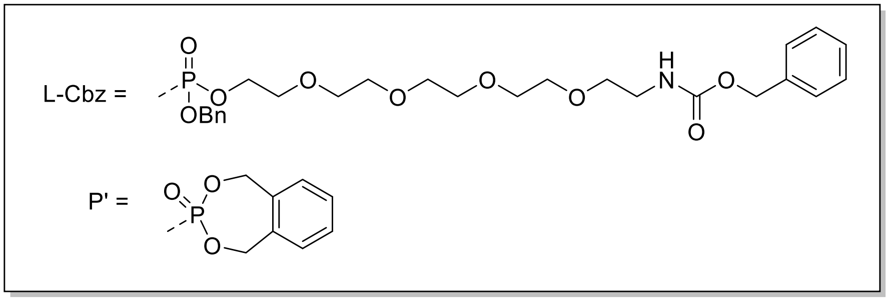


Protected inositol **8** (68 mg, 40.0 µmol, 1 eq.) was dissolved in DCM (2 mL) and cooled in an ice bath. A 20% TFA solution in $CH_2Cl_2$ (2 mL) was slowly added with a syringe. The deprotection progress was monitored *via* LC-MS. After completion, the reaction was diluted with EtOAc (100 mL) and washed twice with $Na_2PO_4$ buffer (1M, pH 7.4, 100 mL). The organic layer was filtered through a water-repellant filter, removed under reduced pressure, and used without further purification. The crude was dried under high vacuum overnight. Phosphoramidite **16** (94 mg, 395 µmol, 10 eq.) was added and the reaction mixture was coevaporated twice with dry ACN (3 mL). Dry ACN (2 mL) was added and the reaction mixture was cooled in an ice bath. 1*H*-Tetrazole (0.45 M, 30.5 mg, 435 µm, 0.97 mL, 11 eq.) was added, and the reaction mixture was allowed to reach room temperature and stirred overnight. The solution was cooled in a $CO_{2(s)}$/acetone bath and *m*CPBA (77%, 98 mg, 435 µmol, 11 eq.) was added carefully: the mixture was stirred for 30 min in a $CO_{2(s)}$/acetone bath and 2 h at room temperature. EtOAc (50 mL) was added and the organic phase was washed with aq. $Na_2S_2O_3$ (30 mL), sat. aq. $NaHCO_3$ (30 mL) and brine (30 mL). The organic layer was filtered through a water-repellant filter, concentrated under reduced pressure, and purified by flash chromatography (0% to 9% MeOH in $CH_2Cl_2$) followed by preparative HPLC (HPLC Method 1) to give the title compound (14 mg, 7.5 µM, 17% yield over two steps) as a white solid containing a mixture of the different diastereomers and regioisomers.

$^{1}$H NMR (600 MHz, $CDCl_3$) [ppm] δ = 7.64 – 6.99 (m, 41H), 5.58 – 4.89 (m, 26H), 3.66 – 3.30 (m, 16H), 2.17 – 1.85 (m, 2H).

$^{31}$P NMR (243 MHz, $CDCl_3$) [ppm] δ = 23.18 – 17.12 (m, 2P), -0.60 – -5.27 (m, 5P).

$^{13}$C-NMR was not informative due to the formation of multiple diastereomers.

Calculated $[M+2H]^{2+}$: 930.7112; Measured: 930.7110.

$R_1 = PO_3^{2-}$, $R_2$ = L-$NH_2$
or
$R_1$ = L-$NH_2$, $R_2 = PO_3^{2-}$

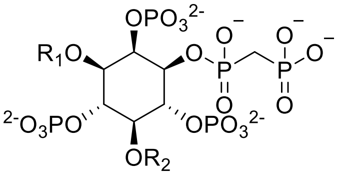

**10**

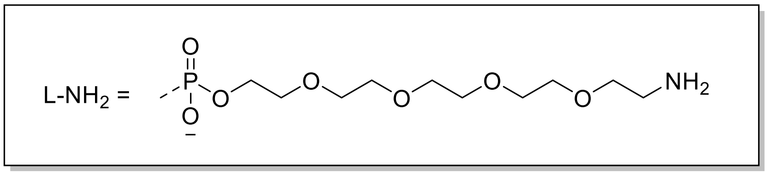


Fully protected **9** (14 mg, 7.5 µmol, 1 eq.) was dissolved in *t*BuOH/water (5 mL, 4:1), $NaHCO_3$ (5.1 mg, 60 µmol, 8 eq.) and Pd/C (10%, 40 mg, 37 µmol, 5 eq.) were added under a $N_2$ atmosphere before purging with $H_2$ gas. The mixture was stirred overnight, filtered through a Whatman® filter (0.45 µm), and lyophilized to yield the final compound as a white solid (6.1 mg, 6.3 µmol, 83%). The product was dissolved in $D_2O$ and the pH was adjusted to 6.0. The yield was determined *via* $^{31}$P-NMR spectroscopy using an internal standard (NMR method 1).

$^{1}$H NMR (600 MHz, $D_2O$ pH=6.1) [ppm] δ = 5.03 (d, J = 57.3, 9.3 Hz, 1H), 4.64 – 4.45 (m, 2H), 4.45 – 4.09 (m, 5H), 3.88 – 3.74 (m, 14H), 3.30 – 3.25 (m, 2H), 2.42 (dt, J = 78.6, 18.8 Hz, 2H).
$^{31}$P NMR (243 MHz, $D_2O$ pH=6.1) [ppm] δ = 20.77 – 17.35 (m, 1H), 16.79 – 13.82 (m, 1H), 2.81 – -2.48 (m, 5H).
$^{13}$C-NMR was not informative due to the low amount of material obtained.
Calculated $[M-2H]^{2-}$: 477.4905; Measured: 477.4909.

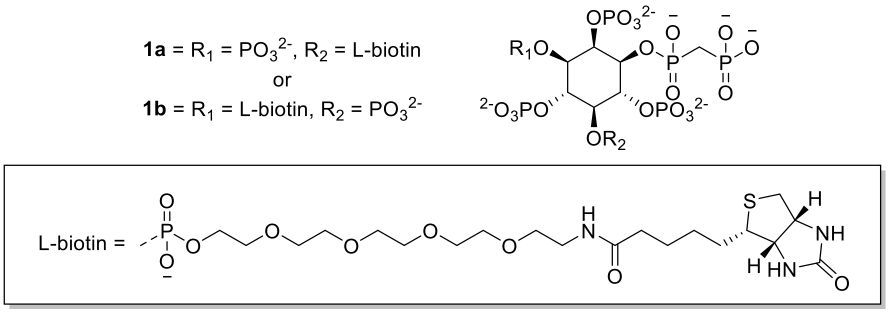


Amino-3/5***L***-1PCP-InsP$_5$ **10** (6 mg, 6.30 µmol, 1 eq.) was dissolved in 10 mL phosphate buffer ($KH_2PO_4$, 100 mM, pH 8.2). NHS-biotin (10.7 mg, 31.3 µmol, 5 eq.) was added and the reaction was stirred at 37 °C for 4 h. The precipitate was filtered off using a Whatman® filter (0.45 µm) and the aqueous phase was extracted with $Et_2O$ (2 x 30 mL) and concentrated by lyophilization. The crude was dissolved in 5 mL and pH was adjusted to 3.0. The crude was purified by preparative HPLC (HPLC method 2). Fractions containing the product were identified using an MDD assay[20], combined, and the solvent removed by lyophilization which affords the product as a white solid (6.0 mg, 5.0 µM, 81%). The product was dissolved in $D_2O$ and the pH was adjusted to 6.0. The yield was determined *via* $^1$H-NMR spectroscopy using an internal standard (NMR method 2).

$^1$H-NMR (600 MHz, $D_2O$, pH=6.1): [ppm] δ = 5.10 (d, *J* = 9.2 Hz, 1H), 4.66 (dq, *J* = 15.5, 9.3, 7.2 Hz, 4H), 4.58 – 4.37 (m, 3H), 4.32 – 4.10 (m, 2H), 3.94 – 3.58 (m, 16H), 3.43 (dt, *J* = 29.1, 4.9 Hz, 3H), 3.06 (dd, *J* = 13.1, 5.0 Hz, 1H), 2.90 – 2.79 (m, 1H), 2.57 (s, 2H), 2.34 (t, *J* = 7.3 Hz, 2H), 1.88 – 1.57 (m, 4H), 1.48 (p, *J* = 7.4 Hz, 2H).

$^{31}$P-NMR (243 MHz, $D_2O$, pH=6.1): [ppm] δ = 17.08 (3***L***, major, 80%), 14.94 (5***L***, minor, 20%), 13.19 (5***L***, minor, 20%), 11.96 (3***L***, major, 80%), -1.74, -2.16, -2.27, -2.53, -2.63, -2.73, -2.92, -3.90, -3.95.

$^{13}$C-NMR was not informative due to the low amount of material obtained.

HRMS Calculated $[M+2Na]^{2+}$: 590.5293; Measured: 590.5262.

$^1$H-$^{31}$P-HMBC, $^1$H-$^{13}$C-BIRD-HMQC-CLIP-COSY, and $^1$H-$^{13}$C-DEPT-HMBC spectra were recorded for complete characterization and analysis of the major product 3***L***-1PCP-InsP$_5$ (**Supplementary Figure S2**)

| | **$^1$H in ppm** | **$^{13}$C in ppm** | **$^{31}$P in ppm** |
|---|---|---|---|
| *myo*-Ins 1 position | 4.42 | 74.56 | 17.03 alpha |
| *myo*-Ins 2 position | 4.98 | 78.77 | -1.74 |
| *myo*-Ins 3 (with linker) | 4.26 | 76.05 | -3.95 |
| *myo*-Ins 4 position | 4.57 | 77.90 | -2.63 |
| *myo*-Ins 5 position | 4.30 | 80.37 | -3.90 |
| *myo*-Ins 6 position | 4.57 | 78.72 | -2.53 |
| PEG ($CH_2$) at 3 position | 4.23 | 67.88 | -2.92 |
| $PCH_2P$ 1 position | 2.35 | 31.28 | 17.03 alpha |

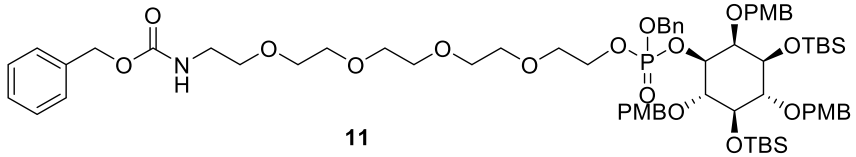

**11**

Inositol **5a** (150 mg, 195 µmol, 1 eq.) and phosphoramidite **17** (237 mg, 390 µmol, 2 eq.) were coevaporated twice with dry ACN and dried under high vacuum for 2 h. Dry ACN (1 mL) was added and the reaction mixture was cooled in an ice bath. 1*H*-Tetrazole (0.45 M, 31.4 mg, 448 µm, 0.96 mL, 2.3 eq.) was added, and the reaction mixture was allowed to reach room temperature and stirred overnight. The solution was cooled in a $CO_{2(s)}$/acetone bath and *m*CPBA (77%, 131 mg, 585 µmol, 3 eq.) was added carefully: the mixture was stirred for 30 min in a $CO_{2(s)}$/acetone bath and 2 h at room temperature. EtOAc (120 mL) was added and the organic phase was washed with aq. $Na_2S_2O_3$ (100 mL), sat. aq. $NaHCO_3$ (100 mL) and brine (100 mL). The organic layer was filtered through a water-repellant filter, concentrated under reduced pressure, and purified by flash chromatography (0% to 4% MeOH in $CH_2Cl_2$) to give the title compound (147 mg, 113 µmol, 58%) as a colorless oil and a mixture of two diastereomers.

$^1$H NMR (600 MHz, $CDCN_3$) [ppm] δ = 7.42 – 7.24 (m, 16H), 7.00 – 6.77 (m, 6H), 5.11 – 4.98 (m, 4H), 4.95 – 4.81 (m, 2H), 4.79 – 4.59 (m, 4H), 4.41 (ddd, *J* = 9.8, 7.2, 2.6 Hz, 1H), 4.35 (ddd, *J* = 9.7, 7.2, 2.6 Hz, 1H), 4.18 (dt, J = 19.3, 2.5 Hz, 1H), 4.15 – 3.98 (m, 2H), 3.87 – 3.73 (m, 9H), 3.71 – 3.46 (m, 18H), 3.27 (td, J = 5.6, 1.9 Hz, 2H), 0.88-0.84 (m, 18H), 0.17- -0.10 (m, 12H).

$^{31}$P NMR (243 MHz, $CD_3CN$) [ppm] δ = -1.63, -1.69.

$^{13}$C NMR (151 MHz, $CD_3CN$) [ppm] δ = 159.27, 158.95, 158.48, 156.38, 137.44, 136.29, 131.32, 131.27, 131.22, 130.80, 130.76, 129.26, 129.15, 129.10, 129.07, 128.59, 128.55, 128.43, 128.08, 127.91, 127.84, 127.73, 127.70, 113.66, 113.63, 113.32, 113.28, 113.12, 81.64, 80.42, 80.35, 80.30, 78.51, 78.47, 75.14, 75.08, 74.62, 74.36, 73.41, 70.16, 70.14, 70.09, 70.06, 69.92, 69.63, 69.59, 69.55, 69.50, 69.43, 69.23, 69.19, 68.96, 68.93, 67.11, 67.06, 66.91, 66.87, 65.82, 54.92, 54.85, 54.80, 54.32, 40.59, 29.18, 25.54, 25.40, 25.38, 17.60, -4.44, -4.85, -4.88, -5.50.

Calculated $[M+Na]^+$: 1314.5953 Measured: 1314.5961.

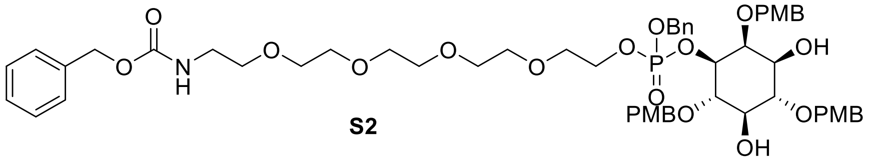

**S2**

Compound **11** (140 mg, 108 µmol, 1 eq.) was dissolved in THF (5 mL) and cooled under an ice bath. TBAF in THF (1 M, 113 mg, 433 µmol, 0.5 mL, 4 eq.) was added dropwise. The reaction mixture was allowed to reach room temperature and was stirred for 2 h. The reaction mixture was diluted with EtOAc (100 mL) and the organic layer was washed twice with saturated $CaCl_2$ solution (100 mL) and brine (100 mL). The organic layer was filtered through a water-repellant filter, concentrated under reduced pressure, and purified by flash chromatography (0% to 4% MeOH in $CH_2Cl_2$) to give the title compound (110 mg, 101 µmol, 97%) as a colorless oil and a mixture of two diastereomers.

$^{1}$H NMR (600 MHz, $CDCl_3$) [ppm] δ = 7.36-7.25 (m, 16H), 6.91-6.83 (m, 6H), 5.47 (m, 1H), 5.12-5.07(m, 4H), 4.83, 4.82-4.65(m, 6Hj), 4.29 (ddd, J = 10.3, 8.0, 2.6 Hz, 1H), 4.22 (m, 1H), 4.17-4.12 (m, 2H), 3.88-3.84 (m, 1H), 3.83-3.79 (m, 9H), 3.65-3.47 (m,18H), 3.39 (q, J = 5.4 Hz, 2H).

$^{31}$P NMR (243 MHz, $CDCl_3$) [ppm] δ = -1.49, -1.75.

$^{13}$C NMR (151 MHz, $CDCl_3$) [ppm] δ = 159.35, 159.25, 156.50, 136.66, 135.85, 130.75, 130.62, 129.73, 129.63, 129.33, 128.58, 128.48, 128.12, 128.04, 127.93, 113.94, 113.84, 113.79, 113.77, 81.00, 79.69, 78.72, 75.11, 74.83, 74.60, 71.76, 70.51, 70.46, 70.23, 70.04, 69.91, 69.34, 66.98, 66.60, 60.38, 55.27, 53.41, 40.89.

Calculated $[M+Na]^+$: 1086.4223 Measured: 1086.4214.

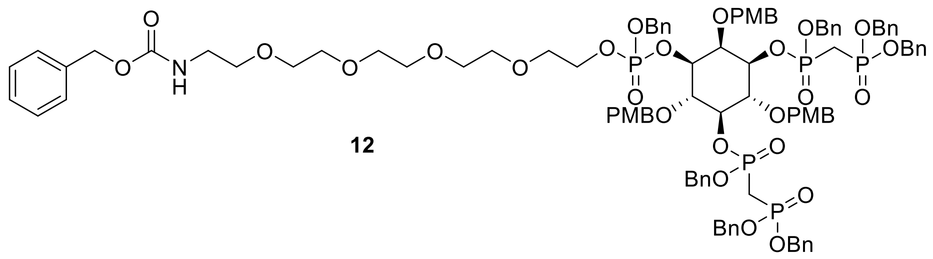

**12**

Diol **S2** (100 mg, 95.4 µmol, 1 eq.) and phosphoramidite **15** (293 mg, 572 µmol, 6.0 eq.) were coevaporated twice with dry ACN and dried under high vacuum for 2 h. Dry ACN (1 mL) was added and the reaction mixture was cooled in an ice bath. Tetrazole (0.45 M, 60.2 mg, 858 µm, 1.90 mL, 9 eq.) was added, and the reaction mixture was allowed to reach room temperature and stirred overnight. The solution was cooled in a $CO_{2(s)}$/acetone bath and *m*CPBA (77%, 171 mg, 763 µmol,8 eq.) was added carefully: the mixture was stirred for 30 min in a $CO_{2(s)}$/acetone bath and 2 h at room temperature. EtOAc (120 mL) was added and the organic phase was washed with aq. $Na_2S_2O_3$ (100 mL), sat. aq. $NaHCO_3$ (100 mL) and brine (100 mL). The organic layer was filtered through a water-repellant filter, concentrated under reduced pressure, and purified by flash chromatography (0% to 5% MeOH in $CH_2Cl_2$) to give the title compound (104 mg, 54.6 µmol, 57%) as a colorless oil and a mixture of two diastereomers.

$^1$H NMR (600 MHz, $CD_3CN$) [ppm] δ = 7.54 – 7.11 (m, 46H), 6.93 – 6.63 (m, 6H), 5.19 – 4.84 (m, 17H), 4.82 – 4.39 (m, 9H), 4.18 – 4.00 (m, 2H), 3.99 – 3.90 (m, 2H), 3.77 (dd, J = 5.9, 2.5 Hz, 2H), 3.72-3.56 (,7H), 3.56-3.45 (m, 15H), 3.25 (q, J = 5.6 Hz, 1H), 2.74 – 2.38 (m, 2H).
$^{31}$P NMR (243 MHz, $CD_3CN$) [ppm] δ = 21.50 – 18.28 (m, 4H), -1.54 – -2.43 (m, 1H).
$^{13}$C NMR (151 MHz, $CD_3CN$) [ppm] δ = 159.30, 159.06, 156.41, 137.44, 136.44, 136.39, 130.84, 130.06, 129.61, 129.49, 129.42, 129.33, 129.26, 129.20, 128.62, 128.58, 128.52, 128.49, 128.43, 128.33, 128.28, 128.25, 128.17, 128.08, 128.02, 127.98, 127.94, 127.87, 127.84, 127.73, 127.69, 113.68, 113.65, 113.59, 113.54, 113.51, 113.48, 113.44, 113.41, 78.26, 77.93, 77.65, 77.25, 77.14, 76.92, 75.30, 73.33, 70.13, 70.12, 70.05, 69.90, 69.45, 69.19, 68.70, 67.69, 67.55, 67.31, 65.83, 54.91, 54.84, 54.80, 54.77, 40.59, 25.63.
Calculated $[M+Na]^+$: 1942.6108 Measured: 1942.6110

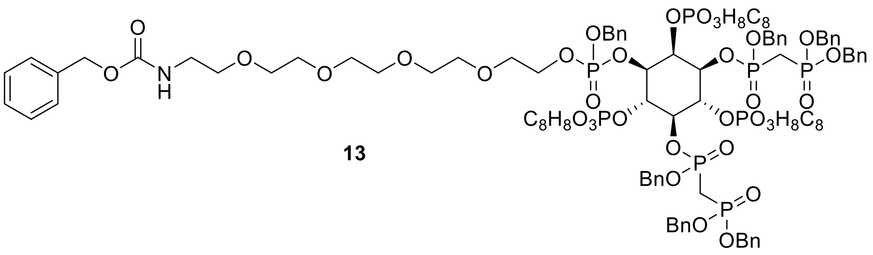

**13**

Protected inositol **12** (50 mg, 26.0 µmol, 1 eq.) was dissolved in $CH_2Cl_2$ (1 mL) and cooled in an ice bath. A 15% TFA solution in $CH_2Cl_2$ (1 mL) was added with a syringe. The deprotection progress was monitored *via* LC-MS. After completion, the reaction was diluted with EtOAc (100 mL) and washed twice with $Na_2PO_4$ buffer (1 M, pH 7.4, 100 mL). The organic layer was filtered through a water-repellant filter, removed under reduced pressure, and used without further purification. The crude was dried under a high vacuum overnight. Phosphoramidite **16** (115 mg, 480 µmol, 15 eq.) was added and the reaction mixture was coevaporated twice with dry ACN (3 mL). Dry ACN (2 mL) was added and the reaction mixture was cooled in an ice bath. Tetrazole (0.45 M, 38.0 mg, 544 µm, 1.21 mL, 17 eq.) was added, and the reaction mixture was allowed to reach room temperature and stirred overnight. The solution was cooled in a $CO_{2(s)}$/acetone bath and *m*CPBA (77%, 136 mg, 608 µmol, 19 eq.) was added carefully: the mixture was stirred for 30 min in a $CO_{2(s)}$/acetone bath and 2 h at room temperature. EtOAc (80 mL) was added and the organic phase was washed with aq. $Na_2S_2O_3$ (15 mL), sat. aq. $NaHCO_3$ (15 mL) and brine (10 mL). The organic layer was filtered through a water-repellant filter, concentrated under reduced pressure and purified by flash chromatography (0% to 9% MeOH in $CH_2Cl_2$) followed by preparative HPLC (YMC Actus Triart C18 (15 x 200 mm) column, solvent: MilliQ+ 0.1 % TFA (A), acetonitrile +0.1 % TFA (B), 35 mL/min. Gradient: 70% B for 1 minute, followed by a gradient to 95% B for 7 minutes and a wash at 100% B for 2 minutes) to give the title compound (38 mg, 71% yield over two steps) as a white solid containing a mixture of the different diastereomers.

$^{1}$H NMR (600 MHz, $CDCl_3$) [ppm] δ = 7.67 – 7.08 (m, 46H), 6.18 – 4.73 (m, 34H), 4.52 – 4.20 (m, 2H), 3.97 – 3.19 (m, 18H), 3.07 (m, 4H).

$^{31}$P NMR (243 MHz, $CDCN_3$) [ppm] δ = 19.26 – 15.99 (m, 4P), -0.92 – -7.57 (m, 4P).

$^{13}$C-NMR was not informative due to the formation of multiple diastereomers.

Calculated $[M+Na]^+$: 2128.4781 Measured: 2128.4792

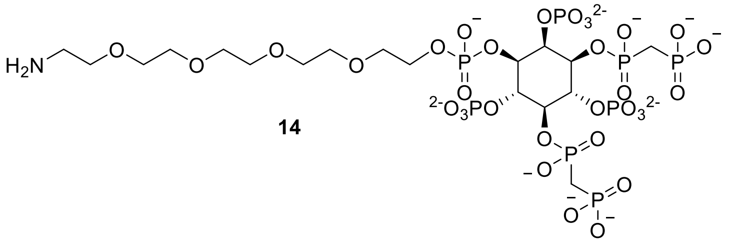

**14**

Fully protected **13** (38 mg, 18 µmol, 1 eq.) was dissolved in *t*BuOH/water (5 mL, 4:1), $NaHCO_3$ (12.2 mg, 144 µmol, 8 eq.) and Pd/C (10%, 191 mg, 180 µmol, 10 eq.) were added under a $N_2$ atmosphere before purging with $H_2$ gas. The mixture was stirred overnight, filtered through a Whatman® filter (0.45 µm), and lyophilized to yield the final compound as a white solid (18.2 mg, 98%). The product was dissolved in $D_2O$ and the pH was adjusted to 6.0. The yield was determined *via* $^{31}$P-NMR spectroscopy using an internal standard (NMR method 1).

$^{1}$H NMR (600 MHz, $D_2O$ pH=6.1) [ppm] δ = 4.70, 4.69, 4.31, 4.29, 4.22, 4.22, 4.19, 4.03, 3.95, 3.94, 3.92, 3.91, 3.49, 3.48, 2.99, 2.98, 2.98, 2.27, 2.24, 2.24, 2.21, 2.20, 2.17, 2.16, 2.14, 2.13, 2.11, 2.10.

$^{31}$P NMR (243 MHz, $D_2O$ pH=6.1) [ppm] δ = 17.70 (d, J = 11.4 Hz, 2P), 11.74 (d, J = 11.1 Hz, 1P), 11.50 (d, J = 11.5 Hz, 1P) -2.85(2P), -3.03(1P), -4.13(1P).

$^{13}$C NMR (151 MHz, $D_2O$ pH=6.1) [ppm] δ = 75.75, 75.37, 75.21, 74.25, 72.88, 71.53, 69.45, 69.40, 69.34, 69.31, 69.28, 69.11, 66.25, 64.94, 64.91, 38.75, 28.61, 27.80, 27.70, 26.98, 26.91.

$^{31}$P NMR (243 MHz, $D_2O$ pH=6.1) [ppm] δ = 17.70 (d, J = 11.4 Hz, 2P), 11.74 (d, J = 11.1 Hz, 1P), 11.50 (d, J = 11.5 Hz, 1P) -2.85(2P), -3.03(1P), -4.13(1P).

Calculated $[M-2H]^{2-}$: 516.4840 Measured: 516.4847

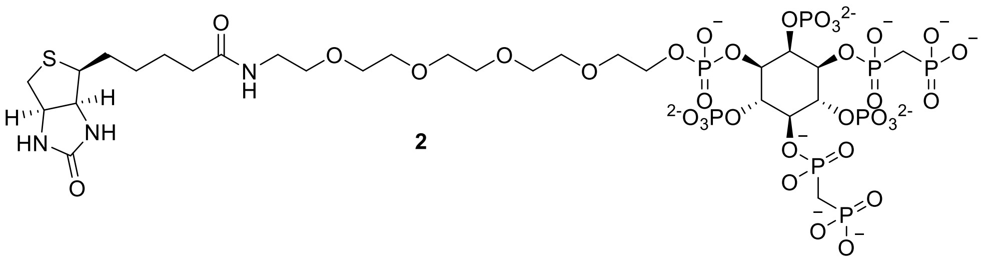

**2**

Amino-3***L***-1,5(PCP)$_2$-InsP$_4$ **14** (6 mg, 5.80 µmol, 1 eq.) was dissolved in 10 mL phosphate buffer ($KH_2PO_4$, 100 mM, pH 8.2). NHS-biotin (19.4 mg, 57.3 µmol, 10 eq.) was added and the reaction was stirred at 37 °C for 4 h. The precipitate was filtered off using a Whatman® filter (0.45 µm) and the aqueous phase was extracted with $Et_2O$ (2 x 30 mL) and concentrated by lyophilization. The crude was dissolved in 5 mL and pH was adjusted to 3.0. The crude was purified by preparative HPLC (HPLC method 2). Fractions containing the product were identified using an MDD assay[20], combined, and the solvent was removed by lyophilization which affords the product as a white solid (4.1 mg, 58%). The product was dissolved in $D_2O$ and the pH was adjusted to 6.0. The yield was determined *via* $^1$H-NMR spectroscopy using an internal standard (NMR method 2).

$^1$H-NMR (600 MHz, $D_2O$, pH=6.1): [ppm] δ = 4.95 (d, J = 9.2 Hz, 1H), 4.65 (dd, J = 8.0, 4.8 Hz, 1H), 4.59 – 4.42 (m, 3H), 4.36 (q, J = 9.7 Hz, 2H), 4.21 (dd, J = 12.0, 6.0 Hz, 3H), 3.82 – 3.73 (m, 14H), 3.67 (t, J = 5.3 Hz, 2H), 3.43 (t, J = 5.3 Hz, 2H), 3.38 (dd, J = 9.3, 4.9 Hz, 1H), 3.04 (dd, J = 13.1, 5.0 Hz, 1H), 2.82 (d, J = 13.0 Hz, 1H), 2.45 (t, 2H), 2.41 – 2.27 (m, 4H), 1.70 (m, J = 43.9, 30.0, 14.8, 7.4 Hz, 4H), 1.45 (m, J = 7.5 Hz, 2H).

$^{31}$P-NMR (243 MHz, $D_2O$, pH=6.1): [ppm] δ = 17.12, 16.97, 10.95, 10.94, -3.52, -3.56, -3.63, -4.68.

$^{13}$C-NMR was not informative due to the low amount of material obtained.

HRMS Calculated $[M-2H]^{2-}$: 629.5228; Measured: 629.5202.

$^1$H-$^{31}$P-HMBC, $^1$H-$^{13}$C-BIRD-HMQC-CLIP-COSY, and $^1$H-$^{13}$C-DEPT-HMBC spectra were recorded for complete characterization and analysis (**Supplementary Figure S3**).

| | **$^1$H in ppm** | **$^{13}$C in ppm** | **$^{31}$P in ppm** |
|---|---|---|---|
| *myo*-Ins 1 position | 4.43 | 74.56 | 16.97 alpha |
| *myo*-Ins 2 position | 4.91 | 78.70 | -4.68 |
| *myo*-Ins 3 (with linker) | 3.27 | 76.17 | -3.63 |
| *myo*-Ins 4 position | 4.56 | 77.32 | -3.56 |
| *myo*-Ins 5 position | 4.40 | 78.23 | 17.12 alpha |
| *myo*-Ins 6 position | 4.56 | 78.06 | -3.52 |
| PEG ($CH_2$) at 3 position | 4.22 | 67.93 | -3.63 |
| $PCH_2P$ 5 position | 2.49 | 31.34 | 17.12 alpha |
| $PCH_2P$ 1 position | 2.39 | 31.29 | 16.97 alpha |

**3a** = $R_1$ = $PO_3^{2-}$, $R_2$ = L-biotin
or
**3b** = $R_1$ = L-biotin, $R_2$ = $PO_3^{2-}$

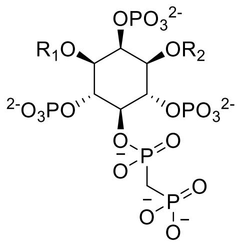


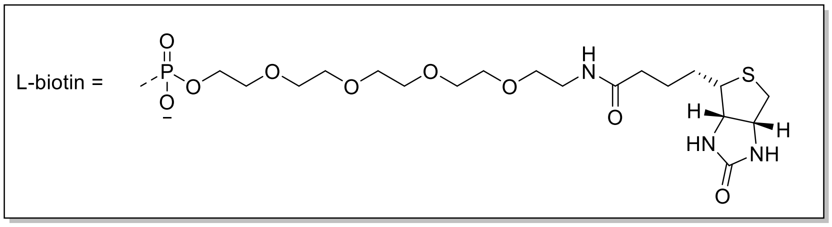


Amino-1/3***L***-5PCP-InsP5 **18** (9.5 mg, 9.9 µmol, 1 eq.) was prepared according to a published procedure[3] and was dissolved in 10 mL phosphate buffer ($KH_2PO_4$, 100 mM, pH 8.2). NHS-biotin (16.9 mg, 49.6 µmol, 5 eq.) was added and the reaction was stirred at 37 °C for 4 h. The precipitate was filtered using a Whatman® filter (0.45 µm) and the aqueous phase was extracted with $Et_2O$ (2 x 30 mL) and concentrated by lyophilization. The crude was dissolved in 5 mL and pH was adjusted to 3.0. The crude was purified by preparative HPLC (HPLC method 2). Fractions containing the product were identified using an MDD assay[20], combined, and the solvent was removed by lyophilization which afforded the product as a white solid (6.0 mg, 51 %). The product was dissolved in $D_2O$ and the pH was adjusted to 6.1. The yield was determined *via* $^{31}$P-NMR spectroscopy and $^{1}$H-NMR spectroscopy using an internal standard (NMR methods 1 and 2).

$^{1}$H-NMR (600 MHz, $D_2O$, pH=6.1): [ppm] δ = 4.95 (dt, *J* = 9.6, 2.5 Hz, 1H), 4.63 – 4.52 (m, 3H), 4.46 (q, *J* = 9.4 Hz, 1H), 4.38 (td, *J* = 7.9, 3.5 Hz, 3H), 4.21 – 4.03 (m, 2H), 3.75 – 3.60 (m, 14H), 3.57 (t, *J* = 5.3 Hz, 2H), 3.33 (t, *J* = 5.3 Hz, 2H), 3.31 – 3.25 (m, 1H), 2.94 (dd, *J* = 13.1, 5.0 Hz, 1H), 2.73 (d, *J* = 13.1 Hz, 1H), 2.58 (t, *J* = 20.3 Hz, 2H), 2.22 (t, *J* = 7.3 Hz, 2H), 1.72 – 1.47 (m, 4H), 1.41 – 1.30 (m, 2H).

$^{31}$P-NMR (243 MHz, $D_2O$, pH=6.1): [ppm] δ = 19.41, 18.74, 033, −0.01, −0.21, −1.48.

$^{13}$C-NMR (151 MHz, $D_2O$, pH=6.1): [ppm] δ = 177.00, 165.35, 76.04, 73.12, 70.02, 69.65.

HRMS Calculated $[M-2H]^{2-}$: 590.5293; Measured: 590.5264.

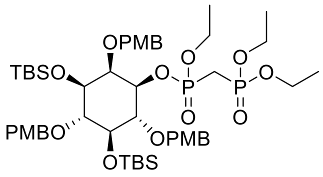

**20**

Enantiomer **5b** (50 mg, 65.0 µmol, 1 eq.) and phosphoramidite **26** (42.6 mg, 130 µmol, 2.0 eq.) were coevaporated twice with dry ACN and dried under high vacuum for 2 h. Dry ACN (2 mL) was added and the reaction mixture was cooled in an ice bath. 1*H*-Tetrazole (0.45 M, 18.2 mg, 260 µm, 0.57 mL, 4 eq.) was added, and the reaction mixture was allowed to reach room temperature and stirred overnight. The solution was cooled in a $CO_{2(s)}$/acetone bath and *m*CPBA (77%, 33.5 mg, 149.5 µmol, 2.3 eq.) was added carefully: the mixture was stirred for 30 min in a $CO_{2(s)}$/acetone bath, and 2 h at room temperature. EtOAc (120 mL) was added and the organic phase was washed with aq. $Na_2S_2O_3$ (100 mL), sat. aq. $NaHCO_3$ (100 mL) and brine (100 mL). The organic layer was filtered through a water-repellant filter, concentrated under reduced pressure, and purified by flash chromatography (0% to 3% MeOH in $CH_2Cl_2$) to give the title compound (40 mg, 39.5 µmol, 61%) as a colorless oil and a mixture of two diastereomers.

$^1$H NMR (600 MHz, $CDCl_3$) [ppm] δ = 7.42 – 7.32 (m, 3H), 7.32 – 7.26 (m, 2H), 7.23 (d, *J* = 8.3 Hz, 1H), 6.95 – 6.91 (m, 2H), 6.89 – 6.82 (m, 4H), 4.98 – 4.88 (m, 2H), 4.83 – 4.62 (m, 4H), 4.34 – 3.99 (m, 8H), 3.89 – 3.79 (m, 10H), 3.76 – 3.62 (m, 2H), 3.54 (s, 1H), 2.45 – 2.04 (m, 2H), 1.36 – 1.26 (m, 9H), 0.86 (d, *J* = 2.7 Hz, 9H), 0.82 (d, *J* = 1.9 Hz, 9H), 0.15 (d, *J* = 5.7 Hz, 3H), 0.04 (s, 3H), 0.02 (d, *J* = 4.6 Hz, 3H), -0.09 (s, 3H).

$^{31}$P NMR (243 MHz, $CDCl_3$) [ppm] δ = 20.75, 19.31, 19.27, 18.93.

$^{13}$C NMR (151 MHz, $CDCl_3$) [ppm] δ = 159.00, 158.62, 158.10, 131.32, 131.00, 130.95, 128.95, 128.80, 128.56, 128.43, 128.24, 127.84, 127.38, 113.58, 113.55, 113.46, 113.29, 113.04, 81.48, 81.16, 80.77, 80.39, 80.24, 75.13, 74.72, 74.53, 73.86, 63.43, 62.88, 62.65, 62.52, 62.34, 62.24, 55.20, 55.07, 46.58, 25.92 (3C), 25.84 (3C), 17.89, 16.23, 16.19, -3.95, -3.99, -4.29, -4.90.

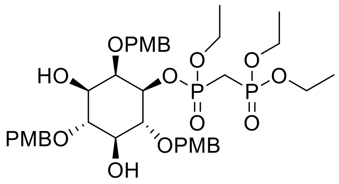

**21**

Compound **20** (327 mg, 323 µmol, 1 eq.) was dissolved in dry THF (18 mL) and cooled under an ice bath. TBAF in THF (1 M, 338 mg, 1.29 mmol, 1.29 mL, 4 eq.) was added dropwise. The reaction mixture was allowed to reach room temperature and was stirred for 2 h. The deprotection progress was monitored *via* LC-MS. The reaction mixture was diluted with EtOAc (150 mL) and the organic layer was washed twice with saturated $CaCl_2$ solution (150 mL) and brine (150 mL). The organic layer was filtered through a water-repellant filter, concentrated under reduced pressure, and purified by flash chromatography (0% to 4% MeOH in $CH_2Cl_2$) to give the title compound (198 mg, 252 µmol, 78%) as a colorless oil and a mixture of two diastereomers.

$^{1}$H NMR (600 MHz, $CDCl_3$) [ppm] δ = 7.29 (dddd, *J* = 15.0, 8.9, 5.8, 3.4 Hz, 6H), 6.98 – 6.80 (m, 6H), 4.88 – 4.53 (m, 6H), 4.41 (dtd, *J* = 22.5, 9.6, 2.7 Hz, 1H) 4.30 – 4.00 (m, 7H), 3.86 – 3.78 (m, 10H), 3.72 – 3.41 (m, 3H), 2.49 – 2.28 (m, 2H), 1.34 – 1.25 (m, 9H).
$^{31}$P NMR (243 MHz $CDCl_3$) [ppm] δ = 20.76, 19.56, 19.15, 19.07.
$^{13}$C NMR (151 MHz, $CDCl_3$) [ppm] δ = 159.41, 159.36, 159.31, 130.63, 130.53, 130.50, 130.04, 129.77, 129.71, 129.67, 129.50, 129.47, 129.33, 114.14, 114.01, 113.96, 113.92, 113.84, 83.85, 81.17, 80.78, 79.89, 79.85, 79.74, 74.59, 74.37, 74.26, 71.64, 63.46, 63.22, 63.10, 62.76, 62.45, 55.27, 46.29, 16.33, 16.29, 16.20.

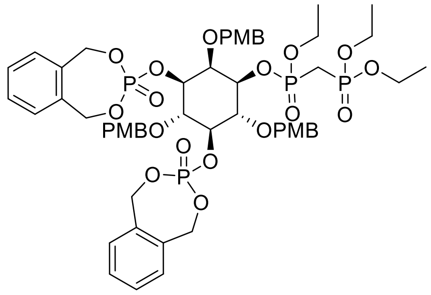

**22**

Protected inositol **21** (200 mg, 255 µmol, 1 eq.) and phosphoramidite **16** (244 mg, 1.02 mmol, 4 eq.) were coevaporated twice with dry ACN (3 mL). Dry ACN (8 mL) was added and the reaction mixture was cooled in an ice bath. 1*H*-Tetrazole (0.45 M, 78.8 mg, 1.12 mm, 2.50 mL, 4.4 eq.) was added, and the reaction mixture was allowed to reach room temperature and stirred overnight. The solution was cooled in a $CO_{2(s)}$/acetone bath and *m*CPBA (77%, 257 mg, 1.15 mmol, 4.5 eq.) in $CH_2Cl_2$ (10 mL) was added carefully: the mixture was stirred for 30 min in a $CO_{2(s)}$/acetone bath and 2 h at room temperature. EtOAc (150 mL) was added and the organic phase was washed with aq. $Na_2S_2O_3$ (130 mL), sat. aq. $NaHCO_3$ (130 mL) and brine (130 mL). The organic layer was filtered through a water-repellant filter, concentrated under reduced pressure, and purified by flash chromatography (0% to 10% MeOH in $CH_2Cl_2$) followed by preparative HPLC (HPLC Method 1) to give the title compound (211 mg, 183 µM, 72% yield over two steps) as a white solid containing a mixture of six diastereomers.

$^{1}$H NMR (600 MHz, $CDCl_3$) [ppm] δ = 7.43 – 7.26 (m, 10H), 7.13 (ddq, *J* = 8.7, 5.5, 3.4, 2.9 Hz, 4H), 6.93 – 6.81 (m, 4H), 6.71 – 6.65 (m, 2H), 5.34 – 4.47 (m, 16H), 4.47 – 4.28 (m, 2H), 4.28 – 3.91 (m, 8H), 3.83 – 3.77 (m, 6H), 3.70 (d, *J* = 4.7 Hz, 3H), 2.40 – 2.10 (m, 2H), 1.31 – 1.14 (m, 9H).

$^{31}$P NMR (243 MHz, $CDCl_3$) [ppm] δ = 20.57, 19.14, 18.88, 18.86, 1.38, 1.29, -1.62, -1.85.

$^{13}$C-NMR was not informative due to the formation of six diastereomers.

Calculated $[M+H]^+$: 1147.3171; Measured: 1147.3176.

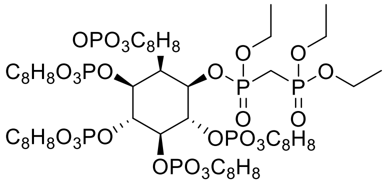

**24**

Protected inositol **22** (100 mg, 127 µmol, 1 eq.) was dissolved in $CH_2Cl_2$ (7 mL) and cooled in an ice bath. A 20% TFA solution in $CH_2Cl_2$ (7 mL) was added with a syringe. The deprotection progress was monitored *via* LC-MS. After completion, the reaction was diluted with EtOAc (100 mL) and washed twice with $Na_2PO_4$ buffer (1M, pH 7.4, 100 mL). The organic layer was filtered through a water-repellant filter, removed under reduced pressure, and used without further purification. The crude was dried under high vacuum overnight. Phosphoramidite **16** (182 mg, 762 µmol, 6 eq.) was added and the reaction mixture was coevaporated twice with dry ACN (3 mL). Dry ACN (2 mL) was added and the reaction mixture was cooled in an ice bath. 1*H*-Tetrazole (0.45 M, 57.9 mg, 862 µm, 1.84 mL, 6.5 eq.) was added, and the reaction mixture was allowed to reach room temperature and stirred overnight. The solution was cooled in a $CO_{2(s)}$/acetone bath and *m*CPBA (77%, 199 mg, 889 µmol, 7 eq.) was added carefully: the mixture was stirred for 30 min in a $CO_{2(s)}$/acetone bath and 2 h at room temperature. EtOAc (200 mL) was added and the organic phase was washed with aq. $Na_2S_2O_3$ (200 mL), sat. aq. $NaHCO_3$ (200 mL) and brine (200 mL). The organic layer was filtered through a water-repellant filter, concentrated under reduced pressure, and purified by flash chromatography (0% to 6% MeOH in $CH_2Cl_2$) followed by preparative HPLC (HPLC Method 1) to give the title compound (80 mg, 60.0 µM, 68% yield over two steps) as a white solid containing a mixture of the different diastereomers.

$^{1}$H NMR (600 MHz, $CDCl_3$) [ppm] δ = 7.45 – 7.26 (m, 20H), 5.77 – 4.97 (m, 26H), 4.45 (p, *J* = 7.2 Hz, 2H), 4.25 (ddd, *J* = 15.5, 13.4, 7.3 Hz, 4H), 2.79 (ddd, *J* = 22.8, 20.5, 15.5 Hz, 2H), 1.48 (t, *J* = 7.0 Hz, 3H), 1.41 (t, *J* = 7.1 Hz, 6H).

$^{31}$P NMR (243 MHz, $CDCl_3$) [ppm] δ = 22.60, 19.29, -2.45, -3.61, -3.69, -4.40, -4.57.

$^{13}$C-NMR was not informative due to the formation of multiple diastereomers.

Calculated $[M+Na]^+$: 1333.1844; Measured: 1333.1838.

**Appendix – NMR spectra**

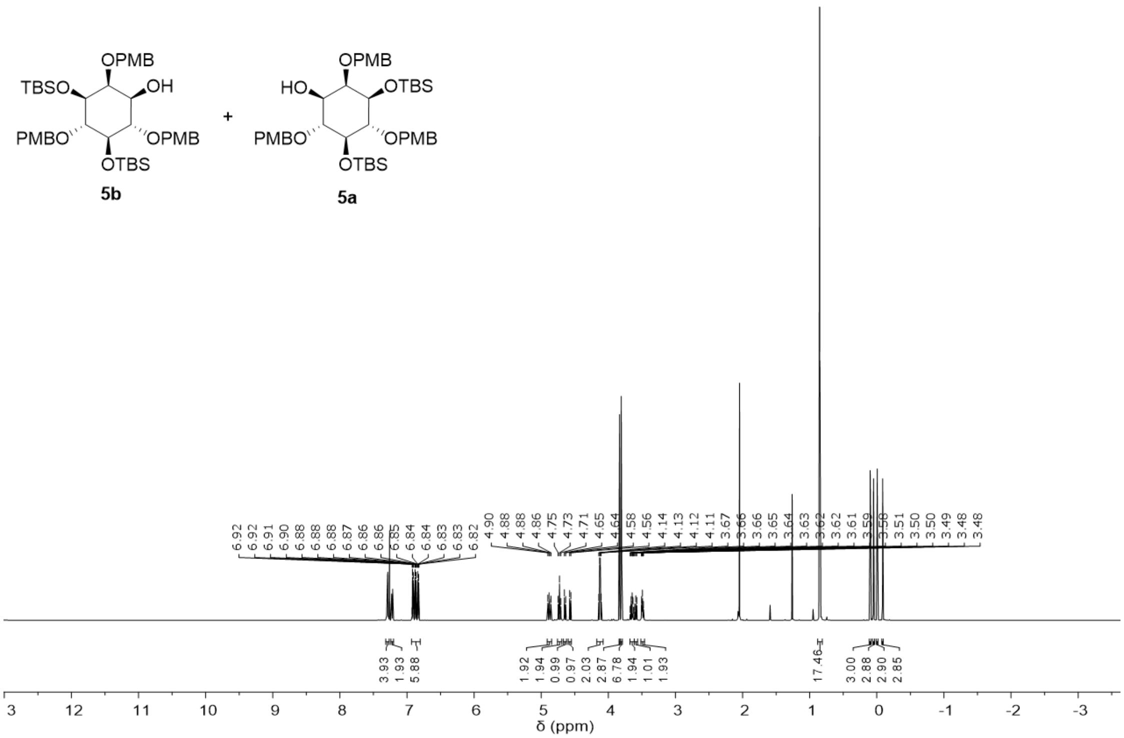


$^{1}H$ spectrum of compound **5**.

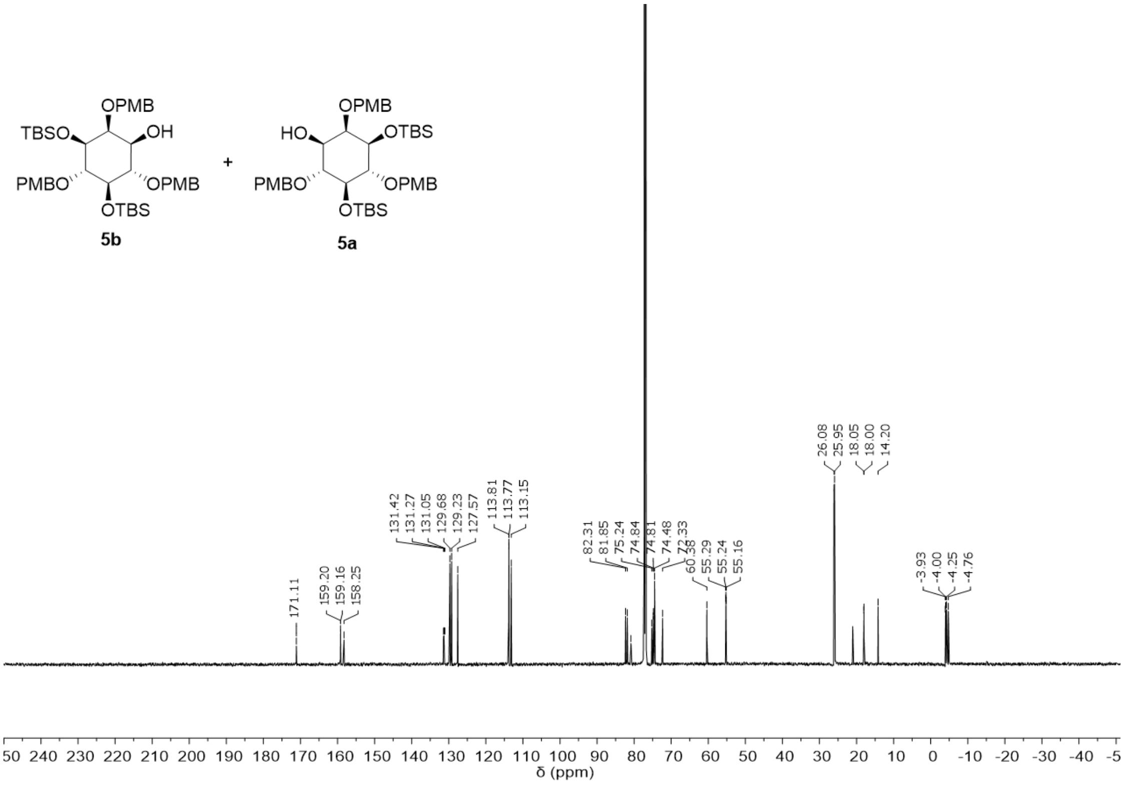


$^{13}C$ spectrum of compound **5**.

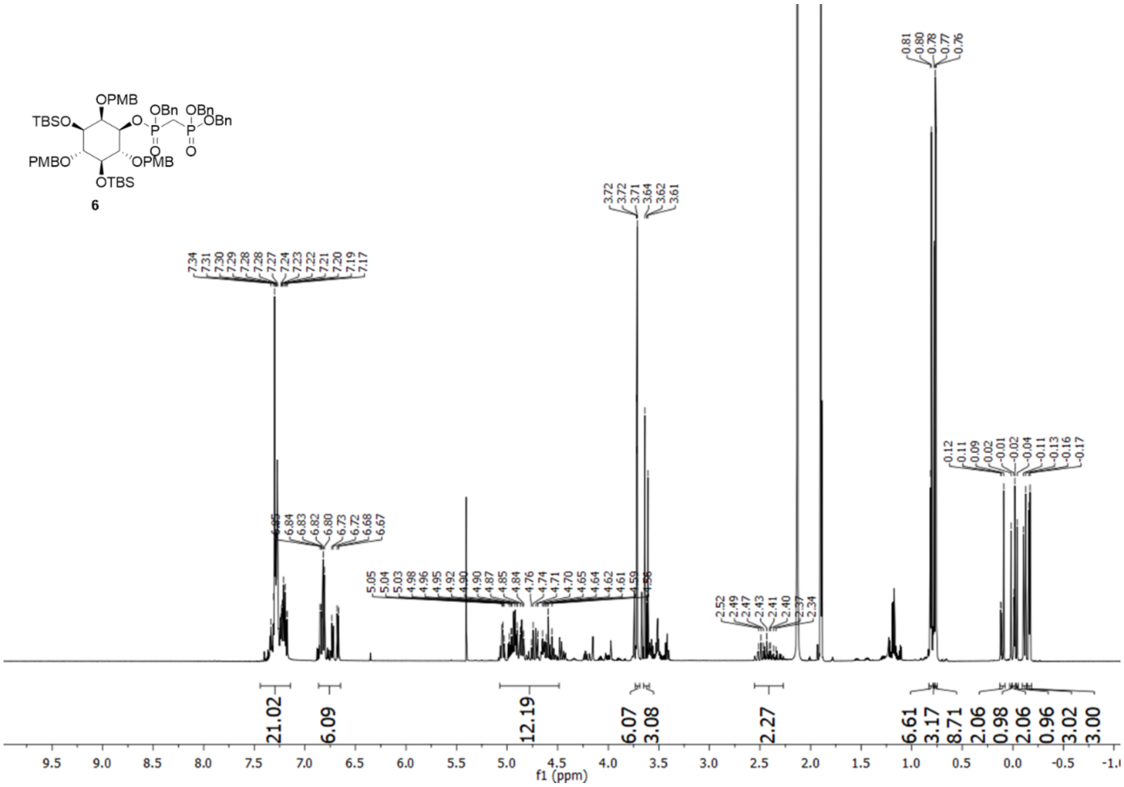


[1]H spectrum of compound **6**.

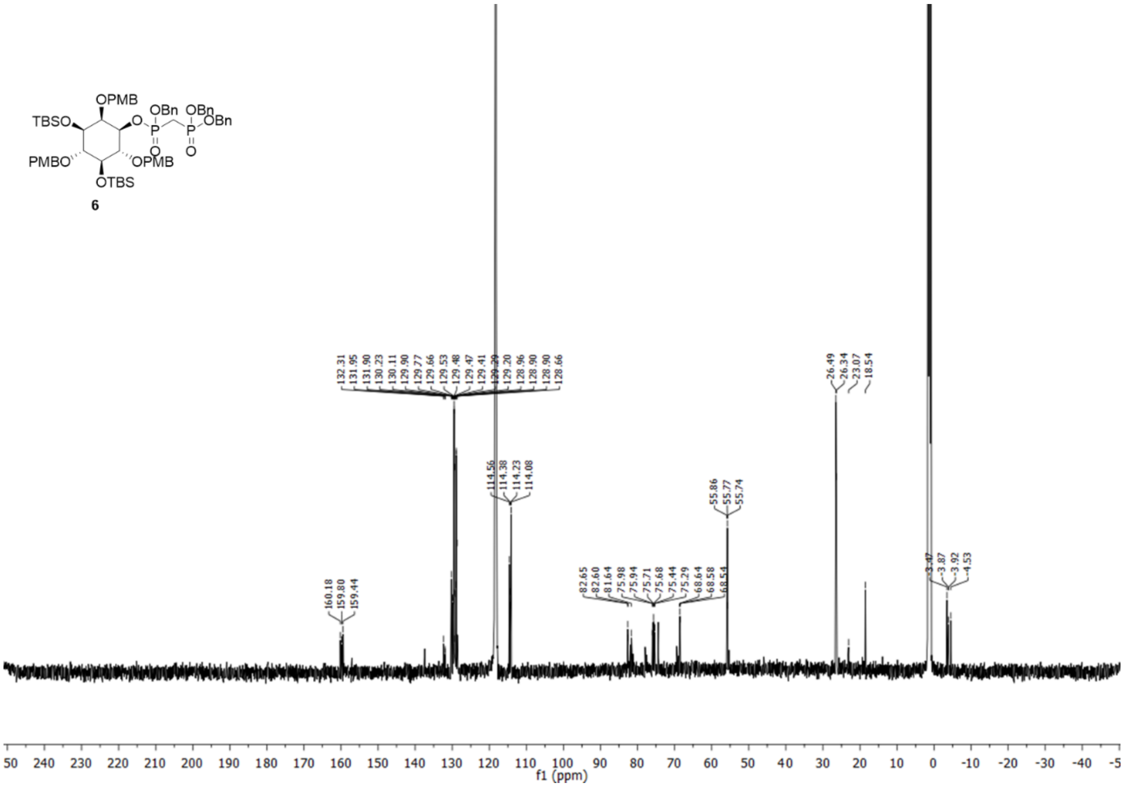


[13]C spectrum of compound **6**.

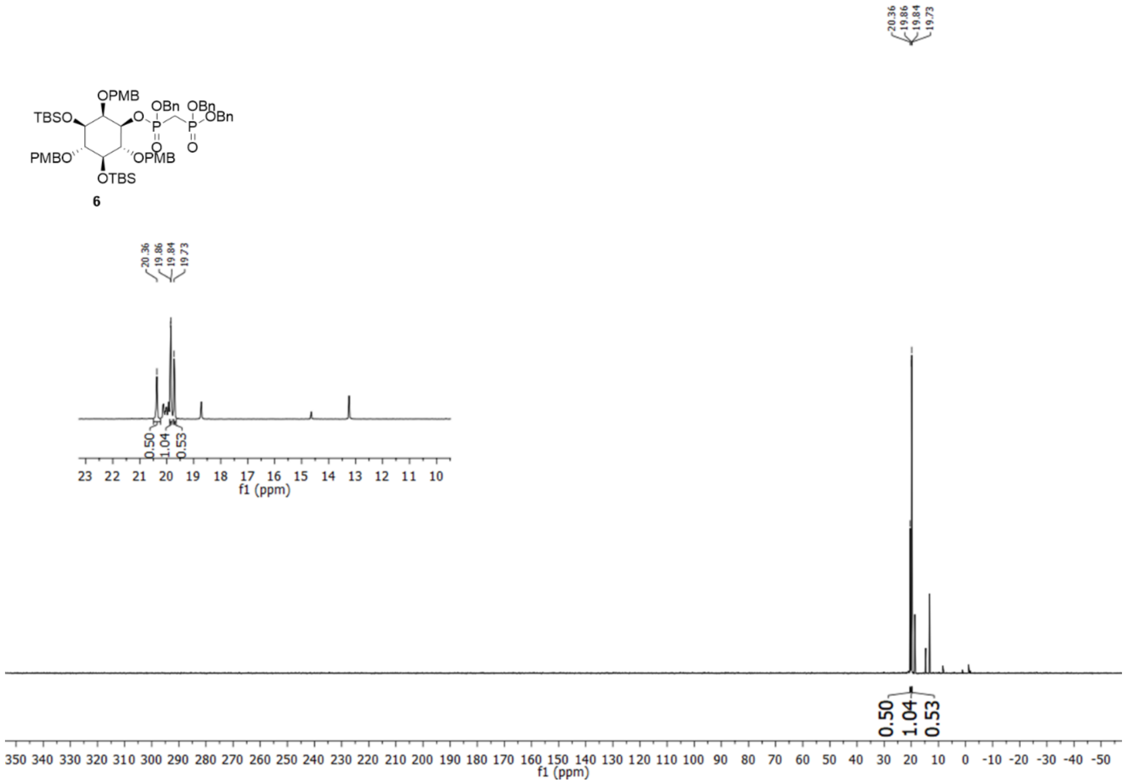

$^{31}$P spectrum of compound **6**.

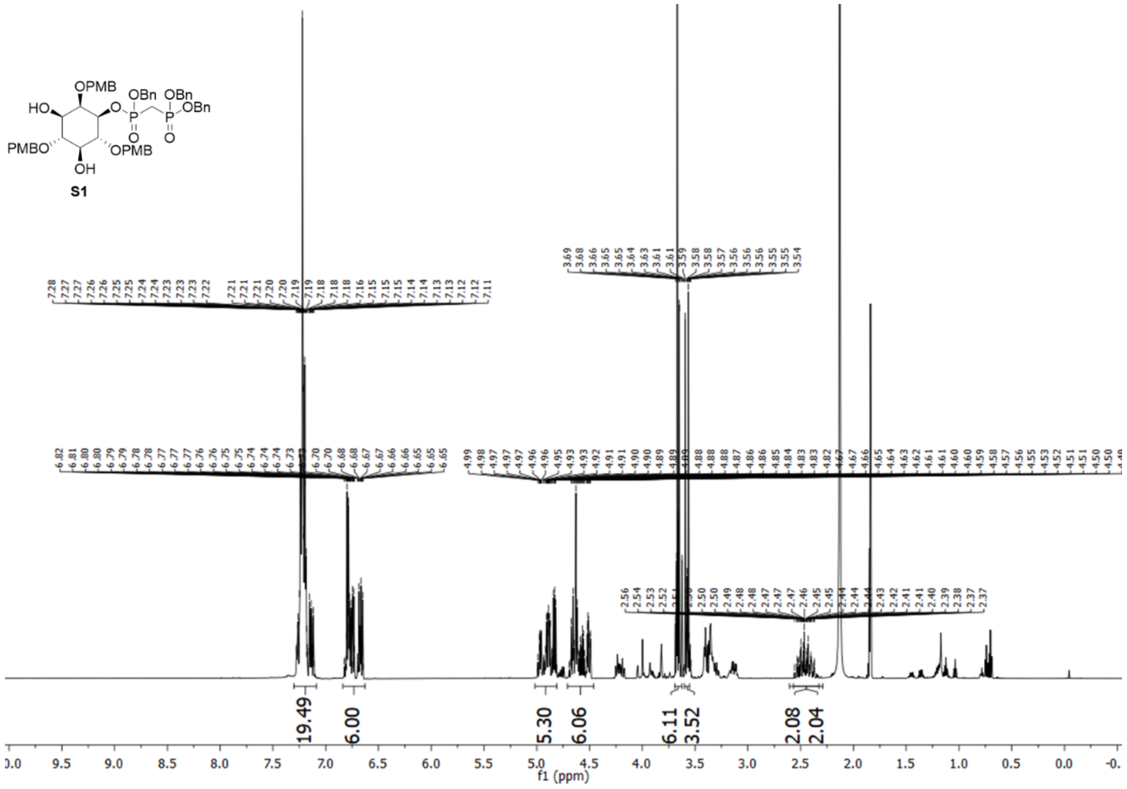

$^{1}$H spectrum of compound **S1**.

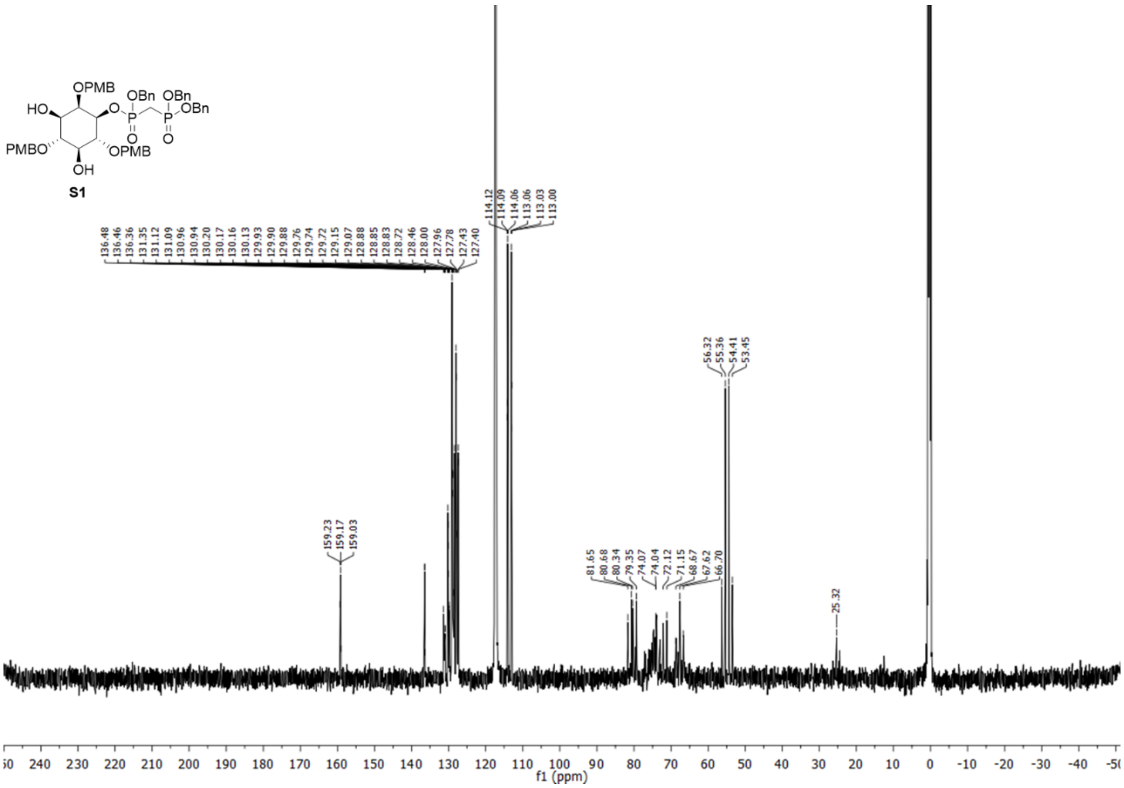


[13] C spectrum of compound **S1**.

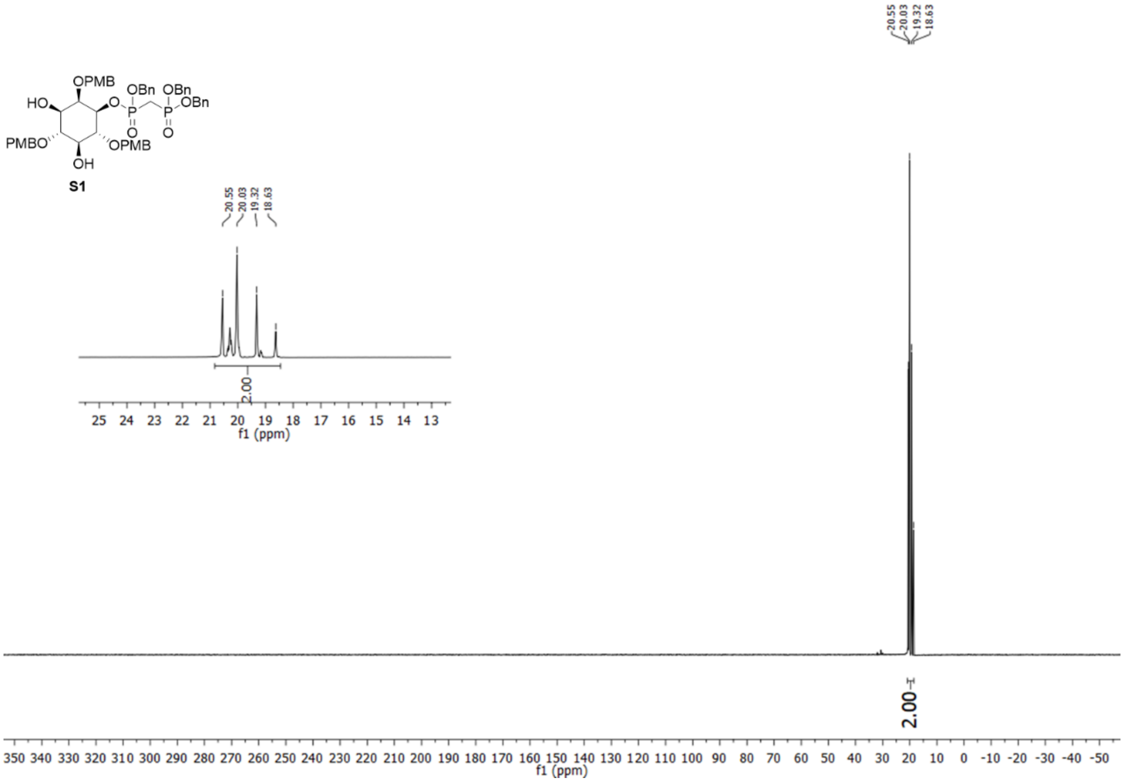


[31] P spectrum of compound **S1**.

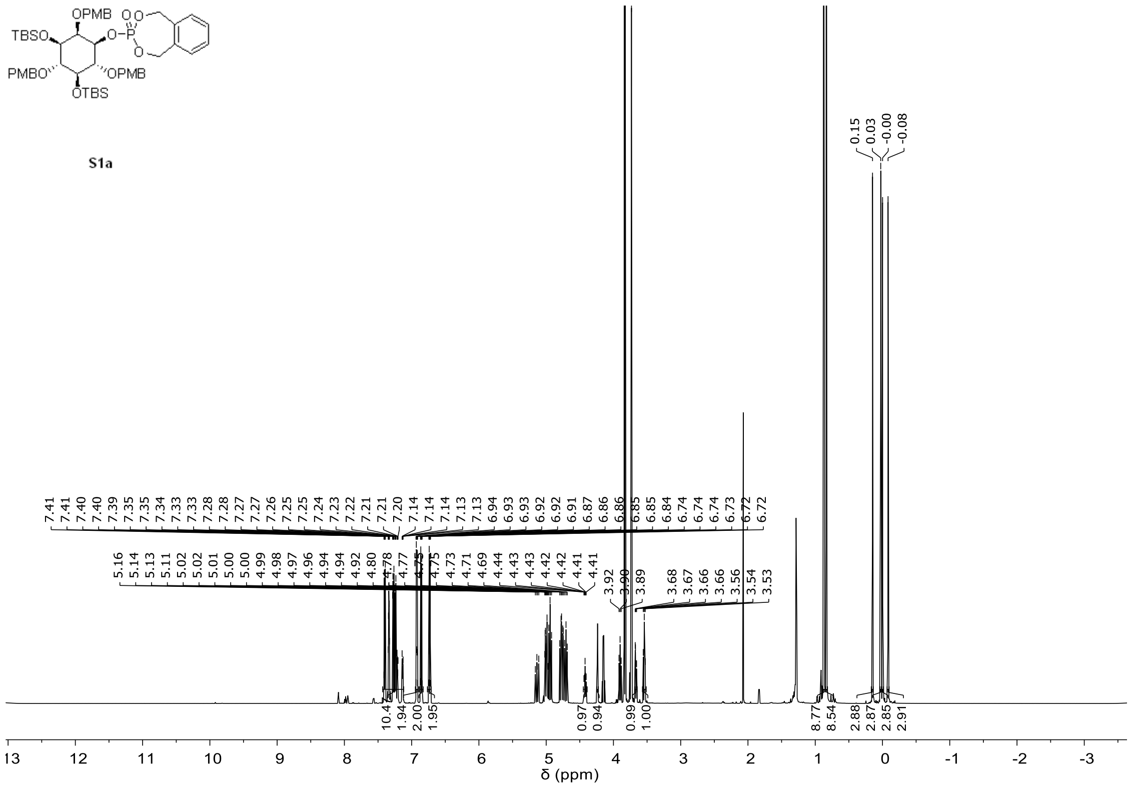


[1]H spectrum of compound **S1a**.

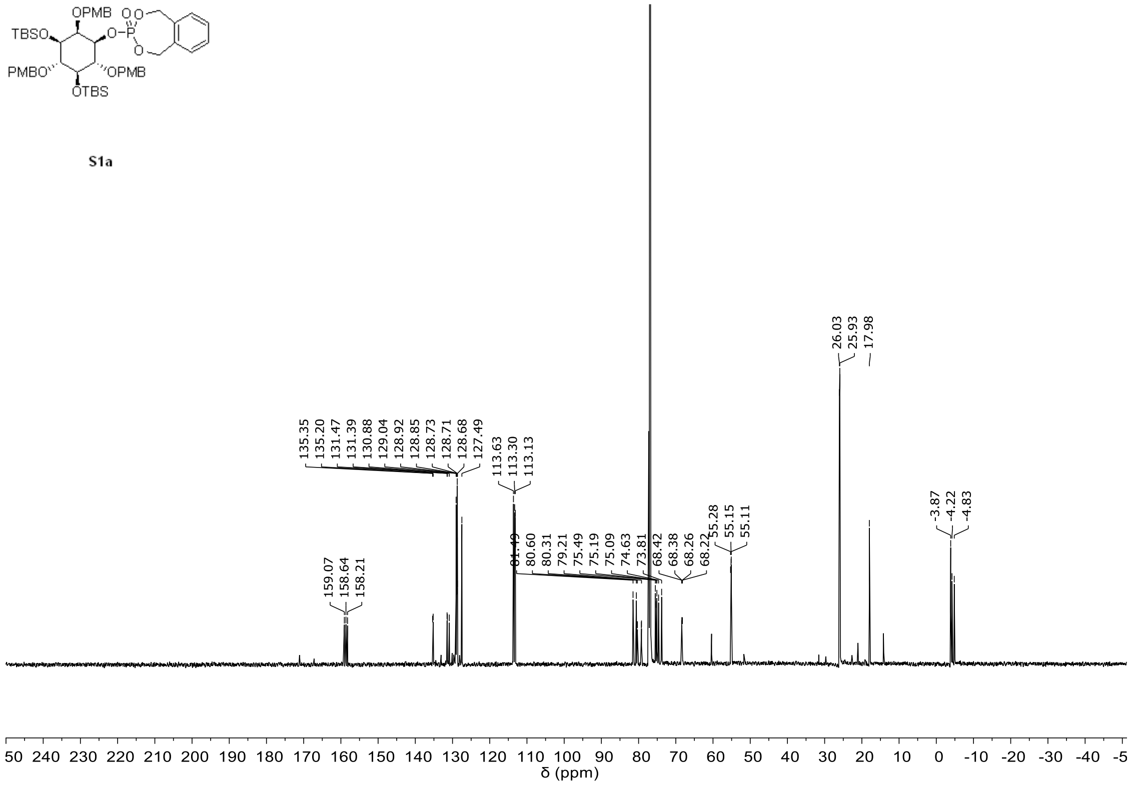


[13]C spectrum of compound **S1a**.

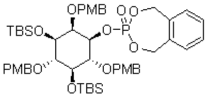


**S1a**

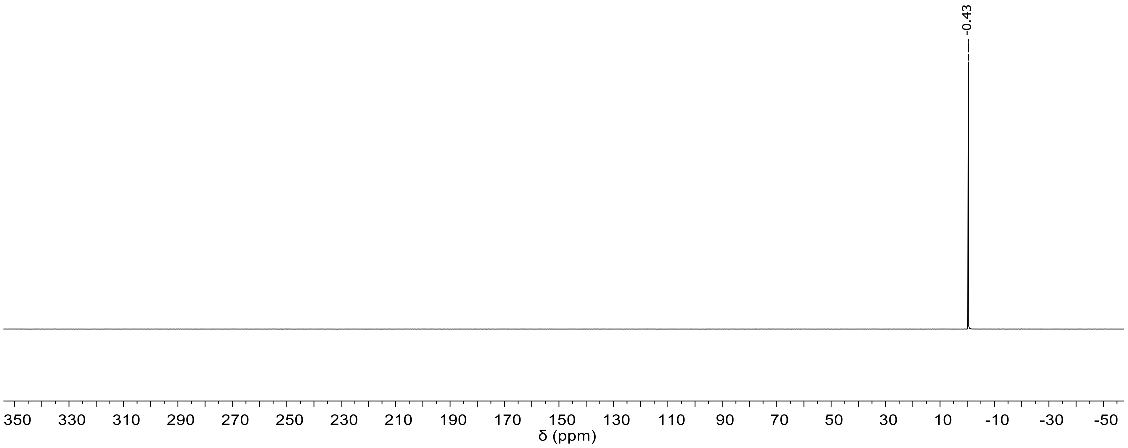


$^{31}P$ spectrum of compound **S1a**.

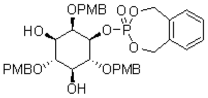


**S2a**

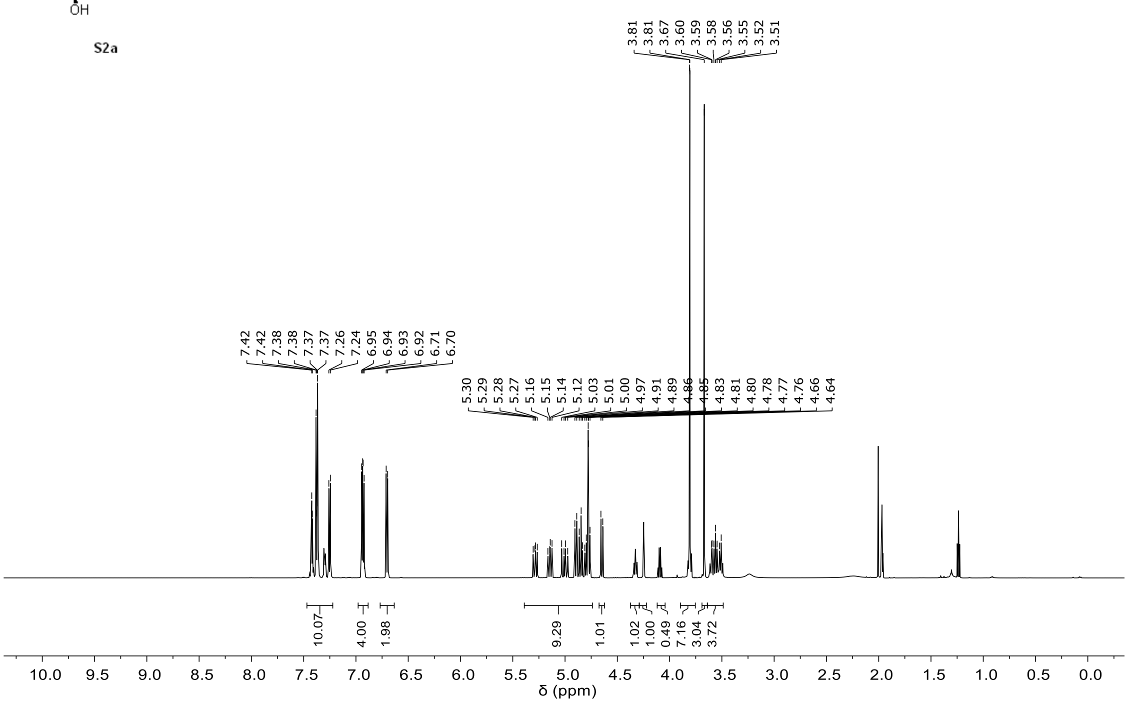


$^{1}H$ spectrum of compound **S2a**.

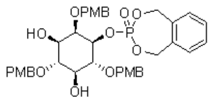

**S2a**

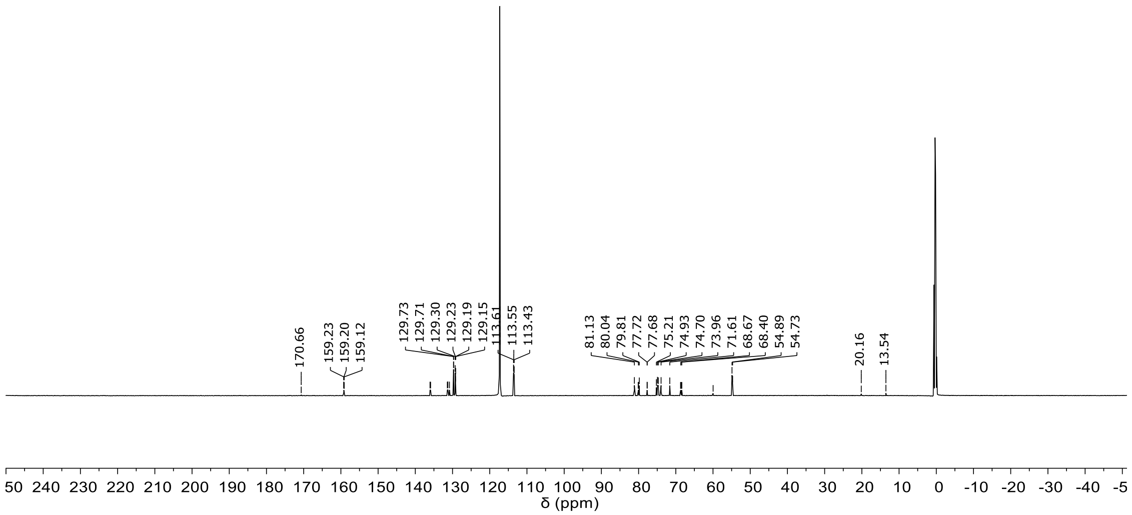


$^{13}C$ spectrum of compound **S2a**.

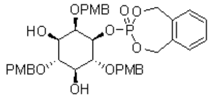

**S2a**

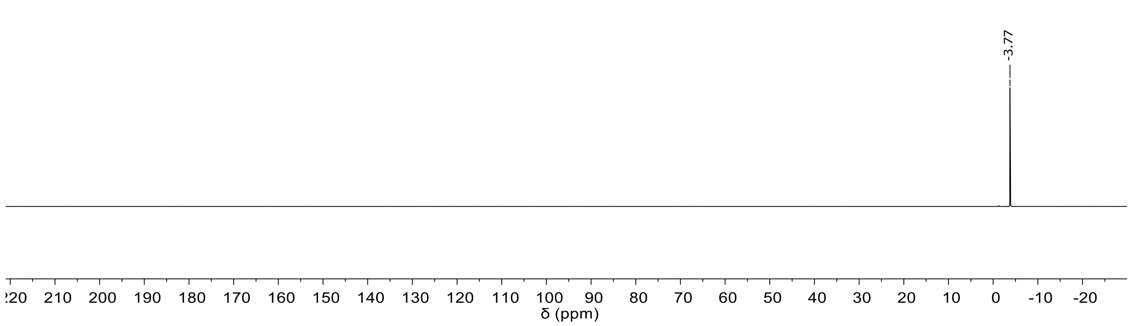


$^{31}P$ spectrum of compound **S2a**.

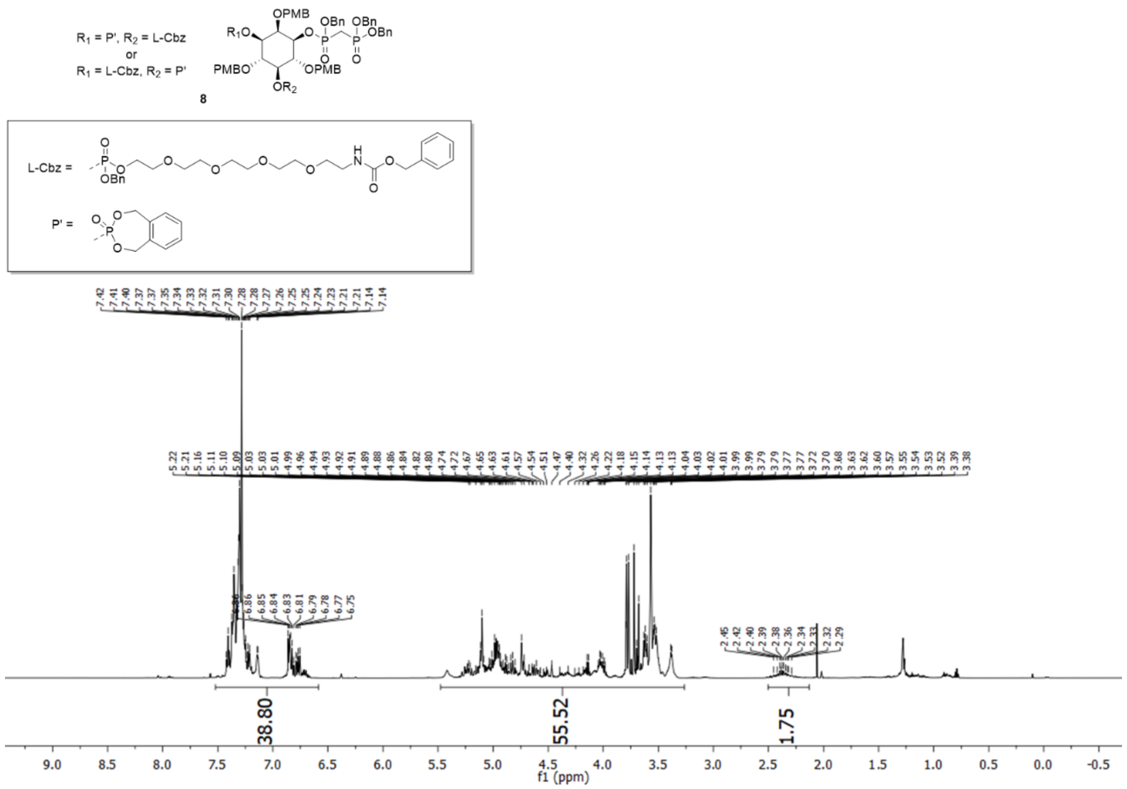


[1]H spectrum of compound **8**.

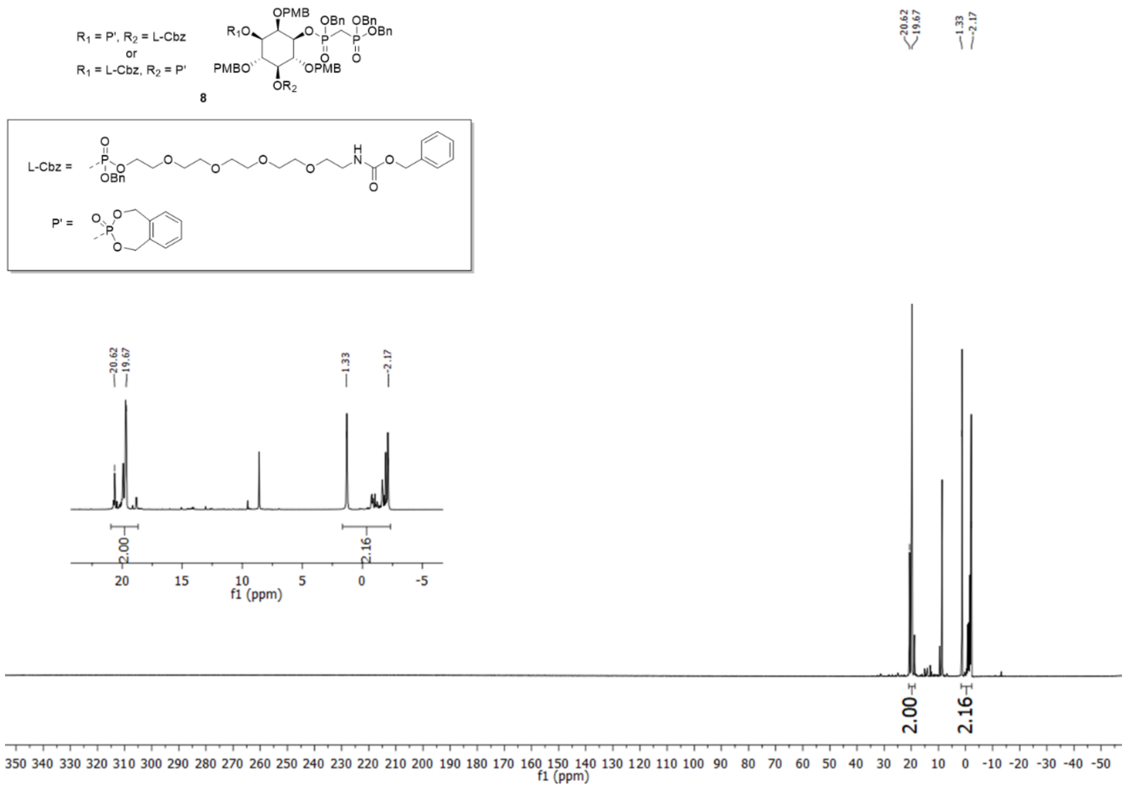


[31]P spectrum of compound **8**.

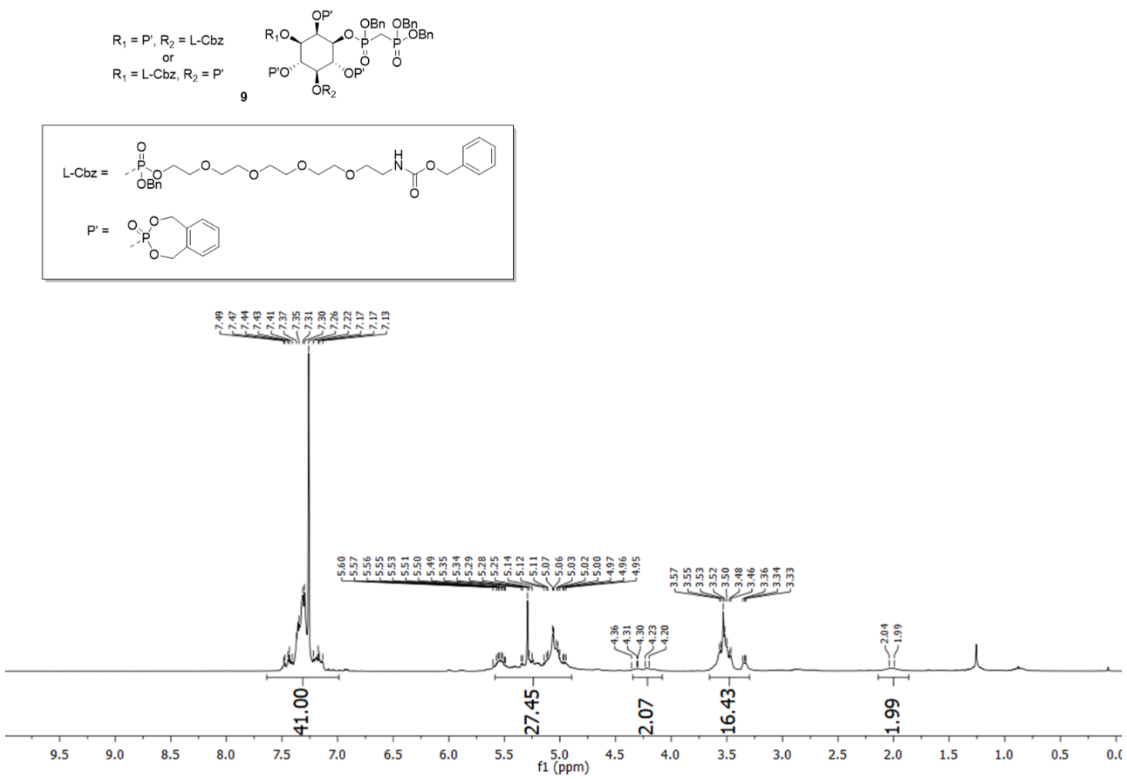

$^{1}$H spectrum of compound **9**.

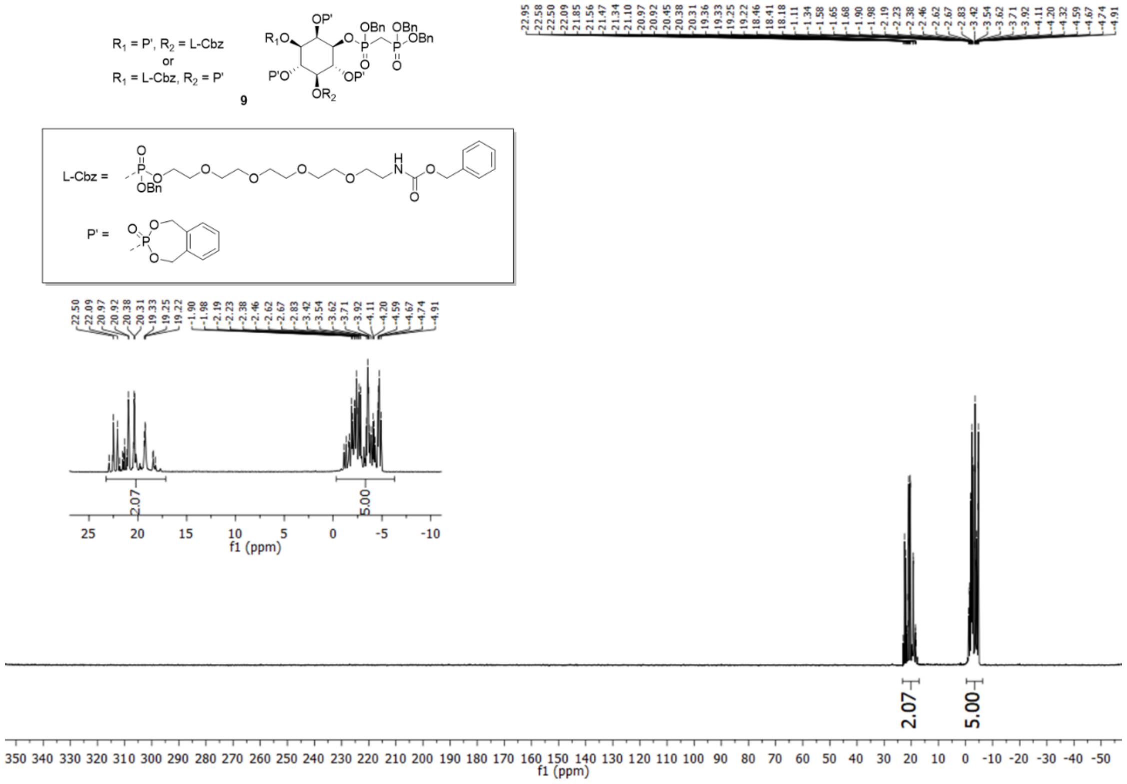

$^{31}$P spectrum of compound **9**.

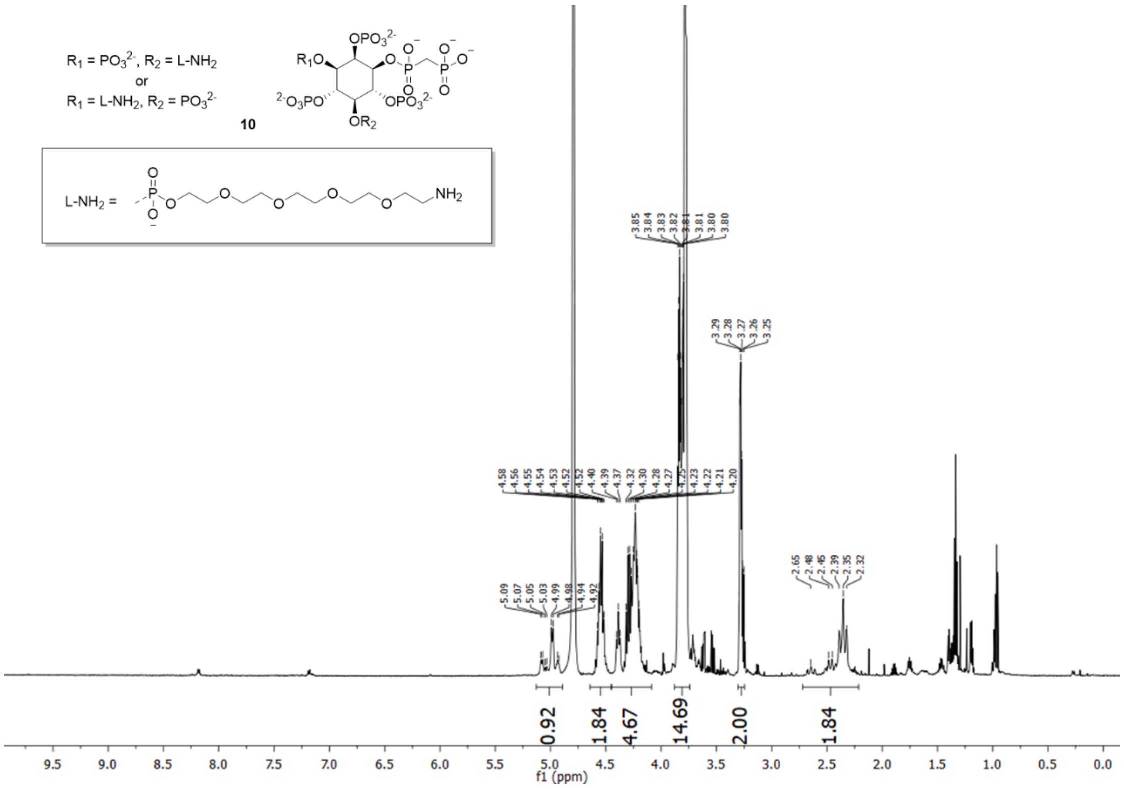

[1]H spectrum of compound **10**.

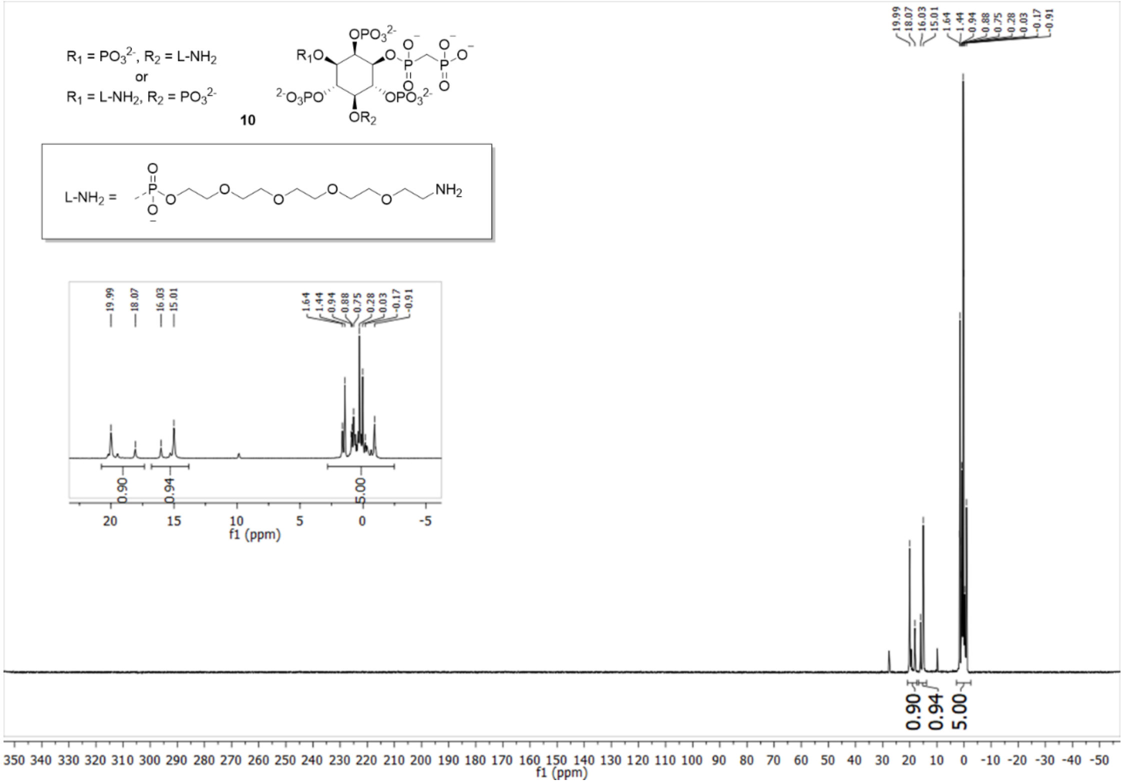

[31]P spectrum of compound **10**.

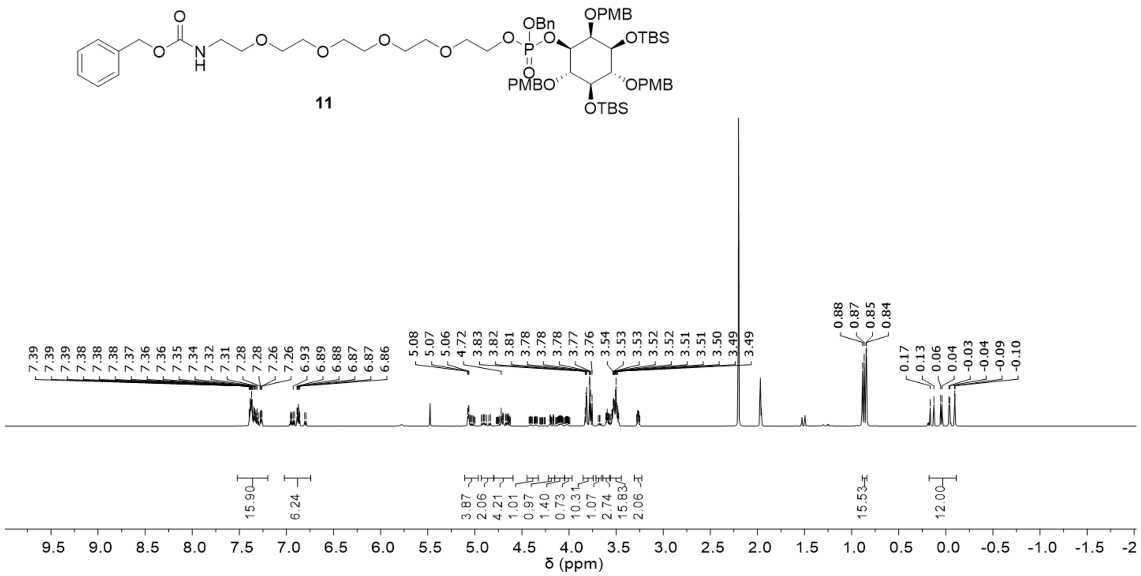

$^{1}H$ spectrum of compound **11**.

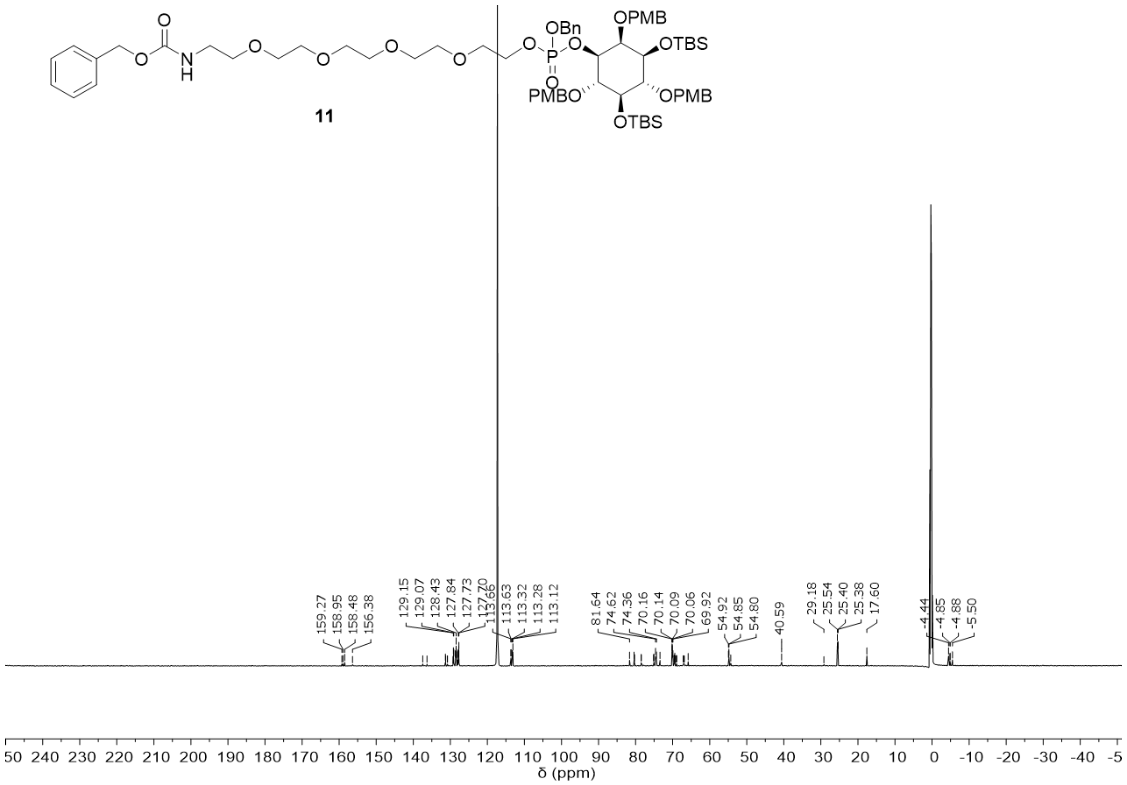

$^{13}C$ spectrum of compound **11**.

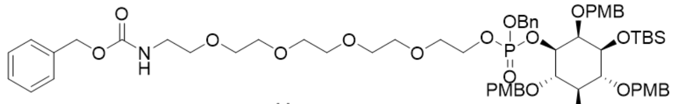


**11**

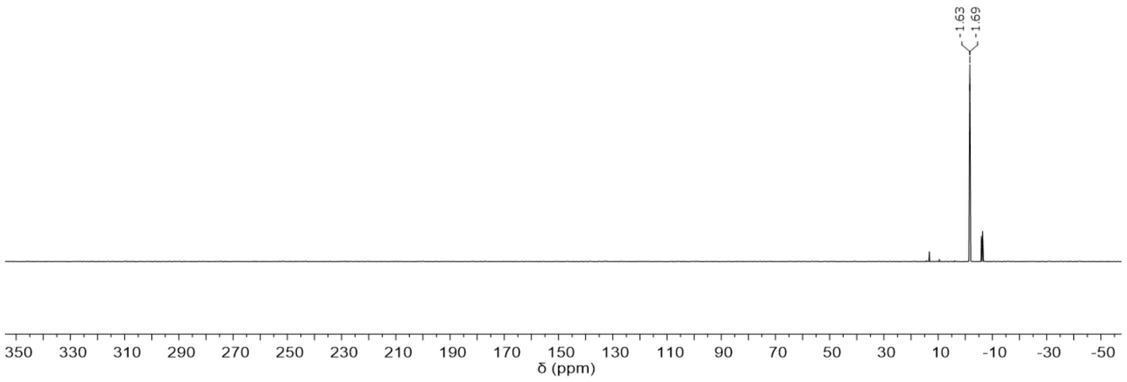


[31]P spectrum of compound **11**.

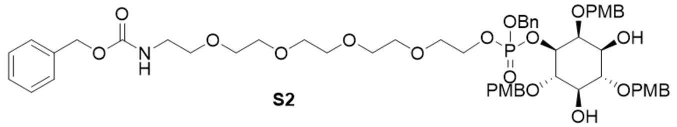


**S2**

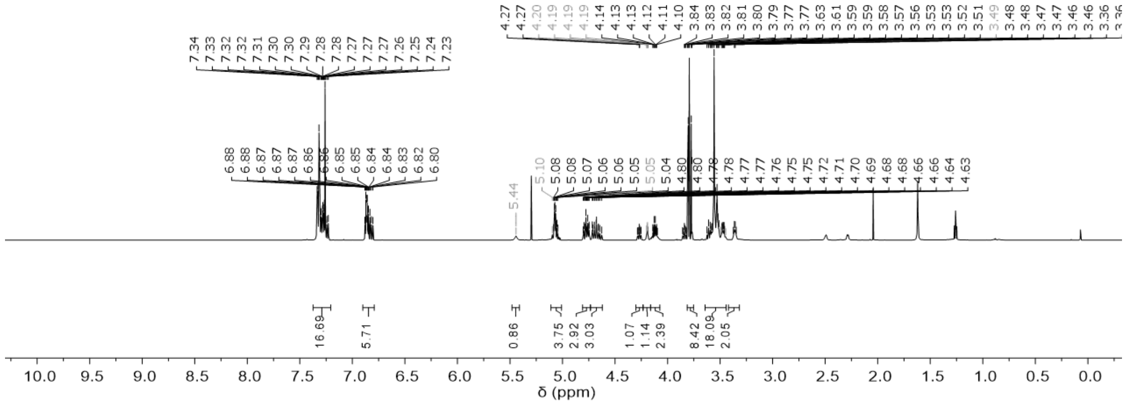


[1]H spectrum of compound **S2**.

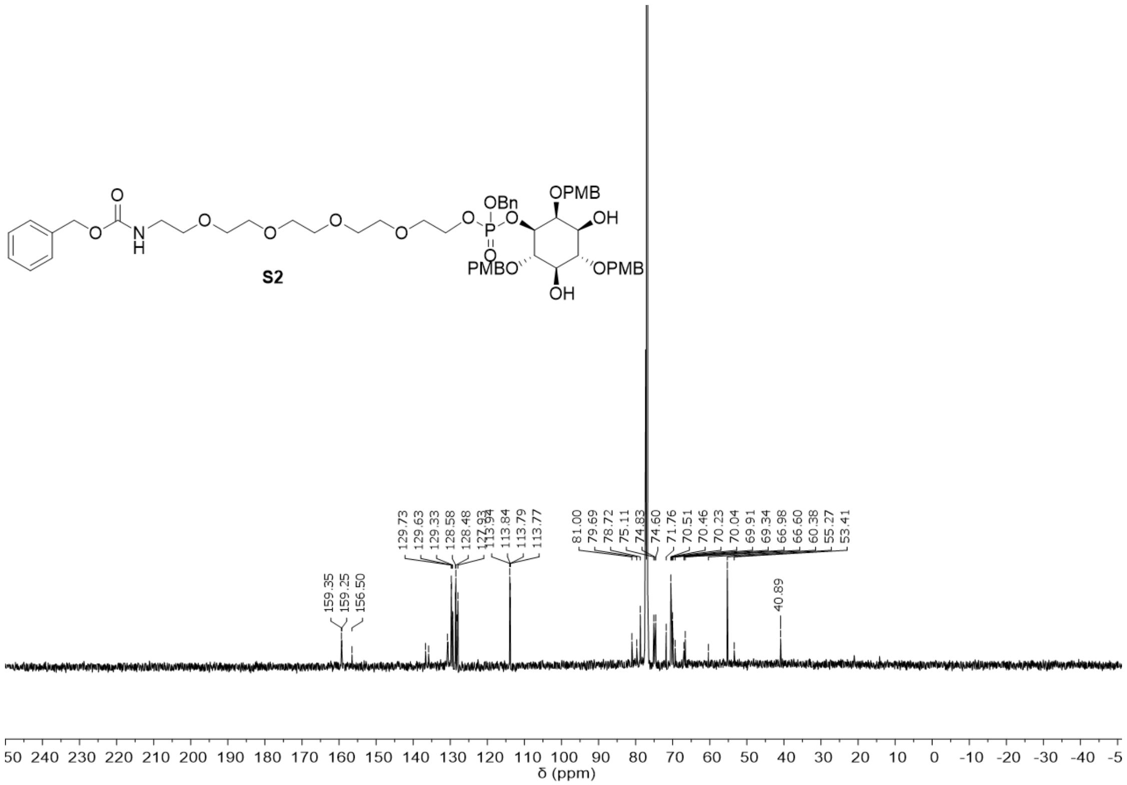


[13]C spectrum of compound **S2**.

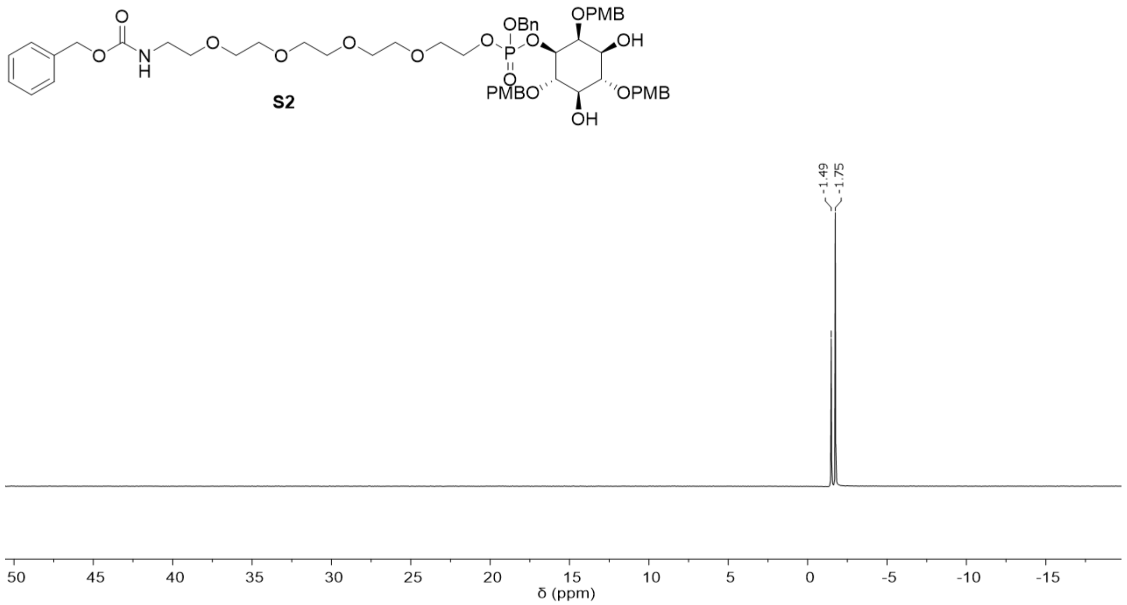


[31]P spectrum of compound **S2**.

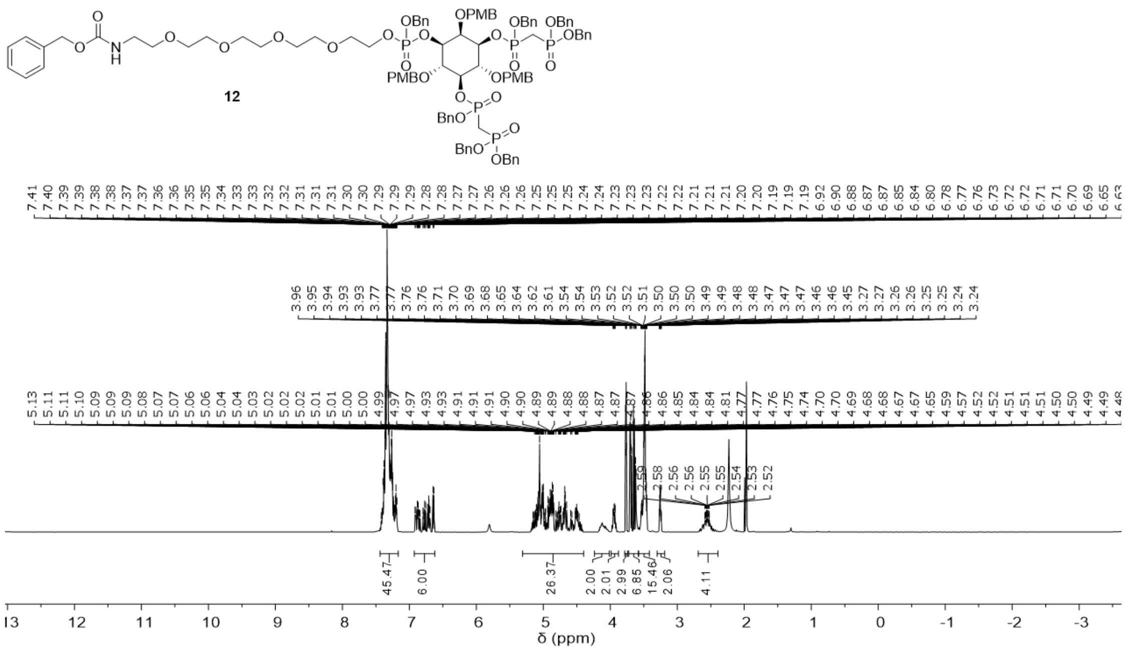

$^{1}$H spectrum of compound **12**.

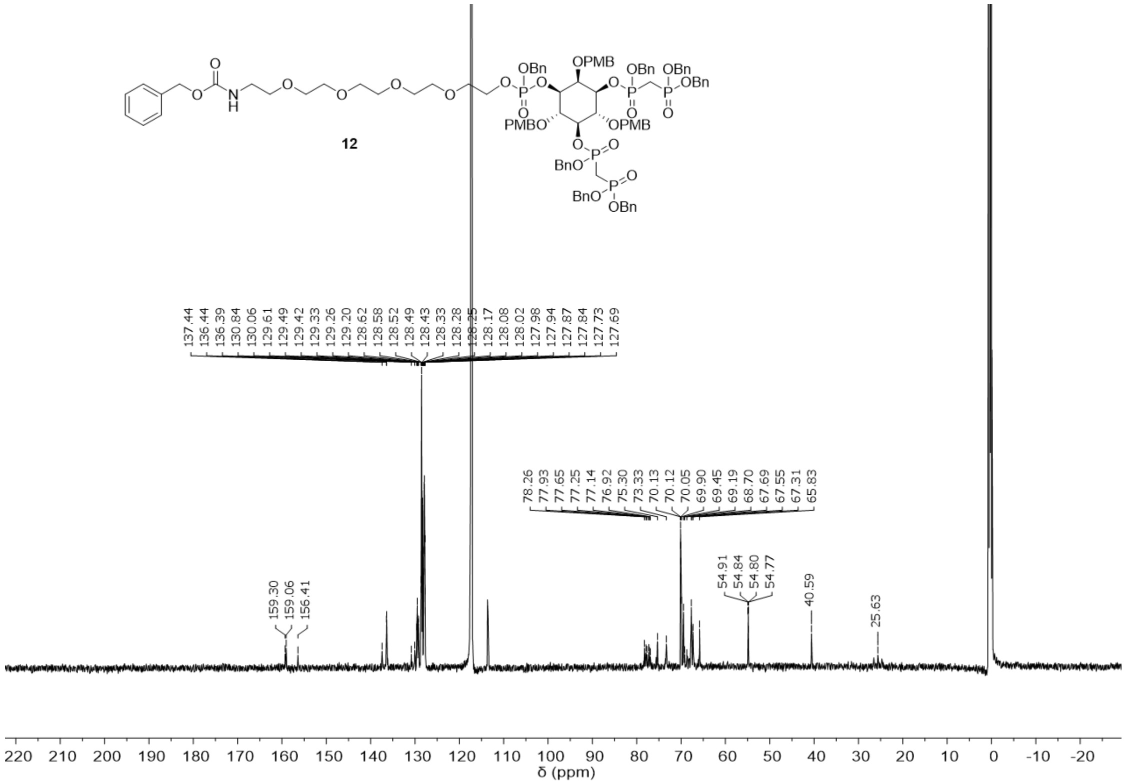

$^{13}$C spectrum of compound **12**.

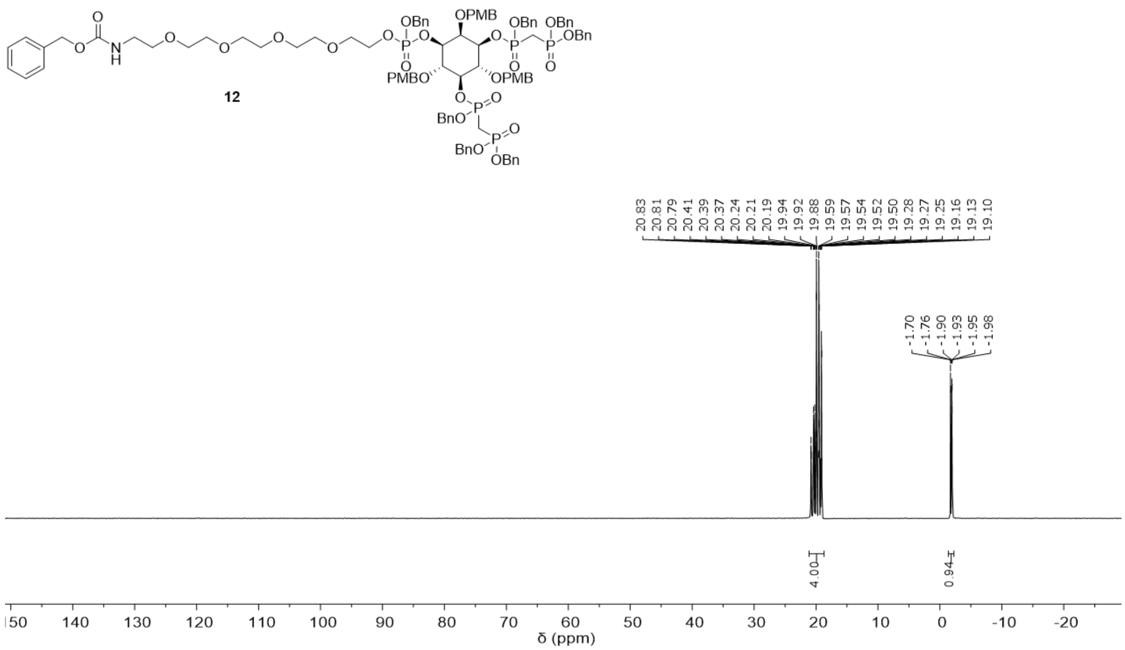


$^{31}$P spectrum of compound **12**.

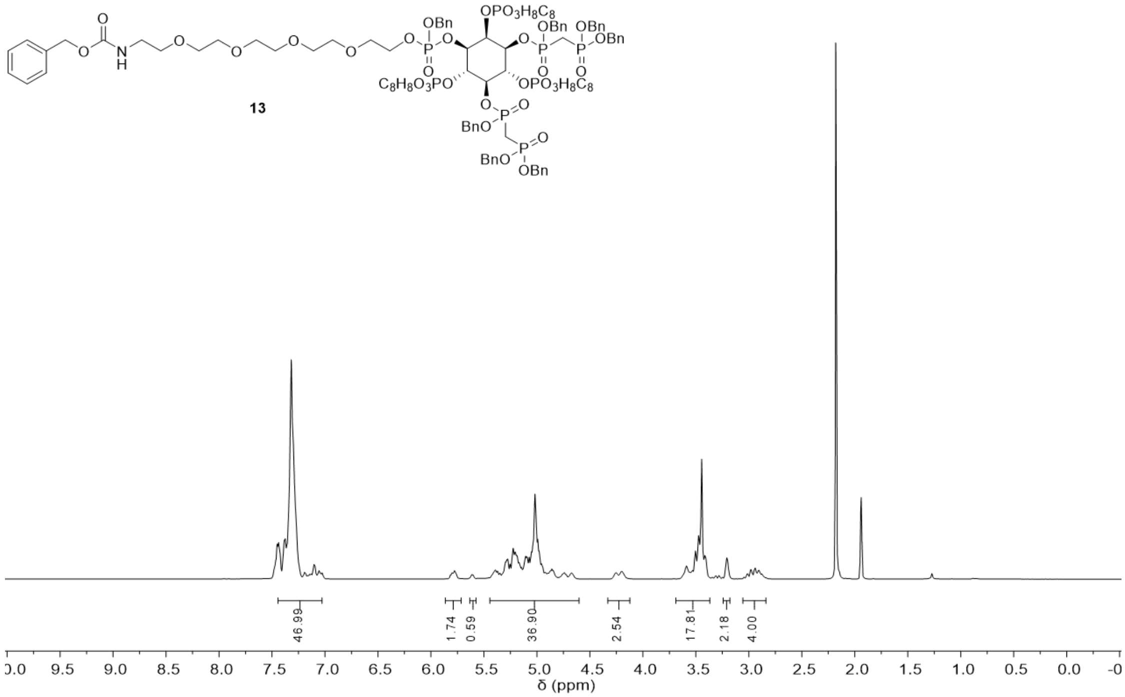


$^{1}$H spectrum of compound **13**.

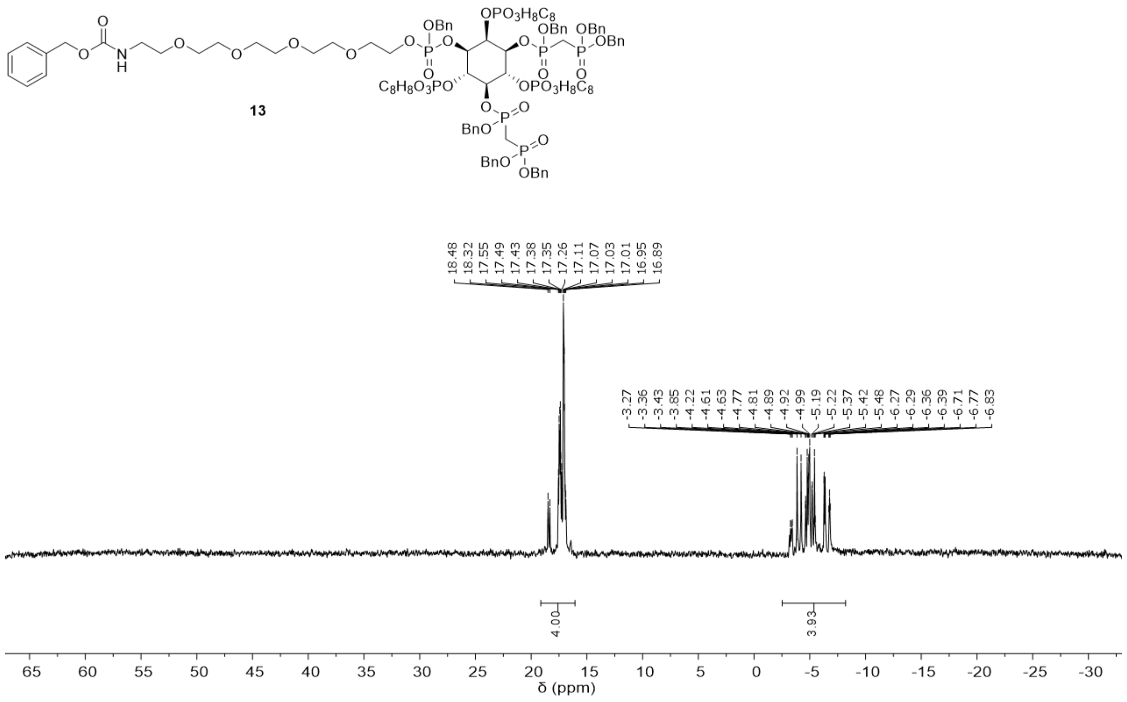

$^{31}P$ spectrum of compound **13**.

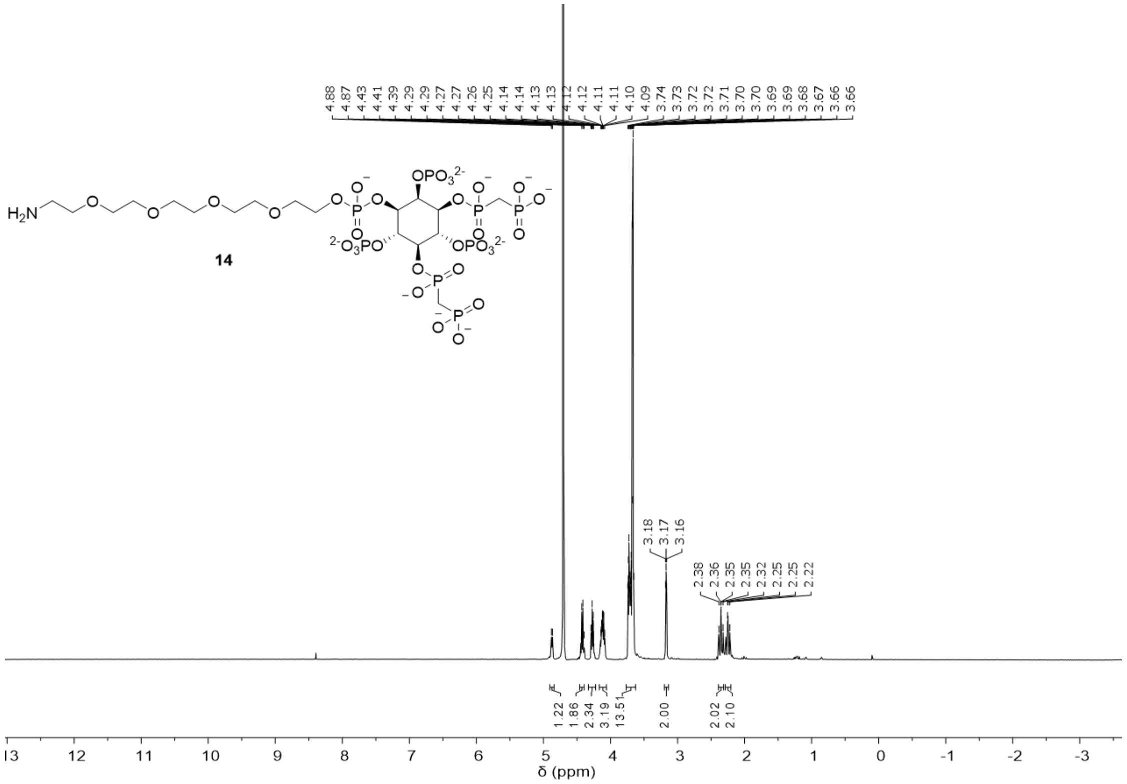

$^{1}H$ spectrum of compound **14**.

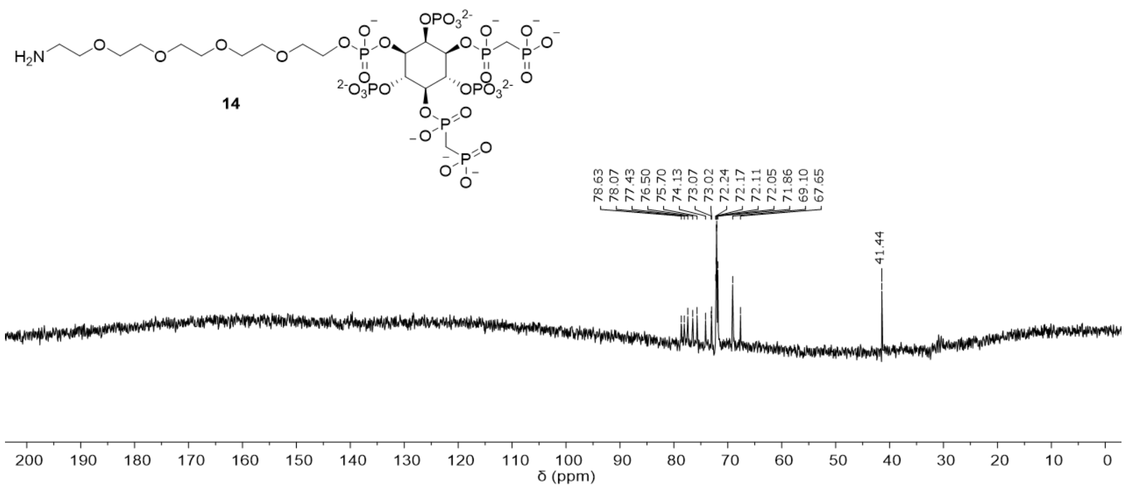


$^{13}C$ spectrum of compound **14**.

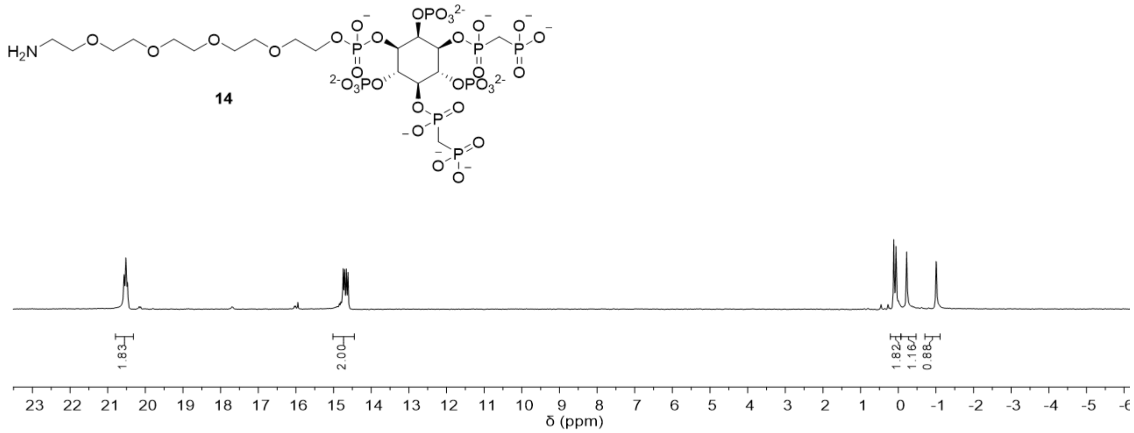


$^{31}P$ spectrum of compound **14**.

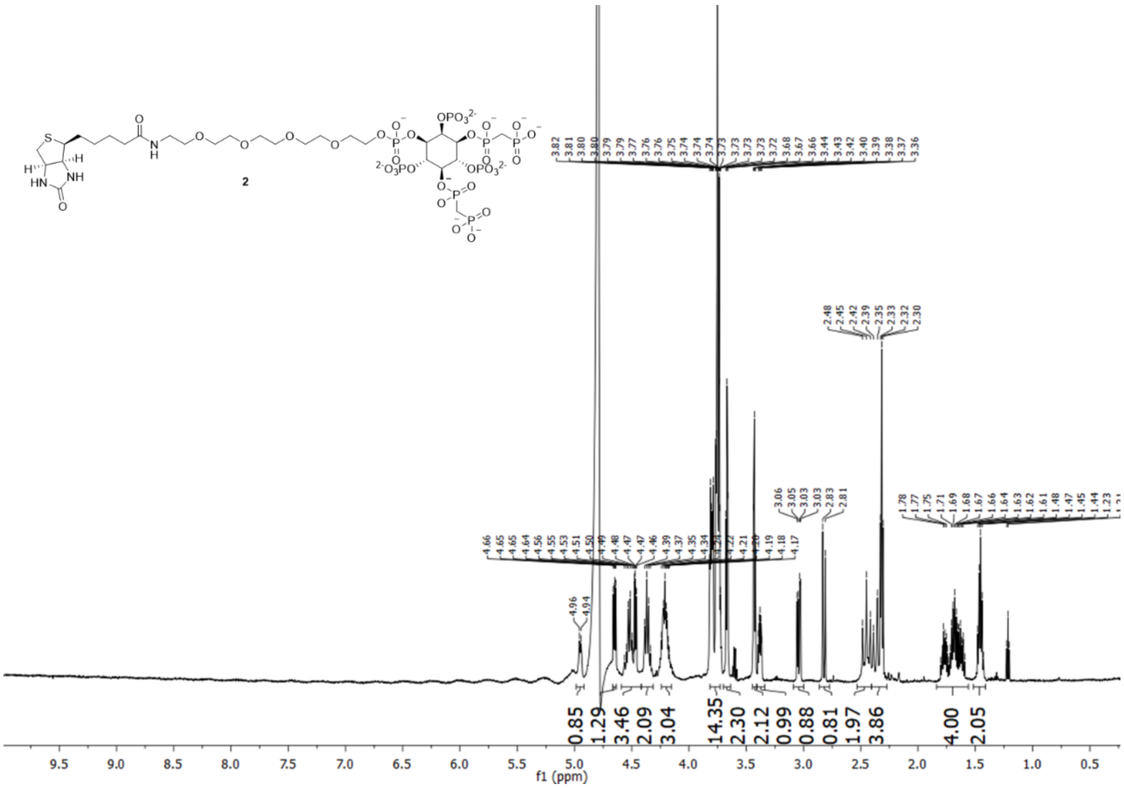


[1]H spectrum of compound **2**.

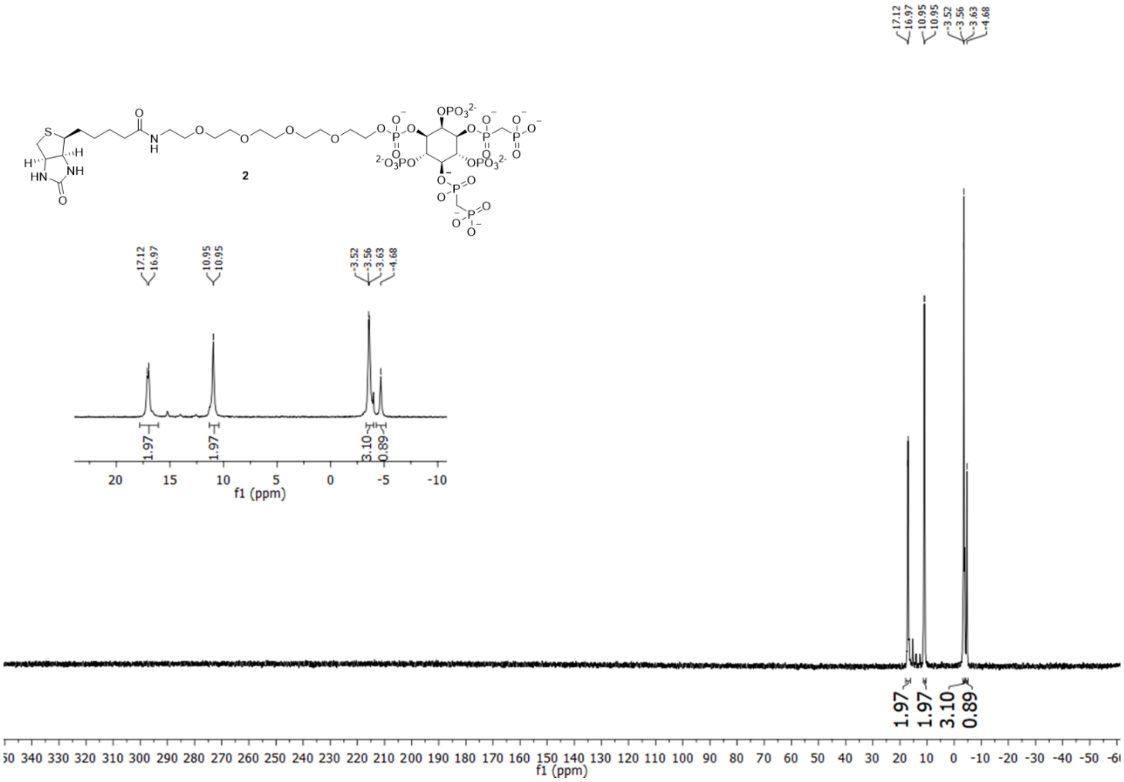


[31]P spectrum of compound **2**.

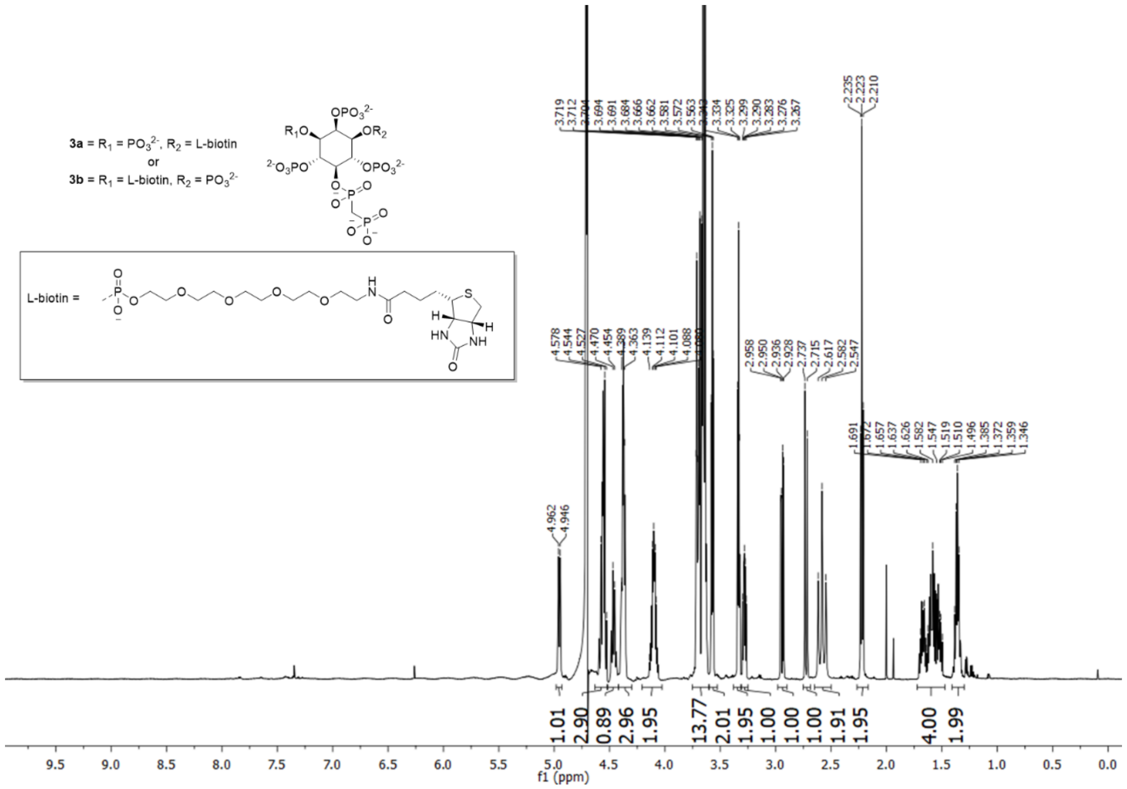


[1]H spectrum of compound **3**.

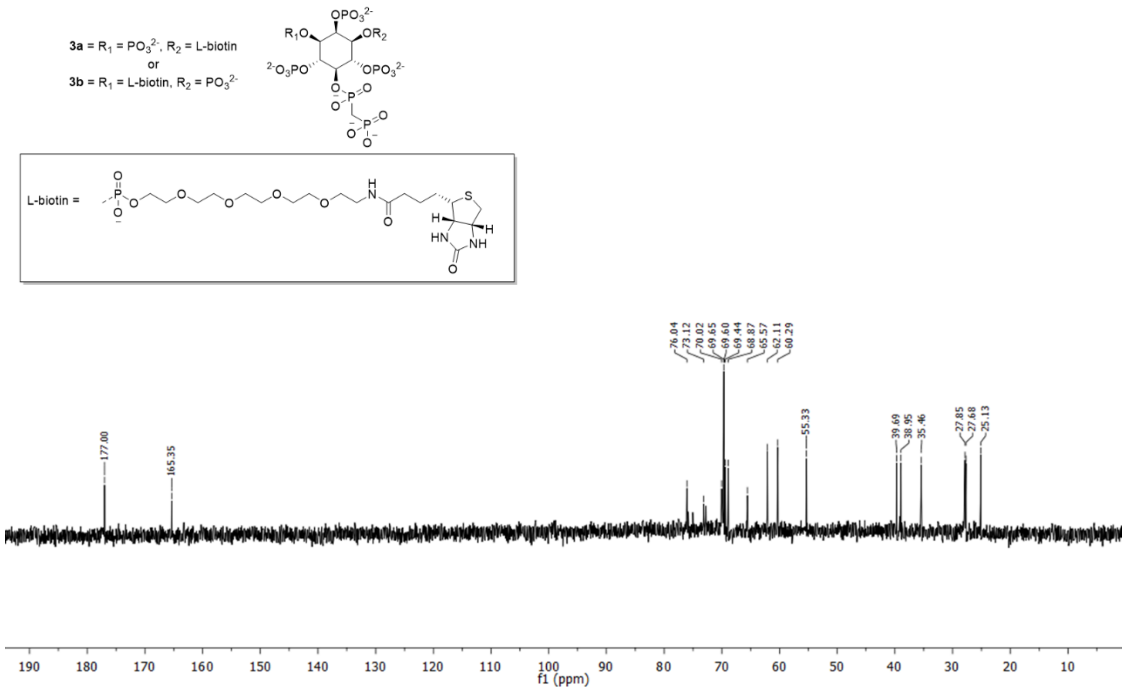


[13]C spectrum of compound **3**.

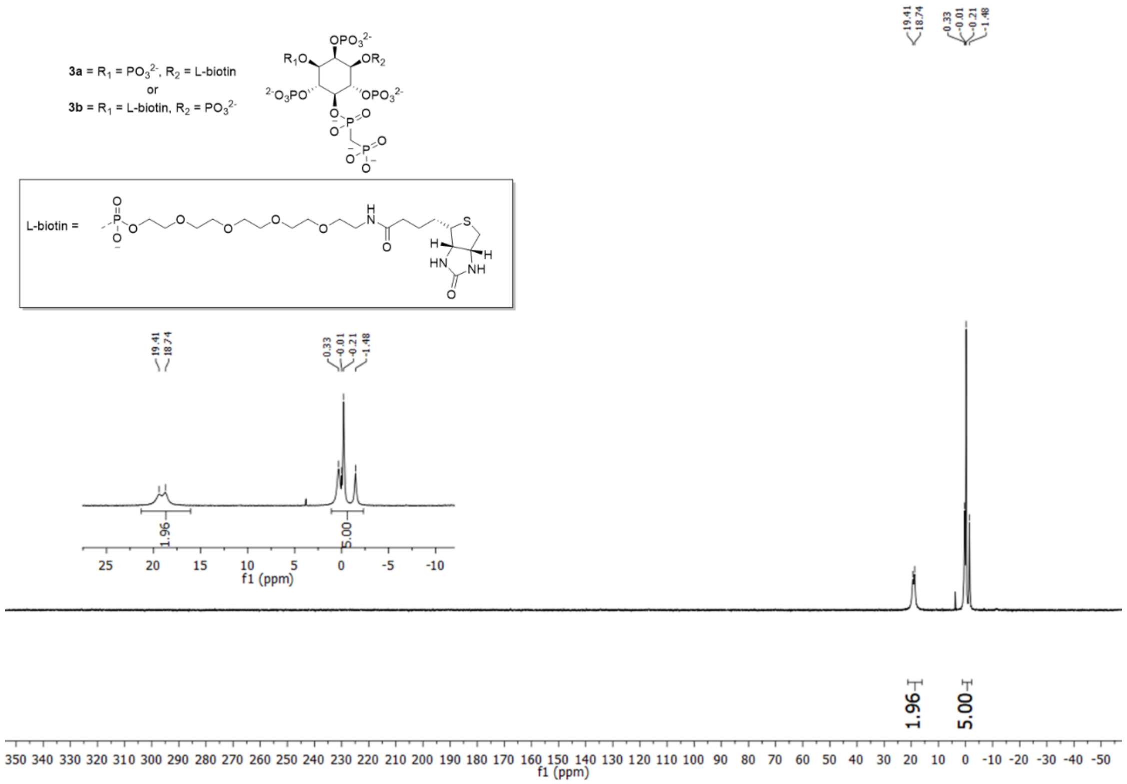


$^{31}$P spectrum of compound **3**.

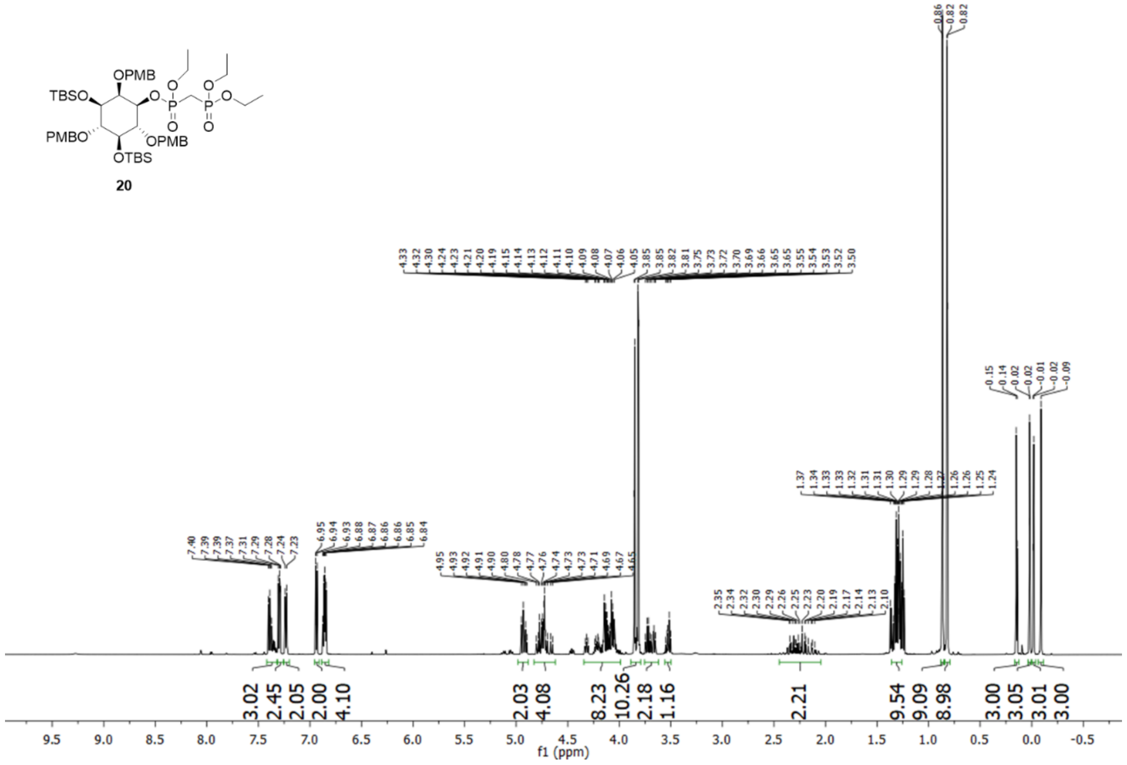


$^{1}$H spectrum of compound **20**.

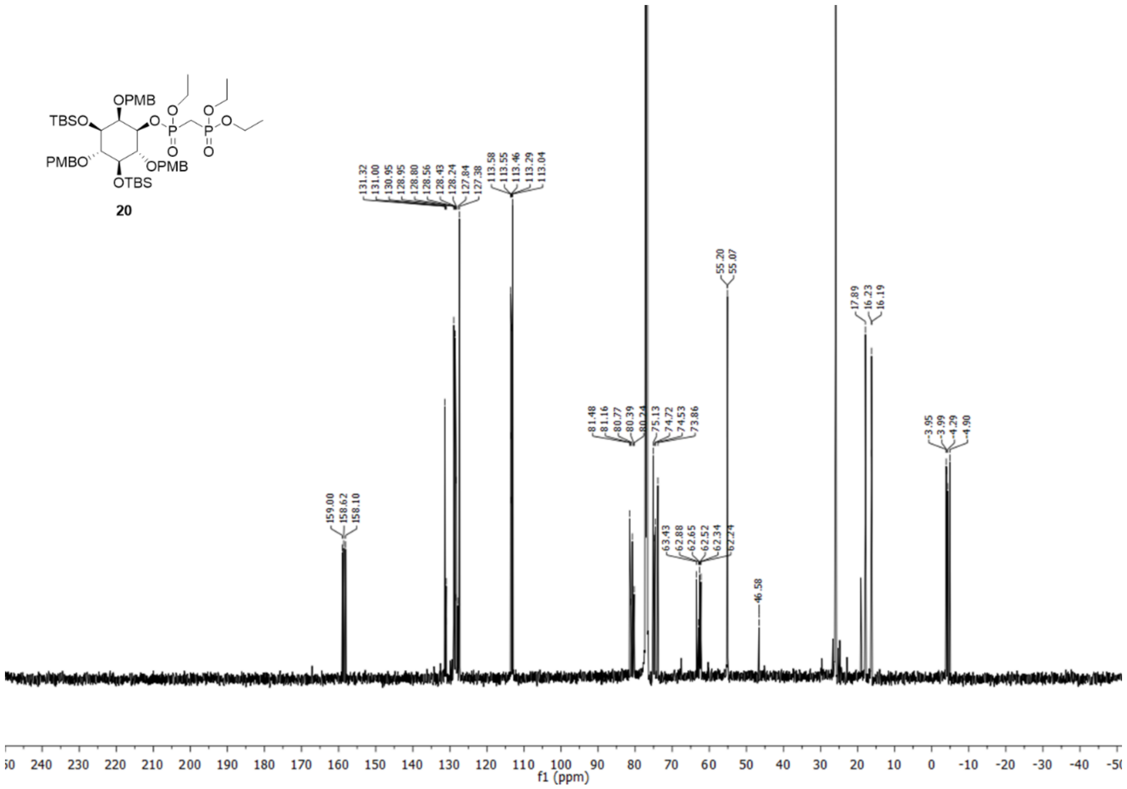

[13]C spectrum of compound **20**.

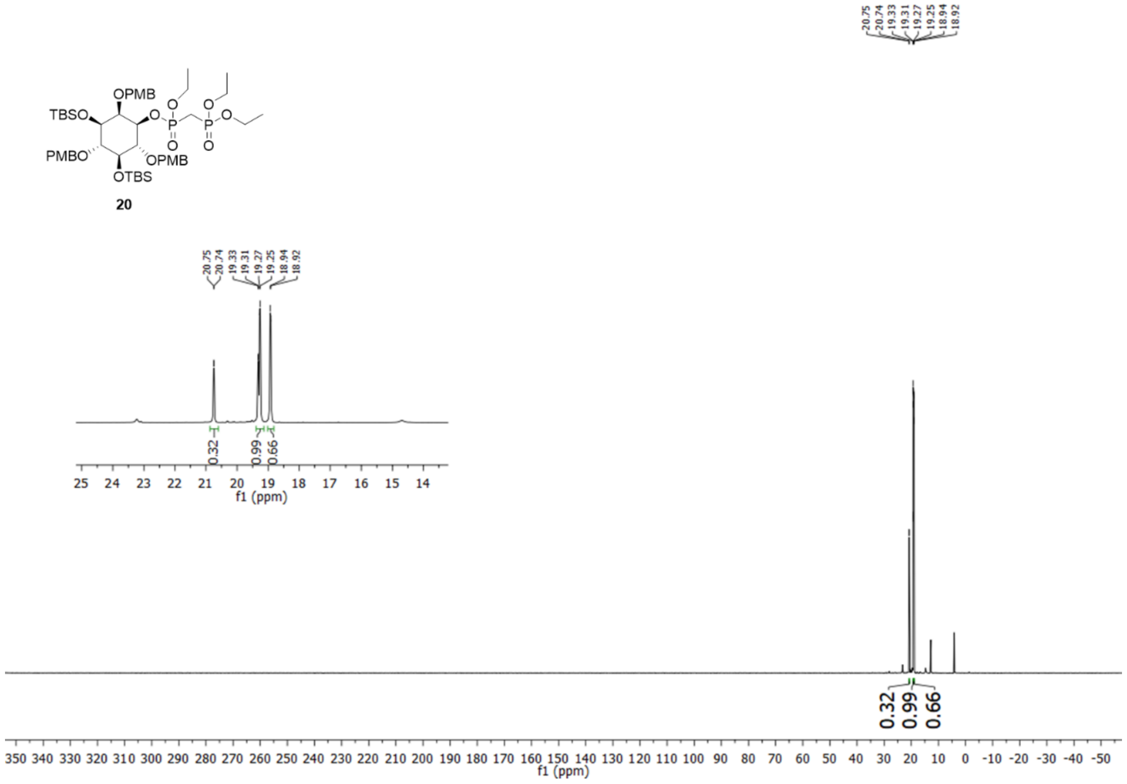

[31]P spectrum of compound **20**.

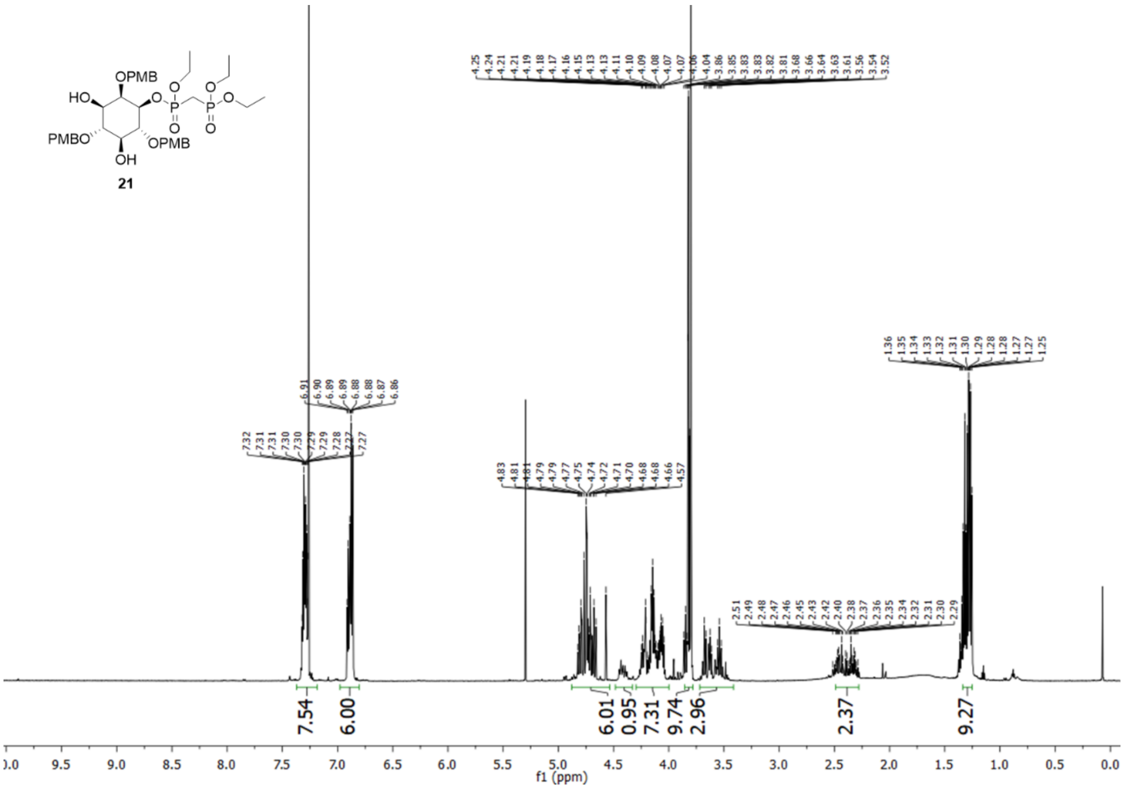

[1]H spectrum of compound **21**.

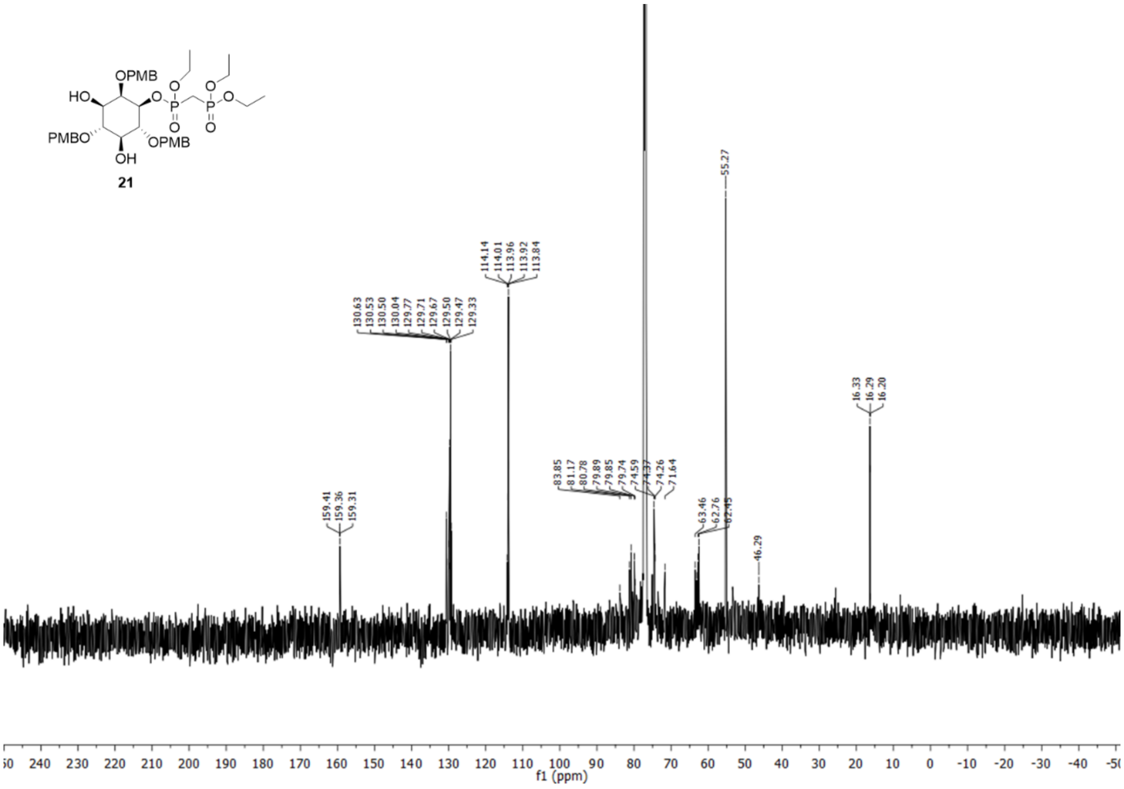

[13]C spectrum of compound **21**.

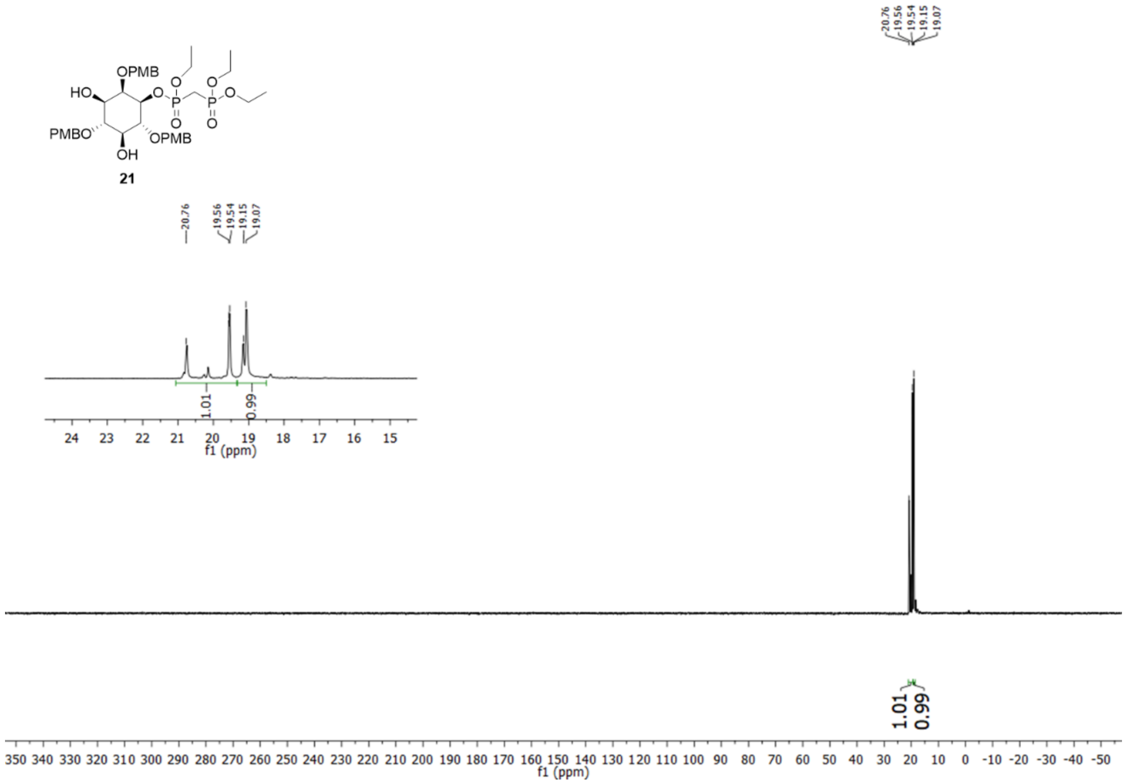


$^{31}$P spectrum of compound **21**.

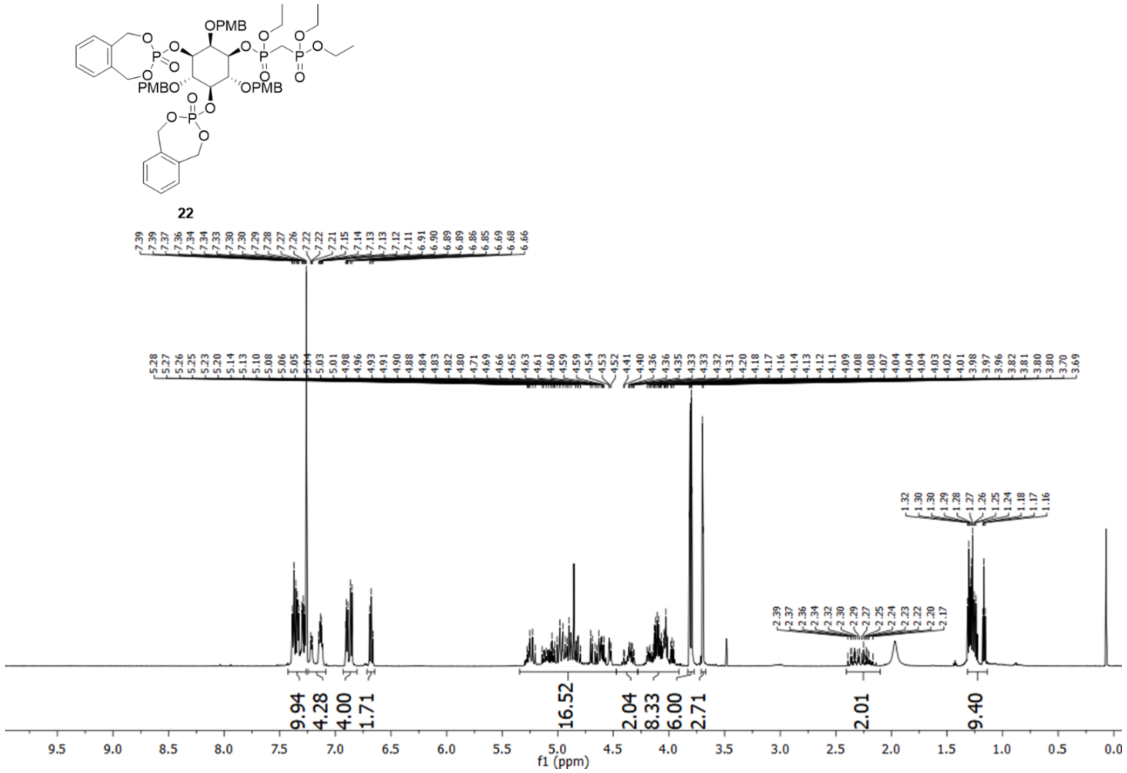


$^{1}$H spectrum of compound **22**.

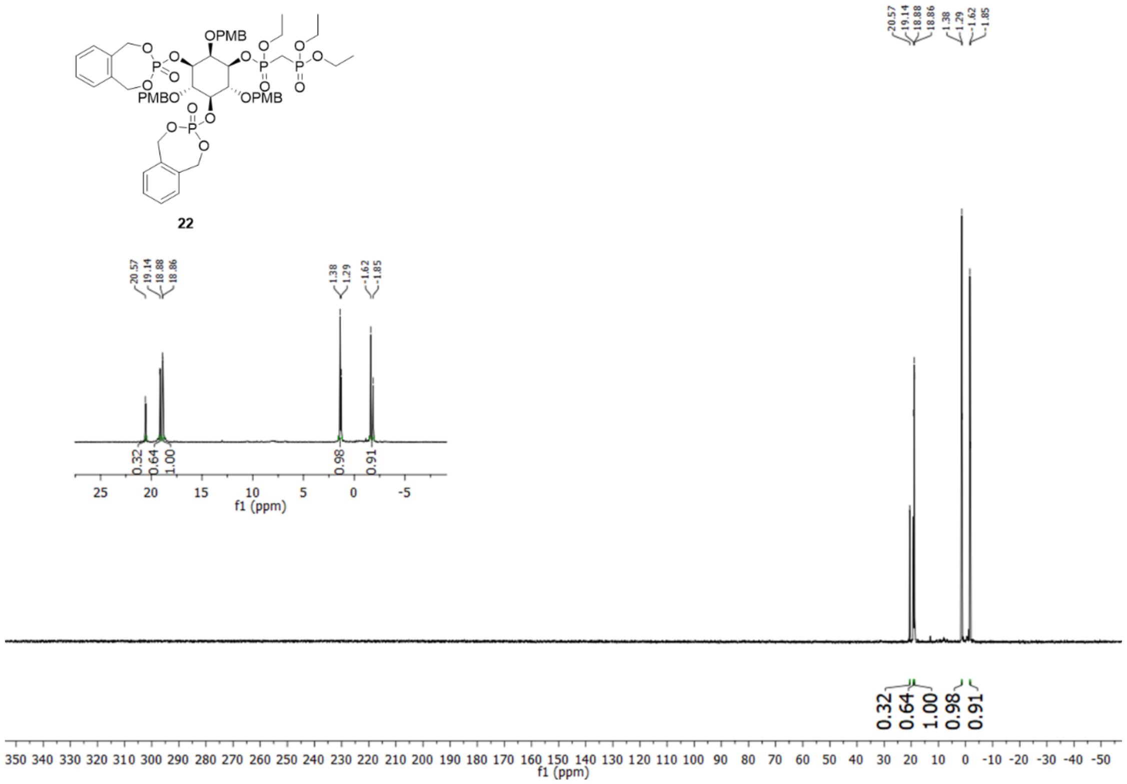

[31]P spectrum of compound **22**.

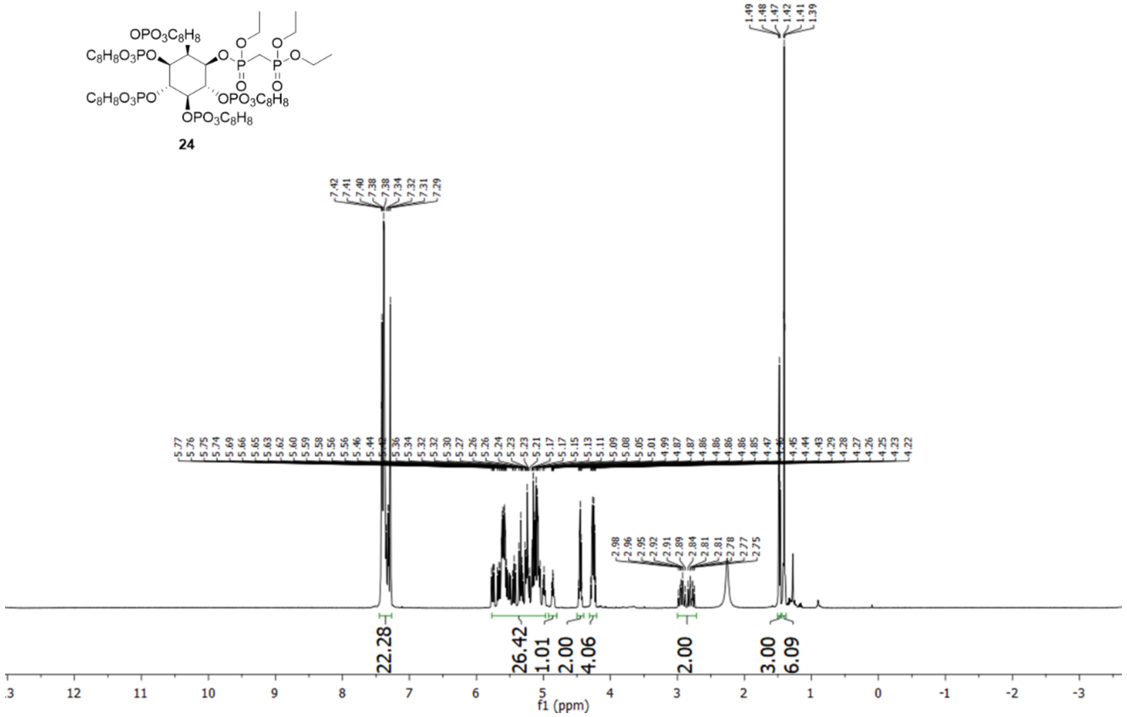

[1]H spectrum of compound **24**.

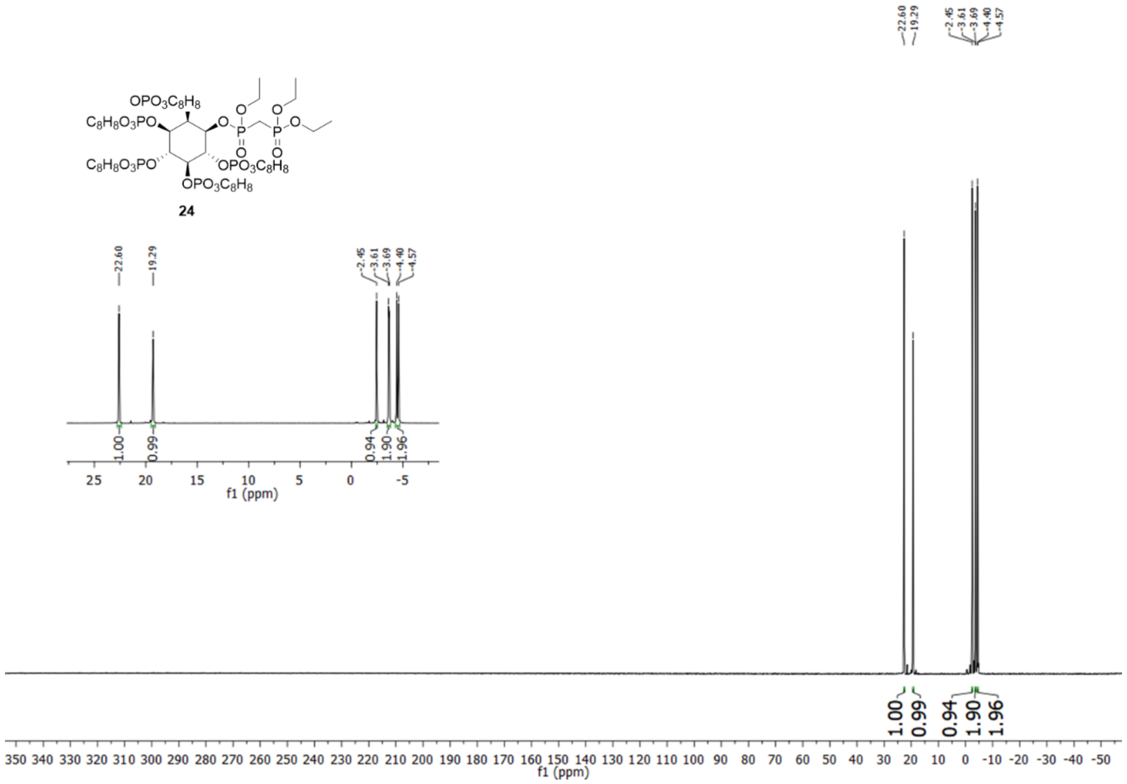


$^{31}$P spectrum of compound **24**.
